# Multi-resolution deconvolution of spatial transcriptomics data reveals continuous patterns of inflammation

**DOI:** 10.1101/2021.05.10.443517

**Authors:** Romain Lopez, Baoguo Li, Hadas Keren-Shaul, Pierre Boyeau, Merav Kedmi, David Pilzer, Adam Jelinski, Eyal David, Allon Wagner, Yoseph Addad, Michael I. Jordan, Ido Amit, Nir Yosef

## Abstract

The function of mammalian cells is largely influenced by their tissue microenvironment. Advances in spatial transcriptomics open the way for studying these important determinants of cellular function by enabling a transcriptome-wide evaluation of gene expression *in situ*. A critical limitation of the current technologies, however, is that their resolution is limited to niches (spots) of sizes well beyond that of a single cell, thus providing measurements for cell aggregates which may mask critical interactions between neighboring cells of different types. While joint analysis with single-cell RNA-sequencing (scRNA-seq) can be leveraged to alleviate this problem, current analyses are limited to a discrete view of cell type proportion inside every spot. This limitation becomes critical in the common case where, even within a cell type, there is a continuum of cell states that cannot be clearly demarcated but reflects important differences in the way cells function and interact with their surroundings. To address this, we developed Deconvolution of Spatial Transcriptomics profiles using Variational Inference (DestVI), a probabilistic method for multi-resolution analysis for spatial transcriptomics that explicitly models continuous variation within cell types. Using simulations, we demonstrate that DestVI is capable of providing higher resolution compared to the existing methods and that it can estimate gene expression by every cell type inside every spot. We then introduce an automated pipeline that uses DestVI for analysis of single tissue slices and comparison between tissues. We apply this pipeline to study the immune crosstalk within lymph nodes to infection and explore the spatial organization of a mouse tumor model. In both cases, we demonstrate that DestVI can provide a high resolution and accurate spatial characterization of the cellular organization of these tissues, and that it is capable of identifying important cell-type-specific changes in gene expression - between different tissue regions or between conditions. DestVI is available as an open-source software package in the scvi-tools codebase (https://scvi-tools.org).

## Introduction

Spatial transcriptomics opens up new opportunities to define the organization of cellular niches and crosstalk that modulate cellular function [1]. In particular, this emerging technology helped study the organization of complex tissues such as the mouse brain [2] and the human heart [3]. The research of human pathologies, such as the structure of tumors, is also an important avenue for spatial transcriptomics [4,5] since the tumor microenvironment consists of a rich milieu of cell types and states that are organized in different anatomical niches.

The landscape of experimental assays for performing spatial transcriptomics analyses of tissue sections is diverse, although all assays are data-rich and require automated and quantitative computational analyses. For example, methods based on fluorescence imaging (MERFISH [6], osmFISH [2,6], seqFISH [7]) have near single-transcript resolution. However, these methods rely on cell-segmentation algorithms [8,9]. Additionally, these studies are dependent on pre-selected marker genes and are not genome-wide and hence require imputation of the missing gene to avoid overlooking critical information [10–12]. On the other hand, pseudo-bulk spatial transcriptomic measurements (Slide-Seq [13,14], 10x Visium [15]) are appealing technologies as they provide measurements of the whole transcriptome, although the spatial resolution, in current versions, is limited to cell aggregates (10 microns for Slide-Seq and 55 microns for 10x Visium). Depending on the density of the tissue, a single bead spot of 10x Visium may have a large number of cells, emphasizing the need for deconvolution of spots to obtain a better resolution view of their cellular content.

To overcome this limitation of current leading genome-wide spatial transcriptomics experimental protocols, these datasets are often matched with a single cell RNA-sequencing (scRNA-seq) dataset from the same tissue. The convention for analyzing such pairs of datasets (as implemented by all existing pipelines, including NMFReg [14], RCTD [16], SPOTLight [17], Stereoscope [18], DSTG [19], and cell2location [20]) is to apply a two-step process. First, a dictionary of cell types is inferred from the scRNA-seq data; then, the proportion of each cell type within each spot is estimated using a linear model. This approach has had promising results, in particular when analyzing brain tissue sections in which the diversity of cellular composition is well captured by a discrete view of cell types [21].

However, the aforementioned methods are more challenging to apply in settings where there is no clear way to stratify cells into discrete types or subtypes. This is especially important when cells that belong to the same overall type (e.g., T helper cells) may carry different functions and span a continuum of states (e.g., following different inflammatory signals) [22]. As a way to resolve this fundamental conundrum of single-cell data analysis, current algorithms leave the user with the choice of setting the granularity in which the data is to be analyzed (i.e., number of clusters per broad cell type). However, there are some inherent trade-offs: deeper clustering of the scRNA-seq data provides more granular transcriptomic resolution but makes the deconvolution problem more difficult, and the results potentially less accurate.

In this manuscript, we propose a conceptually different framework. Instead of limiting the analysis to a discrete view of cell types, we propose to also model the variation within each cell type via continuous latent variables. Towards this end, we introduce DEconvolution of Spatial Transcriptomics profiles using Variational Inference (DestVI), a Bayesian model for multi-resolution deconvolution of cell types in spatial transcriptomics data. Much like existing deconvolution methods, DestVI takes as input a scRNA-seq dataset, with annotations and a spatial transcriptomics dataset. Unlike other methods, DestVI learns cell-type-specific profiles and continuous sub-cell-type variations using a conditional deep generative model [23] and recovers the cell-type frequency as well as a cell-type-specific snapshot of the average transcriptional state at every spot. We also propose a post-hoc analysis pipeline, based on auto-correlation and a spatially-aware version of PCA, to highlight the main axes of spatial variation and help guide the downstream analysis. Our pipeline also helps extract molecular signatures that characterize a given tissue slice or different areas inside the same tissue using cell-type-specific differential expression.

We used simulations to benchmark DestVI against discrete deconvolution approaches applied to different levels of cell state granularity (i.e., number of clusters per cell type) and show that DestVI significantly outperforms every baseline in terms of imputation of cell-type-specific gene expression. We then showcase the broad usability of DestVI in two very different biological models using 10x Visium measurements. First, we applied DestVI to study the murine lymph node, which is a considerably structured and well-studied secondary lymphoid organ. We report the spatial organization of cell types in this organ at steady-state and study the effects of stimulation with pathogens [24]. DestVI accurately identifies the effects on the spatial and transcriptional organization of monocytes that are activated upon immunization and form immune response niches. We then proceeded to apply DestVI to a mouse tumor model. In this more complex tissue, DestVI delineates the spatial coordinates of main immune cells within the tumor microenvironment (TME). Furthermore, we mapped the sub-populations of macrophages onto the tumor and recovered spatial patterns of hypoxia activation within the tumor core [25]. DestVI is implemented in the scvi-tools package [26], and readily available along with accompanying tutorials.

## Results

### Multi-resolution deconvolution of cell states in spatial transcriptomics data using DestVI

DestVI uses two different latent variable models (LVMs) [27] for delineating cell-type proportions as well as cell-type-specific continuous sub-states. The input for DestVI is a pair of transcriptomics datasets: a query spatial transcriptomics data as well as a reference scRNA-seq data from the same tissue (**Figure 1**). DestVI assumes that each cell in the reference dataset is annotated with a discrete cell-type label (**Online Methods**). The output of DestVI consists of two components: first, the expected proportion of cell types for every spot, and second, a continuous estimation of cell-state for every cell type in every spot, which represents an average state for cells of this type in the spot (**Figure 1A**). This spot-level information may then be used for downstream analysis and formulation of biological hypotheses (described later in this section).

**Figure 1:**
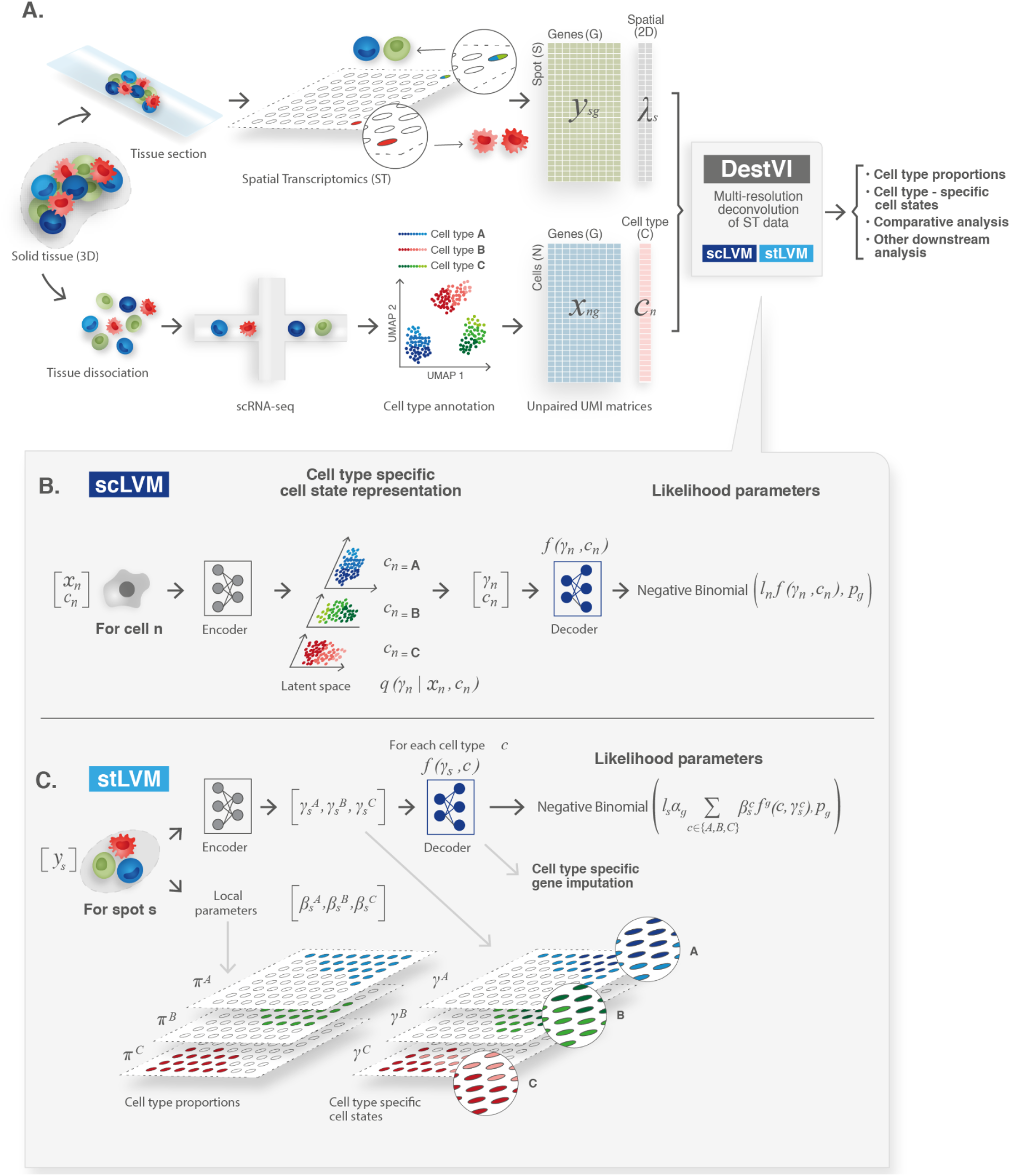
Schematic representation of the spatial transcriptomics analysis pipeline with DestVI. **(A)** A spatial transcriptomics analysis workflow relies on two data modalities, producing unpaired transcriptomic measurements, in the form of count matrices. The spatial transcriptomics (ST) data measures the gene expression *y*_*s*_ in a given spot *s*, and its location λ_*s*_. However, each spot may contain multiple cells. The single cell RNA-sequencing data measures the gene expression *x*_*n*_ in a cell *n*, but the spatial information is lost because of tissue dissociation. After annotation, we may associate each cell with a cell type *c*_*n*_. These matrices are the input to DestVI, composed of two latent variable models: the single-cell latent variable model (scLVM) and the spatial transcriptomics latent variable model (stLVM). DestVI outputs a joint representation of the single-cell data, and the spatial data by estimating the proportion of every cell type in every spot, and projecting the expression of each spot onto cell-type-specific latent spaces. These inferred values may be used for performing downstream analysis such as cell-type-specific differential expression and comparative analyses of conditions. **(B)** Schematic of the scLVM. RNA counts and cell type information from the single cell RNA-sequencing data are jointly transformed by an encoder neural network into the parameters of the approximate posterior of γ_*n*_, a low-dimensional representation of cell-type-specific cell state. Next, a decoder neural network maps samples from the approximate posterior of γ_*n*_ along with the cell type information *c*_*n*_ to the parameters of a negative binomial distribution for every gene. Note that we use the superscript notation *f*^*g*^ to denote the *g*-th output of the network, that is the *g*-th entry ρ_*ng*_ of the vector ρ_*n*_. **(C)** Schematic of the stLVM. RNA counts from the spatial transcriptomics data are transformed by an encoder neural network into the parameters of the cell-type-specific embeddings 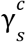. Free parameters 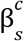 encode the abundance of cell type *c* in spot *s*, and may be normalized into cell-type proportions 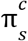 (**Online Methods**). Next, the decoder from the scLVM model maps cell-type-specific embeddings 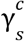 to estimates of cell-type-specific gene expression. These parameters are averaged across all cell types, weighted by the abundance parameters 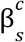, to approximate the gene expression of the spot with a negative binomial distribution. After training, the decoder may be used to perform cell-type-specific imputation of gene expression across all spots.

To model the reference scRNA-seq data, the first LVM (scLVM; **Figure 1B**) of DestVI posits that for each gene *g* and cell *n* the number of observed transcript, *x*_*ng*_, follows a negative binomial distribution, which has been shown to represent the properties of RNA count data [28]. The distribution is parameterized as (*r*_*ng*_, *p*_*g*_), with mean 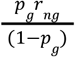 and where *p*_*g*_ is a gene specific parameter determining the mean-variance relationship at every spot. Parameter *r*_*ng*_ = *l*_*n*_ρ_*ng*_ of the negative binomial depends on the type assigned to the cell (*c*_*n*_), its overall number of detected molecules (*l*_*n*_), and a low-dimensional latent vector γ_*n*_ (here 5 dimensions) which captures the variability within its respective cell type. A neural network *f* maps γ_*n*_ and *c*_*n*_ to the vector ρ_*n*_. The gene-specific parameter *p*_*g*_ is optimized using variational Bayesian inference (**Online Methods**). scLVM is closely related to scVI, our previous work in scRNA-seq modeling strategies [29], although here we aim at capturing transcriptional variation that is cell-type specific. To fit scLVM, DestVI relies on amortized variational inference with deep neural networks [30]. After this procedure, we obtain for every cell a distribution *q*_ϕ_ (γ_*n*_| *c*_*n*_, *x*_*n*_)that quantifies the cell state, as well as a measure of its uncertainty (**Online Methods**).

To model the spatial transcriptomics data, the second LVM (stLVM, **Figure 1C**) of DestVI posits that for each gene *g* and each spot *s* the number of observed transcripts *x*_*sg*_ also follows a negative binomial distribution (as in [18], [20]). The rate parameter of the distribution *r*_*sg*_ depends on latent factors that capture technical variation (*l*_*s*_ - the overall number of molecules detected in the spot and α_*g*_ - a multiplicative factor to correct for gene-specific bias between spatial and scRNA-seq measurements) and biological variation, decomposed over cell types. The latter factors come at two levels: 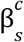 is a scalar proportional to the relative part of cells of type *c* inside the spot, and 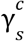 is a low-dimensional vector that estimates the average state of these cells. To ensure this correspondence, stLVM uses the same decoder neural network trained by scLVM - a step that can be interpreted as transfer learning of cell state decoding - from scRNA-seq data to the spatial data. To facilitate the decoupling between the factors that are included in the stLVM model and to further facilitate consistency with the scRNA-seq measurements, we utilize an empirical prior for 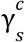. This prior is based on the values of γ^*c*^ inferred for the scRNA-seq data from cell type *c* [31]. To fit stLVM, DestVI relies on an amortized maximum a posteriori (MAP) inference scheme, in which the parameters for the cell-type proportions are kept as free, but the 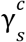 values are tied with a neural network (**Online Methods**). Finally, in our implementation, the actual cell-type proportions 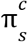 are obtained by normalizing 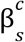 (to sum to one).

### DestVI outperforms competing methods for imputing cell-type-specific gene expression on semi-simulated data

Most benchmarking analyses of deconvolution focus on the ability of a given algorithm to recapitulate the proportion of cell types in every spot. In this setting, it is natural to use a clustered dataset of scRNA-seq data and generate synthetic “spot” measurements by sampling cells from different clusters with a given ground truth proportion. However, to fully assess the performance of DestVI to infer continuous cell states in addition to cell-type proportions, we instead built a more nuanced simulation framework that also accounts for variability within cell types (**Figure 2A, Online Methods**). In this scheme, each spot is defined by a cell-type proportion, as well as the (continuous) state of cells in every type. To model the continuum of cell states, we construct a linear model for every cell type, with a negative binomial likelihood. The coefficients of these linear models are learned using PCA on a cell-type-annotated scRNA-seq dataset. Using PCA ensures that the simulation is based on a probabilistic model different from the one used in DestVI. We then generate a spatial dataset by sampling, for each spot, its cell-type proportion and the coefficients of the cell state representations. We ensure spatial dependency in this data, by sampling those variables from a Gaussian process. The resulting simulator therefore generates spatial transcriptomics measurements while providing ground truth about cell-type-specific gene expression patterns (**Supplementary Figure 1**).

**Figure 2:**
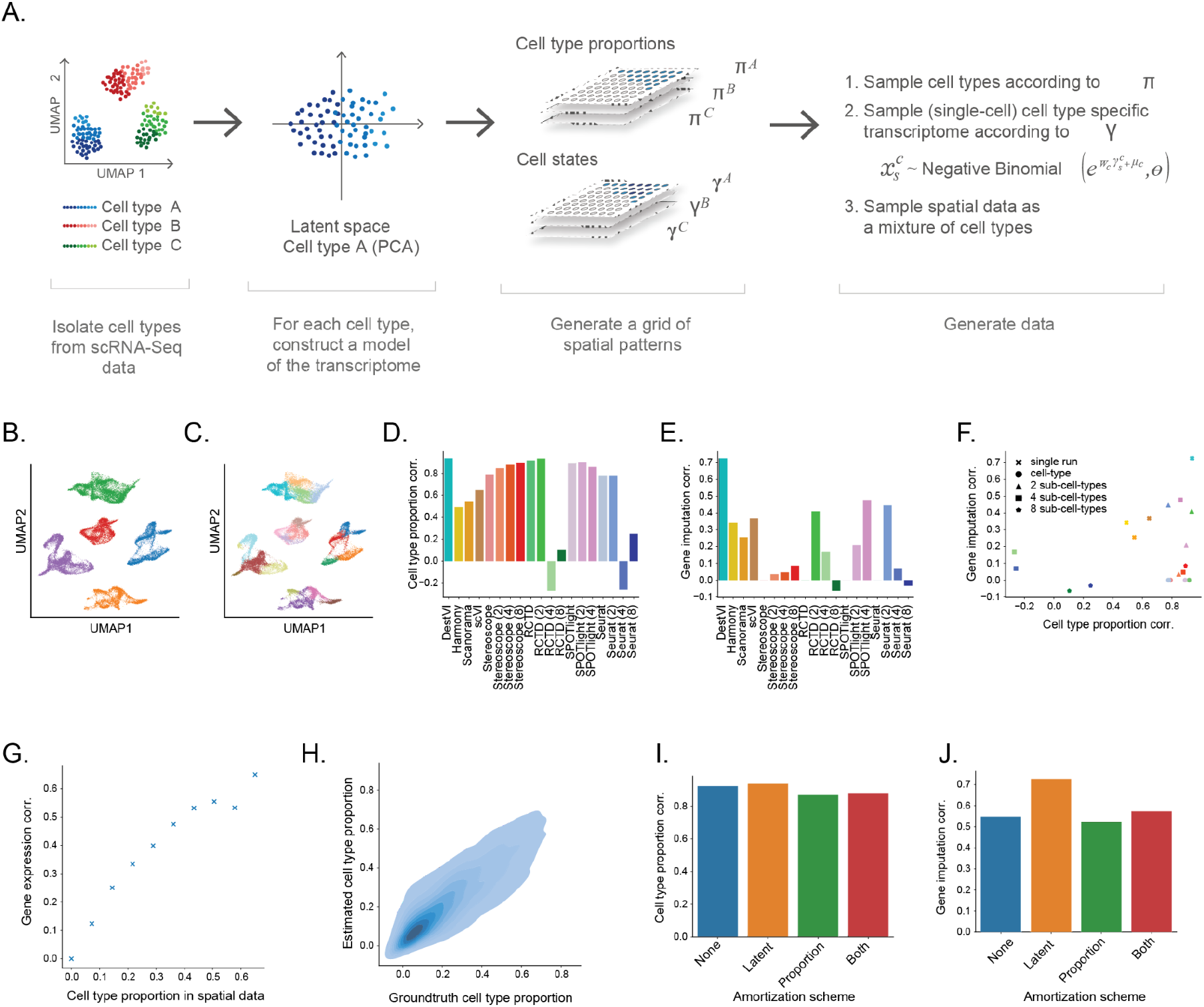
Evaluating the performance on DestVI on simulations. **(A)** Schematic view of the semi-simulation framework. We isolated five cell types from a single-cell RNA sequencing dataset. For each cell type, we learned a descriptive model of transcriptomic changes that are internal to that specific cell type, using a principal component analysis (PCA). We then generated a grid, in which each point represents a spot for the spatial transcriptomic simulation. We sampled spatially-relevant random vectors to encode the proportion of every cell type in every spot 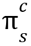, as well as the cell-type-specific embeddings 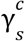. Then, we used those parameters, and the learned weights of the cell-type-specific PCA to generate spatial transcriptomics data (**Online Methods**). **(B-C)** Visualization of the single-cell data, and the cell state labels used for comparison to competing methods. Both scatter plots are UMAP embeddings of the single-cell data (32,000 cells). **(B)** Cells are colored by cell type. **(C)** Cells are colored by the sub-cell types, obtained via hierarchical clustering (here five clusters per cell type). **(D-F)** Comparison of DestVI to competing algorithms, possibly applied to different clustering resolutions. We allotted three hours for each simulation analysis. Performance is not reported for cases that did not terminate by that time (SPOTLight with 8 sub-clusters; **Online Methods**). **(D)** Spearman correlation of estimated cell-type proportions compared to ground truth for all methods. **(E)** Spearman correlation of estimated cell-type-specific gene expression compared to ground truth, focusing on combinations of spot and cell type for which the proportion is sufficiently high (here > 0.4 for the parent cluster) **(F)** Scatter plot of both metrics, that show the trade off reached by all methods. **(G-H)** Follow-up stress tests for DestVI. **(G)** Accuracy of imputation, measured via Spearman correlation as a function of the cell-type proportion in a given spot. **(H)** Head-to-head comparison of estimated cell-type proportion against ground truth across all spots and cell types (8,000 combinations of spot and cell type). **(I-J)** Ablation studies for the amortization scheme used by DestVI. We report performance metrics for variants of DestVI where different possible amortization strategies are used for learning the proportions as well as the cell states in every spot. **(I)** Spearman correlation of estimated cell-type proportions compared to ground truth. **(J)** Spearman correlation of estimated cell-type-specific gene expression compared to ground truth.

We compared the performance of DestVI to a number of leading benchmark methods. First, we compared to discrete deconvolution approaches: RCTD [16], SPOTLight [17], Stereoscope [18], and Seurat [32]. We consider the performance of these methods when trained with different levels of sub-clustering (only for algorithms that completed in less than three hours; **Figure 2B-C**), where the lowest resolution of clustering corresponds to the original cell types, and every subsequent resolution further partitions each type into distinct states. Using this range of sub-clustering levels helped explore the tradeoff between the ability to predict cell-type-specific gene expression (additional sub-clustering should be better, up to a certain extent) to the ability to accurately infer cell-type proportions, considering that sub-clustering may interfere with the ability to predict the frequency of the parent cluster. We also benchmarked against a second set of methods: scVI [29], Harmony [33] and Scanorama [34], that can be used to match the spatial measurements to the scRNA-seq measurements via a common embedding. In these approaches, inferences about cell-type proportions or cell-type-specific gene expression are done using k-nearest neighbors imputation (**Online Methods**).

To evaluate the accuracy of cell-type proportion estimates, we calculated the Spearman correlation between the inferred and ground truth proportions, and reported the average over cell types. We only considered combinations of spot and cell type for which the proportion is sufficiently high (here > 0.4). We also assessed how well each algorithm captures the variation within cell types, by calculating the Spearman correlation between the inferred and ground truth cell-type specific gene expression in every spot (again, reporting the average over all cell types).

Considering the results of cell-type proportion estimation (**Figure 2D**), we find that the embedding methods have a generally lower performance compared to other methods, specifically designed for deconvolution. This has been already reported (e.g., [20]) and is expected, as these embedding methods (e.g., scVI) do not explicitly consider that spatial spots may include a mixture of cell types. In the deconvolution methods, we find that the impact of the clustering resolution on accuracy is different for different algorithms. For example, performance is stable for Stereoscope and SPOTLight (with a slight trend up or down) but changes drastically for RCTD and Seurat where both algorithms perform poorly for more than four clusters per cell type. Finally, we find that DestVI compares favorably to the other methods, regardless of their sub-clustering resolution during training. Considering the results of cell-type-specific gene imputation (**Figure 2E**), we notice that, with the exception of DestVI, the deconvolution methods have a lower performance compared to the embedding-based ones. Specifically, for each discrete - deconvolution method there is often one resolution of sub-clustering for which it yields reasonable imputation results, but the optimal resolution is algorithm specific (four sub-clusters per cell type for SPOTLight; two sub-clusters per cell type for RCTD and Seurat). Since it is hard in practice to estimate the number of sub-clusters, especially in this specific context of deconvolution, this makes the discrete deconvolution approaches less applicable for spatial analysis. DestVI outperforms all methods with this metric and the same results follow using Pearson correlation as well (**Supplementary Figure 2**).

Taken together, these results demonstrate that DestVI provides a compelling alternative to discrete deconvolution algorithms, especially when there are rich continuous patterns of transcriptional variation within cell types, as is the case for most biological models. Specifically, we observe that DestVI demonstrates robust performance in gene expression imputation while still adequately estimating cell-type proportions (**Figure 2F**). Of note, our analysis was limited to spots in which the cell type in question was sufficiently abundant. As expected, we observe that the ability of DestVI to predict cell-type-specific gene expression decreases in the case of low frequency (**Figure 2G**). We do observe, however, much less effect on the accuracy of cell-type proportion estimate (**Figure 2H**). DestVI can therefore provide an internal control for which spots can be taken into account when conducting a cell-type-specific analysis of gene expression or cell state. We leverage this property and propose an automated way of estimating a threshold for the minimal cell-type proportion that is required for such an analysis (**Online Methods**).

Finally, we also utilized the simulation to benchmark several variants of DestVI (**Figure 2I-J**). Specifically, we wanted to verify our design choice for the analysis of spatial data in which we keep the parameters for the cell-type proportion free but treat the parameters 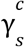 as a function of the input data (i.e., amortizing inference using an encoder neural network; **Online Methods**). Towards this end, we assessed the performance of several variants of DestVI, using an encoder neural network for the proportions, parameters 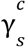, both of them, or none of them. Interestingly, we have noted that using a neural network for the proportions yield lower performance for cell-type proportion estimation compared to keeping free parameters (**Figure 2I**). Conversely, using a neural network for estimating the embedding variables yields much higher performance for gene expression imputation (**Figure 2J**). We attribute this phenomenon to the fact that there are many embedding variables to infer (dimension of latent space times number of cell types) per spot, and that using a neural network may regularize the solution as compared to using free parameters.

### An automated analysis pipeline with DestVI

The resulting model enables several types of downstream analyses for drawing hypotheses on both the spatial structure of an individual sample as well as the differences amongst pairs of conditions. In the following, we propose a standard pipeline for analysis of a single sample. In the subsequent sections we demonstrate how to use this pipeline to gain insight from a simple sample (a single lymph node, or tumor slice) and help guide comparative analysis between samples (between lymph node slices, or distinct areas of a tumor slice). The pipeline consists of two parts. In the first part, we consider the data from a resolution of cell types. For each cell type we report its proportion in every spot, and then highlight cell types that tend to occur at specific niches (i.e., not uniformly distributed across the tissue), using Geary’s C autocorrelation statistic for the inferred proportions [35]. The second part of the pipeline facilitates a more in-depth view - looking at variability within cell types, thus going beyond the functionality that is available in current pipelines. We start by selecting, for each type, spots that have sufficiently high proportion of cells of that type. We propose an automated way of estimating a cell-type-specific threshold for this procedure, but it may be also manually curated. We consider for each cell type different values of the threshold *t* (taken on a grid), and calculate *C*(*t*) - the respective Geary’s *C* statistics, accounting only for spots with proportion higher than *t*. We then select our threshold *t*^*^ to be the inflection point of the resulting *C*(*t*) curve (**Online Methods**). With this constrained view, our pipeline proceeds to report the main axes of variation in each cell type, using the spatial data. This analysis helps highlight and visualize the most dominant transcriptional programs in every cell type, as well as exploring their dependence on the cells’ locations. To this end, we developed a weighted PCA scheme that uses the inferred cell states (in the spatial data) and accounts for the inferred cell-type proportions as well as the spatial layout of the spots (following previous work on robust linear dimensionality reduction [36]; **Online Methods**). We also identify genes that are correlated with each weighted principal component and report enriched gene signatures (using EnrichR [37]) to help with their interpretation.

Our model also provides a natural way to estimate and evaluate the significance of differences between conditions, or between niches in the same tissue slice. Specifically, for each cell type we can compare the extent to which cells of a given type tend to co-localize in specific niches, by comparing the respective Geary’s C statistics. On the gene level, we can identify cell-type-specific differential expression, comparing different conditions or different tissue areas. This analysis draws directly from our probabilistic representation of the data, which allows for uncertainty quantification and hypotheses testing (**Online Methods**).

### DestVI identifies a spatially organized lymph node multicellular immune response following pathogen stimulation

For a first application of DestVI, we aimed to study spatial pattern of antigen-specific Immunity and profiled murine auricular lymph nodes following 48 hr stimulation by Mycobacterium smegmatis (MS), a gram-positive bacteria which induces a robust CD4+ T cell response characterized by IFNγ [24]. For spatial analysis, we used the Visium platform (10x Genomics) to profile four lymph node sections (two sections from MS and two sections from control (PBS) injections). A matching single cell RNA-seq data set was also obtained for these conditions (**Figure 3A; Online Methods**).

**Figure 3:**
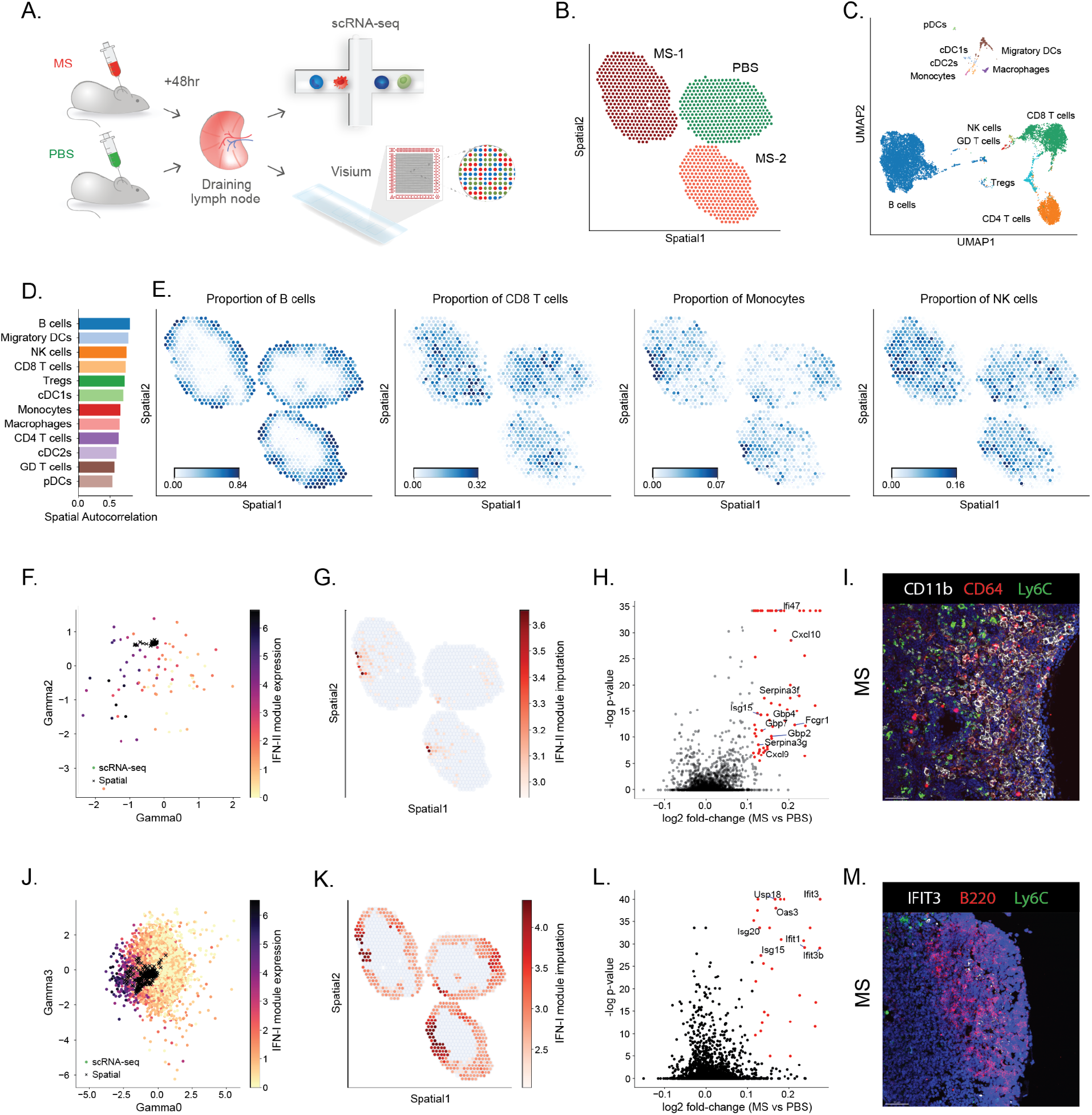
Application of DestVI to the murine lymph nodes. **(A)** Schematics of the experimental pipeline. We performed spatial transcriptomics (10x Visium) and single-cell RNA sequencing (scRNA-seq, 10x Chromium) on murine lymph nodes following 48 hr stimulation by Mycobacterium smegmatis (MS) compared with PBS control (two sections from each condition). **(B)** Visualization of the spatial transcriptomics data used for DestVI analysis (1,092 spots). Only three slices out of the four lymph nodes passed the quality check (**Online Methods**). **(C)** UMAP projection of the scRNA-seq data (14,989 cells), embedded by scVI and annotated by scANVI using a publicly available dataset of murine lymph node cells. **(D)** Spatial autocorrelation of the cell-type proportions for every cell type. **(E)** Spatial distribution of cell-type proportions for B cells, CD8 T cells, Monocytes and NK cells, as inferred by DestVI. **(F)** Joint embedding of the monocytes from the scRNA-seq data (circles; 128 cells) and the spots with high abundance of monocytes from the spatial transcriptomics data (crosses; 79 spots). Single-cell data is colored by expression of the selected IFN-II genes identified by Hotspot (Fcgr1, Cxcl9 and Cxcl10; see **Supplementary Figures 10-11**). **(G)** Imputation of monocyte-specific expression of the IFN-II marker genes for the spatial data (log-scale), reported only on spots with high abundance of monocytes (79 spots across the three slices). **(H)** Monocyte-specific differential expression analysis between MS and PBS lymph nodes (2,000 genes). Significance is calculated with our differential expression procedure (**Online Methods**). **(I)** Immunofluorescence imaging from a MS lymph node, staining for CD11b, CD64 and Ly6C. Scale bar, 50 μm. **(J)** Joint embedding of the B cells from the scRNA-seq data (circles, 8,359 cells) and the spots with high abundance of B cells from the spatial transcriptomics data (crosses, 579 spots). Single-cell data is colored by expression of the selected IFN-I genes identified by Hotspot (Ifit3, Ifit3b, Stat1, Ifit1, Usp18 and Isg15; see **Supplementary Figures 15-16**). **(K)** Imputation of B cell-specific expression of the IFN-I gene module on the spatial data (log-scale), reported only on spots with high abundance of B cells (579 spots across the three slices). **(L)** B cell-specific differential expression analysis between MS and PBS lymph nodes (2,000 genes). Significance is calculated with our differential expression procedure (**Online Methods**). **(M)** Immunofluorescence imaging from a MS lymph node, staining for IFIT3, B220 and Ly6C. Scale bar, 50 μm.

After quality control, we noticed that the number of UMIs per spot in one of the control lymph nodes was significantly lower than the other sections, and discarded this sample from further analysis (**Supplementary Figure 3**). After spot filtering on the remaining lymph nodes; a total of 400, 369 and 323 spots for the MS-1, MS-2 and PBS lymph nodes were used for analysis with DestVI, respectively (**Figure 3B; Online Methods**). As a preliminary test for the validity of the information obtained from 10x Visium, we began with a clustering analysis of the raw data using scanpy (**Supplementary Figure 4A**). We noted the high reproducibility of clusters for the MS conditions, as well as marked differences between the MS-stimulated and control (PBS) tissue sections, characterized by several clusters (**Supplementary Figure 4B**). The matching single cell RNA-seq data yielded 14,989 cells after quality filtering (**Online Methods; Supplementary Table 1**). We annotated the scRNA-seq data by transferring cell type information from a publicly available murine lymph node dataset [38] with scANVI [39] and manually curating the annotation of rare cell types (**Online Methods**). We present this information in a two-dimensional plot of the annotated latent space (**Figure 3C, Supplementary Figure 5**), as embedded by scVI [29], and laid out by UMAP [40]. We observe a similar cellular MS specific response including changes in NK and monocytes abundance and signaling (**Supplementary Figure 6**).

**Figure 4:**
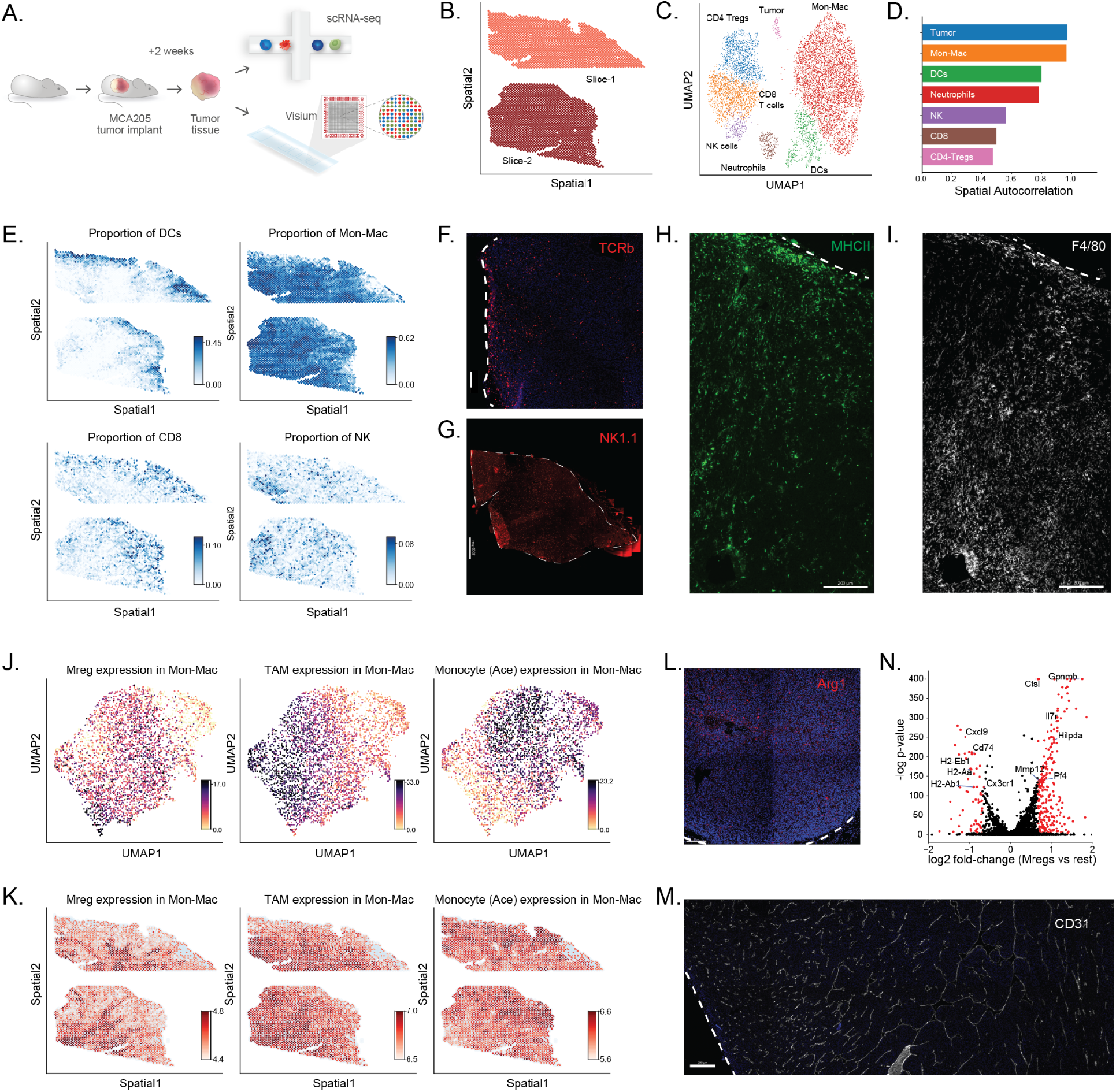
Application of DestVI to a MCA205 tumor sample. **(A)** Schematics of the experimental pipeline. We performed spatial transcriptomics (10x Visium) and single-cell RNA sequencing (scRNA-seq, single-cell MARS-seq protocol) on a MCA205 tumor which contains heterogeneous immune cell populations 14 days after intracutaneous transplantation into the wild-type mouse (two sections). **(B)** Visualization of the spatial transcriptomics data for two tumor slides, after quality control (4,027 spots). **(C)** UMAP projection of the scRNA-seq data (8,051 cells), embedded by scVI and manually annotated. **(D)** Spatial autocorrelation of the cell-type proportions for every cell type, computed using Hotspot. **(E)** Spatial distribution of cell-type proportions for DCs, monocytes and macrophages (Mon-Mac), CD8 T cells and NK cells. Zones emphasized with black dashed lines on the lower-right-hand corner plot designates high-density of NK cells discovered by DestVI. **(F-I)** Immunofluorescence imaging from the tumor using antibodies for **(F)** TCRb. Scale bar, 100 μm. **(G)** NK1.1. Scale bar, 1000 μm. **(H)** MHC II. Scale bar 200 μm. **(I)** F4/80. Scale bar 200 μm. White dashed lines in panels F-I indicate the tumor boundary. **(J)** Visualization of three gene expression modules on the Mon-Mac cells from the scRNA-seq data (4,400 cells), on the embedding from scVI (identified using Hotspot; see **Supplementary Figures 25-26**). These gene modules are named according to our previous myeloid single-cell analysis of the MCA205 tumor [50]. **(K)** Imputation of gene expression for those modules on the spatial dataset (log-scale), reported only on spots with high abundance of Mon-Mac (3,906 spots across the two slices). **(L)** Immunofluorescence imaging from the tumor using a Arg1-YFP transgenic mouse. Scale bar, 150 μm. **(M)** Immunofluorescence imaging from the tumor using antibodies for CD31, showing the abundance of blood vessels in the tumor (quantitative metrics in **Supplementary Figure 30**). Scale bar, 400 μm. **(N)** Mon-Mac cell-specific differential expression analysis between the Mreg enriched zone and the rest of the tumor slices (2,886 genes). Significance is calculated with our differential expression procedure (**Online Methods**).

We applied DestVI in order to explore for each cell type how infection-induced differences in transcriptional states may be associated with changes of spatial organization. The reports that are automatically generated by our analysis pipeline are provided in **Supplementary Note 1**. These reports helped guide our exploration of the data, and we summarize the resulting main findings in the remainder of this section. Using the first part of our automated pipeline, we began by exploring the spatial distribution of cells from each type (**Figure 3D-E; Supplementary Figure 7)**. As expected, the first striking pattern is the organization of the lymph node sections into the B cell follicles (Cd79a; external area) and a T cell compartment (CD8; internal area) (**Figure 3E; Supplementary Figure 8**) [41,42]. Interestingly, we find that monocytes and NK cells also tend to form spatially coherent niches, with a stronger extent of co-localization of the monocyte population in the stimulated lymph node compared to the control (comparing the Geary’s C values of their inferred proportions; **Figure 3E; Supplementary Figure 9**). This finding is consistent with our previous work, which showed that after immunization with MS, NK cells are recruited to the lymph nodes and produce IFNγ. This signaling axis further promotes the up-regulation of IFNγ signaling in monocytes [24]. The spatial data, further identified organized multicellular immune response niches, located between the B cell follicles as shown in **Figure 3E** and also referred to as the interfollicular area (IFA) [43].

We then explored the inferred spatial organization of cell states within every cell type, a unique feature of our pipeline compared to other algorithms for spatial analysis (**Figure 3F-M**). Interestingly, we find that the main axes of variation in the monocyte population (using our weighted PCA scheme to co-analyze both modalities) reflect cytokine and chemokine signaling cascades, as well as a robust interferon response (**Supplementary Note 1**). The importance of interferon signaling as a source of variation in the monocyte population is also apparent in the scRNA-seq data (using Hotspot [38]; **Supplementary Figures 10-12**), and characteristic of molecular differences between MS stimulated dissociated lymph nodes and control lymph nodes (**Supplementary Figure 13**). However, through joint analysis with DestVI, we can now contextualize this variation with the location of the monocytes and their unique spatial niche in the lymph node. To this end, we first inspected our inferred low-dimensional representation of cell state and found that it indeed reflects co-variation in the expression of type-II interferon response genes (here, plotting the cells and spots in dimensions 0 and 2 of the 5-dimensional γ vector; colored by the sum of Fcgr1, Cxcl9 and Cxcl10; **Figure 3F**). Moving to the tissue coordinates, we found a clear localization of inferred monocyte-specific expression of IFN-II genes in the IFA (**Figure 3G**). Comparing the two conditions, we also found that the amount of expression in those constrained niches is markedly higher in the MS versus the control LN. In order to more formally identify the differences in expression of monocytes across the two conditions, we performed differential expression (using the values imputed by DestVI) and discovered a rich set of differentially expressed IFN-II response genes, some already used in our signature analysis (e.g., Fcgr1 and Cxcl9), but also other genes (Gpb2, Serpina3g and Ifi47) (**Online Methods, Figure 3H, Supplementary Table 2, Supplementary Figure 12**). This is consistent with previous findings, where under pathogen immunization monocytes or macrophages carrying antigens migrate to draining lymph nodes through afferent lymphatics to induce immune responses [44–48]. In order to verify this discovery using alternative spatial measurements, we performed immunofluorescence staining of LN from naive and MS treated mice,with CD64 (Fcgr1; a marker we identified with DestVI), CD11b and Ly6C. In line with the results identified by DestVI, we find a higher number of inflammatory monocytes in the MS treated LN localized to the IFA and the peripheral subcapsular sinus (SCS) (**Figure 3I, Supplementary Figure 14**).

Our results also point out to functional heterogeneity within the B cell compartment, with a strong enrichment for type I interferon signaling in the first weighted principal component (**Supplementary Note 1**). As above, we find that this axis of variation is also captured in the scRNA-seq data (**Supplementary Figures 15-17**), and thus proceed to explore its spatial properties. First, we find that it is indeed captured by the joint latent representation of cells and spots (dimension 0 of the 5-dimensional γ vector; colored by Ifit1, Ifit3, Ifit3b, Stat1, Usp18 and Isg15; **Figure 3J**). We then use the B-cell specific gene expression estimates from DestVI to inspect the spatial organization of this signature (**Figure 3K**). Interestingly, the module appears to be expressed across all the lymph nodes, but in lower levels for the PBS LN compared to the MS LN. This observation is also supported by B-cell specific differentiation expression analysis between the MS and control tissues (**Figure 3L, Supplementary Table 3, Supplementary Figure 17**). In particular, we investigated the B-cell specific differential expression analysis within the MS samples with focus on comparison of the enriched zone with the rest of the lymph node slices and noticed a similar strong signature of type I interferon signaling (**Supplementary Table 4**, **Supplementary Figure 18**). These observations were further validated by immunofluorescence staining for IFIT3, B220 and Ly6C (**Online Methods**). We identify IFIT3^+^B220^+^ cells on the MS sample near the SCS and IFA (**Figure 3M**) but not in control samples (**Supplementary Figure 19**). Applying DestVI to spatial transcriptomics data, we detected the unexpected finding of spatially-enriched interferon reaction in B cells.

In summary, the unique features of our analysis enable robust spatial characterization of cell types and states within naive and pathogen challenged lymph nodes. DestVI identifies a clear and specialized immune niche involving IFN signaling of different cell types, including monocytes and B cells activated by MS of infected mice and localized to the peripheral subcapsular sinus and interfollicular area.

### DestVI identifies an hypoxic population of macrophages within the tumor core

We also explored DestVI performance in a complex and less structured tissue. Towards this end, we spatially profiled a syngeneic mice tumor model (MCA205) using Visium. Fourteen days after intracutaneous transplantation of MCA205 tumor cells, we have characterized the tumor using scRNA-seq and Visium (**Figure 4A; Supplementary Figure 20; Online Methods**). A total of 2,125 spots (first tumor section) and 1,902 spots (second tumor section) were used for analysis with DestVI following quality metrics and filtering (**Figure 4B**). Along with the Visium data, we also collected cells from a separate tumor for single-cell RNA sequencing. After processing and filtering, this dataset comprised a total of 8,051 immune cells and tumor cells (**Online Methods; Supplementary Table 5**).

We annotated the single cell RNA-seq data by labeling clusters of the latent space from scVI [29] based on marker genes from immune cells (**Online Methods**). We present this information in a two-dimensional plot of the annotated latent space (**Figure 4C**), laid out by UMAP [40]. We then explored the spatial distribution of these cell types and states using DestVI. After running DestVI, we applied our post-hoc interpretative pipeline (**Online Methods**) and reported the whole analysis in **Supplementary Note 2**. We first inspected the cell-type proportions of the major immune subsets (CD8 T cells, monocytes, macrophages, dendritic cells, and NK cells; **Figure 4D-E**; remaining cell types in **Supplementary Figure 21**). We observed that both types of T cells were highly abundant on the boundary of the tumor. We further verified this observation by staining for T cell specific markers (TCRb) using immunofluorescence (**Figure 4F,** larger field of view in **Supplementary Figure 22**) [49]. We also observed that NK cells occupy very specific niches within the tumor and verified this spatial property by staining for NK1.1 (**Figure 4G;** a slice at distance 30 microns from Slice-2 and 80 microns from Slice-1; larger image in **Supplementary Figure 23**). Direct comparison of the images is difficult because the slice used for staining is different from the one used in visium, with the visium slice truncated by the visium capture area (**Supplementary Figures 24-25**). However, we notice a similar pattern of NK cells in DestVI and the immunofluorescence staining. Antigen-presenting DCs also exhibited non-uniform spatial organization, with marked localization at the boundary of the tumor (**Figure 4D**), a property we further tested by staining for MHCII (**Figure 4H**). Finally, we noted that monocytes and macrophages (jointly labeled as Mon-Mac in the scRNA-seq data; as there was no clear demarcation in latent space) were present broadly in the tumor, with no specific pattern (**Figure 4D**), and we verified this by staining for F4/80 (**Figure 4I**). Together, these results suggest that DestVI is able to provide a reproducible overview of the organization of major immune subsets in the tumor.

Since the Mon-Mac population did not have a specific spatial pattern (unlike DCs, NK cells and T cells), we hypothesized that the spatial coordinates may reflect different cell states within the Mon-Mac population. Indeed, using DestVI for analysis of spatial patterns within the Mon-Mac population reveals a stratification of this subset into spatial niches, each with a distinct expression signature (**Supplementary Note 2**). We therefore proceeded to an in depth analysis of the Mon-Mac populations.

We started by exploring the scRNA-seq data and, using Hotspot, identified several gene expression programs that distinguish different states within the Mon-Mac population (each represented by a different module of co-expressed genes; see **Supplementary Figures 26-27**). This observed variation in cell state matched our previous findings in independent biological replicates of the same tumor system [50]. Three of the detected modules of co-expressed genes pertain to general monocytes markers and help distinguish a Monocyte (Ace) gene expression program (defined by Ly6c2, Plac8, Ly6a, Ace, Ear2), a Mon-DC program characterized by MHC class II genes (H2-Aa, Cd74, H2-Ab1), and a Monocyte (IFN) program characterized by type I interferon signaling (Ifit1, Ifit2, Ifit3, and Isg15). The remaining two modules define specific tumor-infiltrating myeloid suppressive cell populations [50]. The first one corresponds to a Mreg population discovered previously [50], with expression of Trem2, Gpnmb, Mmp12 and Il7r, as well as markers of hypoxia - Hmox1, and Hilpda. The second module corresponds to tumor associated macrophages (TAM), expressing C1qa, C1qb, C1qc, Ms4a7 and Apoe (**Figure 4J**).

To inspect the spatial distribution of these subpopulations, we used DestVI to infer the Mon-Mac specific expression of the corresponding gene modules in our Visium data (**Figure 4K**; The spatial distribution of all other modules is displayed in **Supplementary Figure 28**). Interestingly, we noticed several spatial patterns. First, the tumor-infiltrating myeloid suppressive cells (Mreg and TAM) were mostly abundant in the inner layers of the tumor. We hypothesized that the inner-tumor macrophage state detected by DestVI corresponded to the population of regulatory macrophages (Mreg) which express Trem2 and Arg1 and has been previously observed in human tumors [51] as well as murine models [50]. These Arg1^+^ myeloid cells are associated with poor antitumor response [52] and have interestingly been observed to congregate in hypoxic tissue niches (cancerous [53] and non-cancerous [54]). To validate this observation and localize these unique myeloid cells within the tumor, we have used an Arg1-eYFP transgenic mouse model (YARG) (**Figure 4L**). We found that inner-tumor had more Arg1-eYFP positive cells than the boundary of the tumor. To further validate the inner hypoxia tumor microenvironment, we have stained CD31 to detect the presence of blood vessels (**Figure 4M**). As expected, the density of blood vessels decreases in the inner layers of the tumor, which suggests the deprivation of oxygen induces hypoxia regulated signaling in macrophages (larger field of view with quantitative metrics in **Supplementary Figure 29**). A more striking spatial pattern is the expression of hypoxia genes, underlying the localization of the Mreg population (**Figure 4K**). We have further characterized this myeloid population by searching for Mon-Mac specific differentially expressed genes between the hypoxia area (Mreg) and the rest of the tumor slides (**Supplementary Figure 30**), and identified known markers from including Ctsl and Il7r [50] (**Figure 4N, Supplementary Table 6**).

In summary, DestVI correctly maps cell types of immune cells onto the spot coordinates and identifies a clear and specialized niche involving metabolic changes in response to hypoxia within the macrophage population.

## Discussion

We introduce DestVI, a multi-resolution approach to deconvolution of spatial transcriptomics profiles using an auxiliary single-cell RNA sequencing dataset. Via simulations, we show that classical deconvolution approaches that are based on clustering the scRNA-seq data may be difficult to apply and miss important information in the case of marked variation within cell types. DestVI circumvents this problem by learning cell-type-specific latent variables on the scRNA-seq data, using a deep generative model, and mapping those latent variables onto the spatial data. Coupled to the automated pipeline we developed, DestVI is capable of interpretable analyses to compare the within-cell-type gene expression levels across different conditions, or different niches of the same tissue slice.

An important feature of our work is the ability to perform cell-type-specific differential expression in the spatial data. Notably, because we do not perform fully-Bayesian inference for the spatial latent variable model (MAP inference is used), we only obtain point estimates of the cell-type-specific cell states 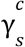. As a result, we must use a frequentist test for the differential expression. A more principled approach, and a subject for future work, is to apply variational inference and exploit the uncertainty for Bayesian differential expression [55,56].

Although deconvolution of spatial transcriptomics data has recently received considerable attention, none of the methods benchmarked in this manuscript (including DestVI) make explicit use of the spatial coordinates during inference (e.g., inference of cell type proportions in a given spot should be influenced by its nearby spot). A plethora of computational techniques, based on black-box inference techniques [57,58] for Gaussian processes [59] is likely to make this possible. However, the level of technical development to achieve this may not be reasonable, because the parameters studied in this work (cell-type abundance) may not be spatially smooth (e.g., neighboring spots may be very different) for some tissues, such as the tumor. Consequently, we developed a simpler modification of DestVI that takes into account enforces smoothness of the cell-type abundance parameters over the spatial coordinates with a spatially-aware penalization based on a quadratic cost (the scaling factor can be set via cross-validation, holding out parts of the transcriptome). This promotes neighboring spots to have similar cell-type abundance, and improves the results on the simulations (**Supplementary Note 3**).

Spatial transcriptomics is a promising approach for unravelling cell-cell interactions [60] and other forms of cellular communication and function in a tissue [61]. We expect that approaches such as DestVI will provide the necessary level of resolution and help further our understanding of the local signaling environments and how they impact cell functions and spatial cues, such as interaction between specific cellular subsets, chemical gradients and metabolic cross talk.

## Online Methods

### Model of single-cell RNA sequencing data

#### Assumption and model for the single-cell data (scLVM)

Let *n* designate a cell in the scRNA-seq dataset. We assume that each cell is annotated with cell-type label *c*_*n*_, but those labels are broad enough such that the introduction of continuous covariates γ_*n*_ into the model helps explain additional variance in gene expression (i.e., within-cell-type variation). For example, *c*_*n*_ represents a discrete cell type (e.g., B cells or CD8 T cells) while γ_*n*_ is a continuous vector summarizing a sub-cell state of interest (e.g., B cell activation, CD8 T cell exhaustion).

In the following, we assume that *c*_*n*_ is observed (e.g., obtained via clustering) and that γ_*n*_, however, is unobserved and treated as a latent variable. We posit the following generative model for our data:

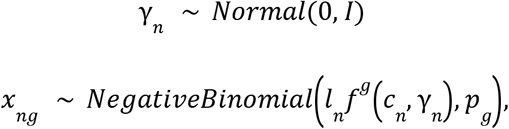

where *l*_*n*_ is the library size of cell *n*, *f* is a two-layers neural network and *p* is a *G*-dimensional vector. *f* takes as input the concatenation of the one-hot encoding of *c*_*n*_, as well as the scalar γ, and outputs a *G*-dimensional vector. *f* has an softplus non-linearity at its output to ensure positivity.

This generative model has significant overlap with our previous proposed method single-cell Variational Inference (scVI [29]), which is also a conditional deep generative model. On top of the conceptual difference that scVI conditions on the batch identifier, whereas scLVM conditions on the cell-type information, there are several technical points in which these two models differ. First, we use the standard parametrization (*r*, *p*) for the negative binomial distribution: the number of successful independent and identical Bernoulli trials before *r* failures are achieved, in which *p* is the probability of failure of each Bernoulli trial. This is in contrast with scVI that relies on the mean-dispersion parameterization for the negative binomial distribution, and is necessary to make the definition of the spot gene expression level as the sum of contributions from individual cells correct (as emphasized in [18]). Furthermore, changes were required to the neural network architecture for the transfer learning to work adequately. Indeed, we found that using a decoder with randomness such as dropout [62] or with running parameters as in batch normalization [63] did not work, so we replaced those with layer normalization [64].

#### Variational inference

We use auto-encoding variational Bayes [30] to optimize the marginal conditional likelihood *log p*_θ_(*x*_*n*_|*l*_*n*_, *c*_*n*_). We use a mean-field Gaussian variational distribution *q*_ϕ_(γ_*n*_|*x*_*n*_, *c*_*n*_), parameterized by a two-layer neural network *g*. This neural network takes as input the concatenation of the gene expression vector *x*_*n*_ as well as the one-hot encoding of the cell-type label, and outputs the mean and the variance of the variational distribution for γ_*n*_. We optimize the evidence lower bound:

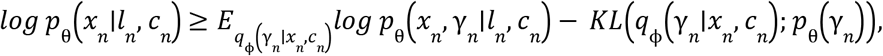

where *p*_θ_(γ_*n*_) denotes the prior likelihood for γ_*n*_. For this, we subsample the observations in mini-batches and we sample from the variational distribution using the reparameterization trick. Additionally, we reweight cells by their inverse cell-type proportion (capped with a minimal proportion of 5%). We have found this to be an effective method for learning a better representation of the lowly abundant cell types (e.g., monocytes in the lymph node).

### Model of spatial transcriptomics data

#### Assumption for the spatial data

In the spatial data, we assume that the gene expression of each spot is the combination of multiple cells, each potentially being from different cell types. A standard modeling assumption is that a spot *s* has for expression *x*_*s*_ the sum of individual cells [16,18]. Similarly, let us assume each spot has *C*(*s*) cells, and that each cell *n* in spot *s* is generated from latent variables (*c*_*ng*_, *γ*_*ng*_). We then have a distribution of gene expression:

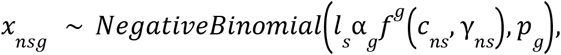

with *l*_*s*_ is a spot specific scaling factor and α_*g*_ is a correction term for the difference in experimental assays. From the standard property of the rate-shape parameterization of the negative binomial distribution, the distribution of the total gene expression *x*_*sg*_ in spot *s* and gene *g* is:

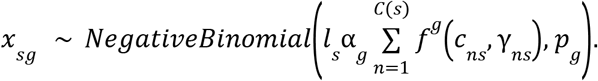

We now assume that all the cells from a given cell type *c* in a given spot *s* must all be generated from the same covariate 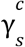. Instead of determining the cell identity of all individual cells in the spot, we focus on determining the density into broad cell types, as well as the archetype of the sub-cell state, which is a simpler problem. More concretely, we assume that there cannot be both significantly different cell states of the same cell types within a radius of 50 microns (i.e., a spot).

#### Generative model

These points in mind, we parameterize the sum in the previous equation to be over cell types. We obtain the following generative process:

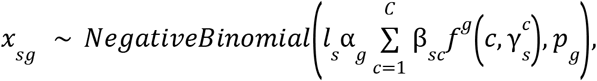

where *f* and *p* denote respectively the decoder network and the rate parameter of the negative binomial, transferred from scLVM. The gene-specific multiplicative factor α explicitly controls for discrepancies between the technologies. Parameters β_*sc*_ are positive, and designate the abundance of every cell type in every spot. These parameters may be normalized per spot to return an estimate of the cell-type proportion. In our implementation, we also add a constant term that serves as an unknown cell-type, as in [18].

An important technical component of DestVI is the empirical prior we use for the per-spot per cell type 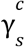 latent variable. Indeed, the model is susceptible to factor technology discrepancies into the latent space γ^*c*^ instead of the multiplicative factor α if an informative prior is not used and we noticed this pathological behavior with an isotropic normal prior. Consequently, we designed an empirical prior based on the single-cell data, for each cell type *c*:

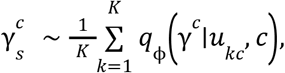

where 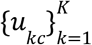 designates a set of cells from cell type *c* in the scRNA-seq dataset, and *q*_Φ_ designates the variational distribution from the single-cell latent variable model. In another context, this prior over γ^*c*^ is referred to as a variational aggregated mixture of posteriors (VampPrior, [31]). However, the objective here is simply to use the points 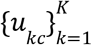 as a more informative prior for deconvolution, the VampPrior method seeks to learn a multi-modal, more complex prior in order to better fit the data.

#### MAP inference

We infer point estimates for random variables γ^*c*^ and for parameters α, β using a penalized likelihood method. In addition to vanilla MAP inference, we introduced two key ideas that stabilized the performance of DestVI. First, we added to the likelihood a variance penalty for the parameter α, calculated across all the genes. This strategy was applied previously by ZINB-WaVE to regularize estimates of dispersion parameters in their likelihood-based matrix factorization of single-cell data [65]. Second, compared to a standard deconvolution model that has exactly *C* parameters per spot, stLVM has *C* parameters and *dC* latent variables per spot, where *d* denotes the dimension of the latent space learned by scLVM. In order to avoid overfitting, we therefore proposed to use a neural network to parameterize the latent variables as a function of the input data (as in auto-encoding variational Bayes). Namely, we proposed several variants of the algorithm in which either both, part of or none of β and γ^*c*^ may be parameterized by neural networks. Intuitively, the use of a neural network for inference of γ^*c*^ may be helpful whenever there are shared transcriptomics profiles across cell types (such as inflammatory signals). These points in mind, the objective function to train this generative model is simply composed of (i) the negative binomial likelihood (ii) the likelihood of the empirical prior and (iii) the variance penalization for α.

### Simulations and data generative process

The benchmarking of spatial deconvolution methods often relies on using a single-cell RNA sequencing dataset with cell-type annotation, and aggregating multiple cells into a pseudo-bulk using a ground truth proportion. The algorithms are then evaluated on the prediction of the cell-type proportions [18]. This approach is not entirely adequate to our setting, as we would like to assess imputation of cell-type-specific variations of the transcriptome that are lost when combining together random cells of a given cluster. Instead, we used a real dataset of scRNA-sequencing data to simulate paired spatial and single-cell transcriptomics data. This helps properly benchmark our method against existing deconvolution methods.

#### Learning cell-type-specific transcriptomic modules from single-cell data

We simulated cell-type-specific transcriptional modules based on the lymph node scRNA-seq dataset of this manuscript. Out of all the cells, we kept five cell types: B cells, CD4 T cells, CD8 T cells, migratory DCs and Tregs. Those cell types were selected because they were the most abundant in the dataset. For each cell type, we learned patterns of transcriptomic variation within each cell type using a sparse PCA model, on log-normalized data [66]. We favored sparse PCA over classical PCA because for some cell types, the number of cells was much lower than the number of genes (e.g., around *N* = 300 cells for the regulatory T cells and *G* = 2, 000 genes). The counts were normalized using scanpy [67] with a target count of 10,000 UMIs and the sparse PCA model was fit using sklearn [68], using four components and a Lagrange multiplier of 5 for the ∥. ∥_1_ penalty. The output of this procedure is a cell-type-specific embedding for every cell γ_*n*_, a mean expression profile for every cell type μ_*c*_, and a dictionary of within cell-type transcriptomic variation *W*_*c*_.

The parameters μ_*c*_ and *W*_*c*_ may be explicitly used to build a simulation process for generating single-cell data *x*, given its cell type *c* and embedding γ:

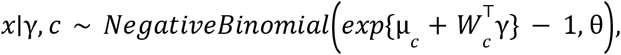

where we used the mean-dispersion parameterization for the negative binomial distribution. The dispersion parameter θ_*g*_ for every gene *g* is estimated from the data using scVI [29].

#### Generating spatial maps of transcriptome

To build the spatial transcriptomics data, we first constructed a regular grid of dimension 40 × 40. Each spot *s* (i.e., point on the grid) is associated with spatial coordinates *t*_*s*_. For the cell-type proportions π_*s*_, we built a covariance matrix *K* based on the spatial coordinates:

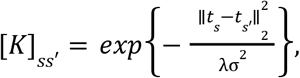

where σ denotes the median distance between all pairs of points in the grid. The λ parameter controls the level of spatial smoothness of the stochastic process and is fixed to 0. 1. This kernel matrix *K* may be used to sample *c* independent draws from a Gaussian process:

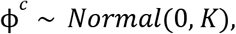

and interpret them as an energy to derive cell-type proportions at every spot:

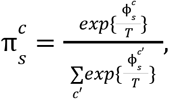

where *T* is a temperature parameter, set to 1. Large *T* would correspond to all proportions to be equal to 20%, while small *T* would tend to make the proportions binary. Regarding the embedding variables for every cell type 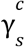, we treat them as 4*C* independent draw from the same kernel.

For every spot *s*, we sample a fixed number of cells *K*. For every single-cell, we decide on its cell-type *c* based on a draw from the categorical distribution parameterized by the proportions π_*s*_. We then use the previously introduced simulation model to generate the transcriptome of that cell (of cell type *c* and embedding 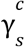). In our simulation, the scRNA-seq dataset has *K* = 20 cells per spot. Finally, for every spot, we average the mean parameter of the negative binomial distribution across all *K* cells and sample from the same observation model.

#### Comparison to competing methods

Our major claim is that DestVI is able to infer cell-type proportions but also to detect within cell-type transcriptomic variations in the spatial transcriptomics data. Although cell-type proportion estimation methods are reasonably simple to benchmark based on simulations, there is more ambiguity with respect to the second task. We provide a robust evaluation of the performance of algorithms at identifying continuous cell states within each cell type, based on cell-type-specific gene expression imputation.

We therefore provided a slight modification of every algorithm so that it may be used to impute gene expression at a given cell type *c* and a given spot *s*. For example, DestVI directly performs this task by accessing the inferred variable 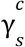 and decoding as 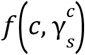, which serves as an unnormalized gene expression. For algorithms based on embeddings and nearest-neighbors imputation (scVI, Harmony, Scanorama), we impute by calculating the average gene expression of the *k*-nearest neighbors of the embedding of the spot *s* restricted to the single-cell data of cell type *c* only. Similarly, we estimate the proportion using the empirical proportion of cell types for the *k*-nearest neighbors of the embeddings of the spot *s* restricted to all the single-cell data. Finally, we also evaluate the performance of discrete deconvolution algorithms (Stereoscope, Seurat, RCTD and SPOTLight) for within cell-type gene expression. For this, we re-cluster the single-cell data for each cell type at several depths, using hierarchical clustering (2, 4 and 8 clusters per cell type). Then, we run the discrete deconvolution methods on the re-clustered data. We also calculate the mean gene expression for every cluster based on the single-cell data. Finally, we impute gene expression for cell type *c* at spot *s* by averaging the gene expression of every cluster in cell type *c*, weighted by the conditional proportions of every cluster at the spot. Given these modifications, we evaluated the imputation based on correlation metrics (e.g., spearman correlation) across an oracle list of genes for each cell type. The list of genes for each cell type is given by the indices of the non-zero coefficients of the matrix *W*_*c*_, learned via sparse PCA. We have found that this selection of genes helps make the evaluation more robust.

Interestingly, SPOTLight did not terminate after three hours for the most granular clustering (8 clusters per cell type).

### Automated pipeline for data exploration

Although DestVI may be used in a biologically informed setting, when practitioners are seeking for spatial relevance of specific transcriptomic modules in a given cell type (e.g., interferon type I response in B cells), we also designed a more agnostic pipeline to quickly explore, interpret and visualize the results of DestVI.

#### Spatially localized cell types

In this first step we simply calculate the auto-correlation index as reported by Hotspot [69] on the cell-type proportion for each individual cell type.

#### Selecting informative thresholds for cell-type proportions

We wanted to select the threshold based on the spatial information. Therefore, we chose a characteristic point of the auto-correlation curve, as a function of the thresholding. More precisely, for every cell type *c*, we apply the hard-thresholding operator *T* to the proportion vector β^*c*^ at different levels *t*_*i*_ (noted *T*(β^*c*^, *t*_*i*_). We take *t*_*i*_ to be the empirical percentiles of the proportion across all spots. For each thresholded vector *T*(β^*c*^, *t*_*i*_), we then calculate the auto-correlation metric from Hotspot [69] for all of those thresholded proportions. The result is a curve, that may be interpolated using splines using the scipy.interpolate function from SciPy [70]. We then analytically differentiate the spline and look for an inflection point (null second derivative).

#### Finding main axes of variation in the combined data

Interpreting the cell-type-specific latent dimensions of DestVI may be challenging. We therefore present here a visualization technique that aims at summarizing the inferences. The major question we would like to solve here is the following: within a single cell type, which directions of γ vary spatially? In other words, we wish to find within cell-type and spatially-varying transcriptomic programs.

General matrix factorization methods (such as NMF, PCA or CCA) could be applied to find those important components. However, they present several crucial limitations: (i) the samples are not reweighted by the cell-type proportion. This is especially important because γ_*c*_ is inferred at every spot, even when β^*c*^ is null, (ii) we are only interested in variations that are relevant with respect to the spatial coordinates λ.

To identify those, we focus on identifying the directions of γ that vary the most (as in PCA), but while enforcing some agreement with respect to the spatial location, and by taking into account the cell-type proportions. More precisely, let *c* be a fixed cell type. We define first spatial PC of the γ^*c*^ space, noted *u* ∈ *R*^*d*^, as the argument of the solution to the variational problem:

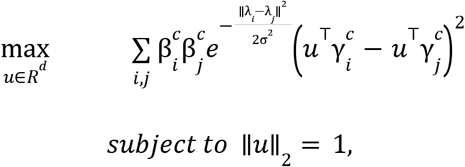

This problem, whether we are interested in a single or in several PCs, is tractable, as it may be formulated as an eigenvalue problem. We present this formulation below, providing a more succinct presentation than in [36]. We then explain in further details how this weighted PCA is related to a classical PCA. Interestingly, the previous optimization problem could be extended to find pairs of cell-type-specific transcriptional programs that are spatially co-activated (via a weighted CCA), but we leave this to future work.

#### Principal Component Analysis (PCA)

Let (*x*_1_, …, *x*_*n*_)denote a dataset, in which each datapoint *x* ∈ *R*^*d*^. The problem of finding the first principal component *u*_*PC*_ may be formulated as finding the direction of maximal variance in the data:

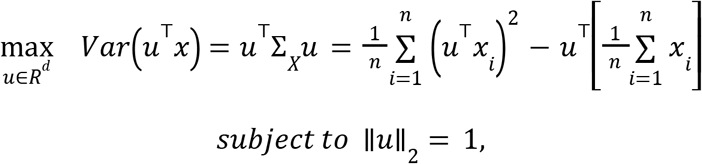

#### Laplacian-weighted PCA

We wish to incorporate sample-to-sample weights into this objective function, so that we can accordingly reweights pairs of observations according to a non-negative, symmetric dissimilarity matrix *d*_*i,j*_. From this matrix, we form the Laplacian matrix *L*^*d*^ ∈ *M*_*n,n*_(*R*). A Laplacian matrix is a PSD matrix with zero-sum rows and columns. Then, relying on the the identity 2*Var*(*x*) = *E*[(*x*_1_ − *x*_2_)^2^], in which *x*_1_ and *x*_2_ are iid copies of *x*, we define the first weighted principal component 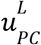 as the solution of the following optimization problem:

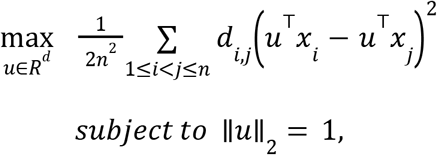

We now argue that this optimization problem is no different than the original PCA formulation, and can be solved via eigenvalue decomposition. This is because the objective function from the weighted optimization problem is still the evaluation of a quadratic form:

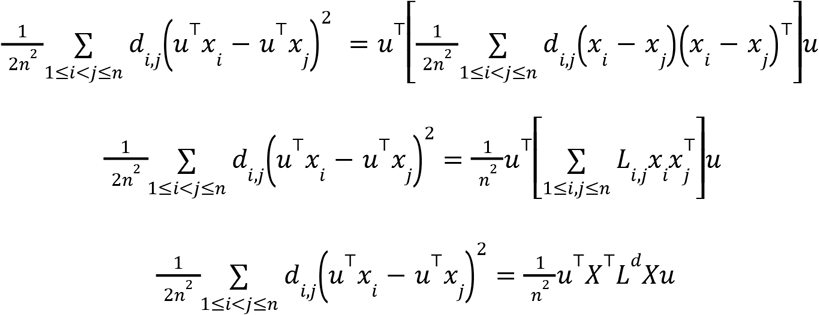

Consequently, we can find the Laplacian weighted principal components via eigenvalue decomposition of the matrix *X*^T^ *LX*. When the random variable *x* is centered, and the dissimilarity matrix is trivial, we obtain *X*^T^*LX* = Σ_*X*_. More importantly, it seems we can rotate the data 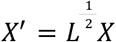 and solve PCA on it.

#### Interpretation of the weighted PCA

For interpreting the spatially-weighted PCA for every cell type, we first project the cell-type-specific cell state of the spatial data as well as the scRNA-seq data onto the two weighted principal components. We then use a two-dimensional colormap to assign a color to the spots in the spatial data, according to the position of the (2D) projection of the cell-type-specific cell state. This is helpful to prioritize which cell type may have spatially consistent variations in cell states. To relate this to the functions of each axis of the PCs, we filter genes that are the most correlated with each PC (Pearson correlation) and we ran gene set enrichment analysis using EnrichR [36,37] for the top 50 genes.

### A post-hoc recipe for differential expression with DestVI

Performing differential expression with a probabilistic model is always challenging, but is crucial for making robust scientific discoveries. In our previous work, we used a purely Bayesian approach to differential expression [55,56]. Because we apply MAP inference (and not fully-Bayesian inference) to DestVI, we instead developed a hybrid approach to differential expression, where we sample from the adequate generative distribution and then derive a p-value with a frequentist test. More precisely, for two bags of spots (*x*_*a*_)_*a*∈*B*_ and (*x*_*b*_)_*b*∈*B*_, and a cell type *c*, we operate as follows.

#### Characterizing cell-type-specific gene expression

We query the parameters of the generative distribution for that spot, but only for the contribution from cell type *c*. We do this by embedding all the spots, and keeping only the latent variable for the cell type of interest 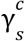 and then querying the decoder network 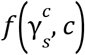 for this particular 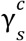 and cell type *c*. This vector, along with *p*_*g*_, fully characterize the transcriptome of a fictitious cell *x*_*sc*_ of type *c* in spot *s*:

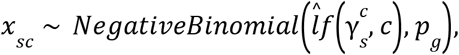

where 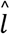 is a fixed scalar equal to the average library size in the single-cell data.

#### Simulating cell-type-specific gene expression

Then, we sample multiple times per spot from the mean of the Poisson distribution underlying the negative binomial:

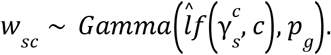

There are several considerations for sampling this way. First, as we underlined in our previous work for calculating gene correlations in totalVI, using the mean of the generative distribution provides all forms of biases. Second, sampling from the full negative binomial distribution introduces Poisson noise, and reduces sensitivity of the method. Sampling from the Gamma distribution is an intermediate, although post-hoc, solution. This step plays the role of denoising and is especially important when there are very few spots in the bags *A* or *B*, because we can generate more data points.

#### Hypothesis testing

For every cell in every bag, we can generate between 10 and 100 independent samples of the random variables *w*_*sc*_ and apply a two-sample Kolmogorov–Smirnov test, and we correct for multiple testing using the Benjamini-Hochberg procedure [71]. Additionally, we tag genes as differentially expressed if two conditions are met: (i) the null hypothesis is rejected, after control of the FDR at level 0.05, and (ii) the log-fold change (LFC) is greater than a data-driven threshold, in absolute value. To calculate this threshold, we assume that a significant amount of LFCs will concentrate around zero (which correspond to the genes that are equally expressed), while DE genes will concentrate around other modes. Based on this assumption, we fit a three-components Gaussian Mixture Model (GMM) to the LFC, and keep the mean of the mode with largest absolute value, whose associated distributions should contain differentially expressed genes.

### Experimental conditions and sample preparation

#### Mice

C57BL/6J mice were purchased from Harlan and housed in the Weizmann Institute of Science animal facility, under specific pathogen-free conditions. YARG mice (B6.129S4-Arg1tm1Lky/J, Jax015857) are bought from Jax Laboratory. For consistency, female mice, 6-8 weeks of age, were used for all experiments. All mice were provided with normal chow and water ad libitum, and housed under a strict 12-hour light-dark cycle. All experimental protocols were approved by the Institutional Animal Care and Use Committee (IACUC).

#### Preparation of antigens

Inactivated Mycobacterium Smegmatis (MS) was prepared as previously reported in [24]. Simply M. smegmatis was grown in Luria-Bertani broth overnight at 37°C. Bacteria were washed thrice in PBS containing 0.05% Tween 80 and heat-killed at 75°C for 1 hour, then aliquoted into a useful size and stored at −80°C.

#### Immunizations

Mice were anesthetized and M. smegmatis was administered by intradermal injection into the ear pinna. PBS was injected into the ear pinna of control animals. The pathogen immunization dose is 4M CFU, according to our previous study [24]. The auricular lymph nodes were harvested 48 hours after immunization.

#### Tumor model

MCA205 fibrosarcoma cell lines were kindly provided by Sergio Quezada group at UCL cancer institute, London, UK. Cells were cultured in DMEM (41965-039) medium supplemented with 10% heat-inactivated FBS, 1mM sodium pyruvate, 2mM l-glutamine, 1% penicillin-streptomycin (Thermo Fisher Scientific). Cells were cultured in 100 mm tissue culture plates in an incubator with humidified air and 5% CO2 at 37°C. For establishment of solid tumors, 8 weeks, female mice were injected intradermally with 5 × 10^5^ MCA205 tumor cells suspended in 100 μL PBS on their right flank. On day 14, tumor volume was measured using a caliper and prepared according to Visium spatial protocols of tissue preparation guide (CG000240).

### RNA-sequencing and data processing

#### Single-cell RNA-Sequencing for the lymph node

To prepare single cell suspensions for scRNA-seq using 10x Genomics system, auricular LNs were digested in IMDM containing 100 μg/mL Liberase TL and 100 μg/mL DNase I (both from Roche, Germany) for 20 minutes at 37°C. Last 5 minutes of incubation, EDTA was added at a final concentration of 10 mM. Cells were collected, filtered through a 70 μm cell strainer, washed with PBS + 0.04% BSA for a final concentration of 1000 cells/μL.

Cellular suspension was immediately loaded onto Next GEM Chip G targeting ~5000 cells and then ran on a Chromium controller instrument (10x Genomics) to generate GEM emulsion. Single cell 3’ RNA-seq libraries were generated according to the manufacturer’s protocol using the v3.1 Next GEM dual index workflow. Final libraries were quantified using NEBNext Library Quant Kit for Illumina (NEB) and high sensitivity D1000 TaepStation (Agilent). Libraries were pooled and sequenced on an SP 100 cycles reagent kit on a NovaSeq6000 instrument (Illumina), aiming for ~ 50,000 reads per cell. Reads from raw FASTQ files were processed with Cell Ranger 4.0 and mapped to the mouse mm10–2020 reference (10x Genomics). No read depth normalization was applied when aggregating datasets.

#### Lymph node scRNA-seq data processing

Using Scanpy [67], we filtered out cells with less than 200 UMIs and genes expressed in less than 3 cells, as well as cells with more than 20% mitochondrial genes. We estimated doublet scores for all barcodes using Scrublet [72]. Because we did not expect any novel cell types in the data, we seeked to automatically annotate our samples based on publicly available murine lymph node scRNA-seq [38]. We therefore harmonized all the samples from both datasets using scVI [29], and transferred labels using scANVI [39]. For both algorithms, we filtered genes to match the highly variable genes from the dataset of [38]. After manual examination of the expression of marker genes (Supplementary Table 4 of [38]) and its adequation with the automated labels, we noticed some mislabeling inaccuracies on the rare cell types of the myeloid cluster. Consequently, we clustered the myeloid compartment with Louvain and curated the annotation, based not only on the automated annotation, but also the gene expression of the key marker genes. Finally, we also removed a cluster of doublets, as predicted by Scrublet. We recapitulate the number of cells for each cell type in **Supplementary Table 1**.

#### Single-cell RNA-Sequencing for the tumor

To prepare tumor infiltrating leukocytes single cell suspensions, the tumors underwent mechanical (gentle-MACSTM C tube, Miltenyi Biotec Inc., San Diego, CA) and enzymatic digestion (0.1mg/ml DNase type I (Roche), and 1mg/ml Collagenase IV (Worthington) in RPMI-1640) for 10 minutes at 37°C and repeat one more time. Cells then filtered through 100 μm cell strainer, washed with an ice cold sorting buffer, centrifuged (5 min, 4°C, 350g), and stained with fluorophores conjugated anti-mouse CD45 antibodies on ice 30 minutes avoid light. After staining, cells were washed and resuspended in a cold washing buffer (0.5% BSA and 2 mM EDTA in PBS), filtered through a 70 μm cell strainer. Before sorting, cells were stained with propidium iodide to exclude dead/dying cells. Cell sorting was performed using a BD FACSAria Fusion flow cytometer (BD Biosciences), gating for CD45+ cells after exclusion of dead cells and doublets (**Supplementary Figure 20**). Single cells were sorted into 384-well plates and single cell RNA-seq libraries were generated using a modified version of the single cell MARS-seq protocol [73] [74]. In brief, mRNA from cells sorted into cell capture plates were barcoded, converted into cDNA and pooled using an automated pipeline. The pooled cDNA are then amplified and Illumina libraries are being generated. Final libraries were quantified using Qubit and high sensitivity D1000 TapeStation (Agilent). Libraries were pooled and sequenced an SP 100 cycles reagent kit on a NovaSeq6000 instrument (Illumina). Sequences were mapped to the mouse (mm10). Demultiplexing and filtering was performed as previously described [74], with the following adaptations: Mapping of reads was performed using HISAT (version 0.1.6); reads with multiple mapping positions were excluded. Reads were associated with genes if they were mapped to an exon, using the ensembl gene annotation database (embl release 90). Exons of different genes that shared a genomic position on the same strand were considered as a single gene with a concatenated gene symbol. The level of spurious unique molecular identifiers (UMIs) in the data were estimated by using statistics on empty MARS-seq wells, and excluded rare cases with estimated noise > 5% (median estimated noise over all experiments was 2%).

#### Tumor scRNA-seq data processing

Using Scanpy [67], we filtered out cells with less than 200 UMIs and genes expressed in less than 10 cells. We selected 4,000 highly variable genes using scanpy and reduced dimensionality of the data using scVI [29]. We applied leiden clustering and annotated the data based on marker genes (CD4 and Icos for CD4 T cells, Cd8a and Cd8b1 for CD8 T cells, Gzma and Prf1 for NK cells, C1qa and Ly6a for macrophages / monocytes, S100a8 for Neutrophils, H2-Ab1, H2-Eb1 and H2-Aa for DCs and Cd63 and Col3a1 for tumor cells. We recapitulate the number of cells for each cell type in **Supplementary Table 5**.

#### Visium data generation

Auricular LNs and the MCA205 tumor were prepared according to Visium spatial protocols of tissue preparation guide (10x genomics). Firstly, freshly obtained tissue samples were snap frozen in liquid nitrogen, then embedded in chilled Optimal Cutting Temperature compound (OCT; Tissue-Tek) and frozen on dry ice, then stored at −80°C in a sealed container for later use. For Visium samples preparation, OCT-embedded tissue blocks were cut to 10 μm thick using a LEICA CM1950 machine and mounted on the Visium spatial gene expression slide. For gene expression samples, tissues were permeabilized for 18 minutes, based on tissue optimization time course experiments. Brightfield histology images were taken using a 10X objective (Plan APO, NA 0.25) on Leica DMI8 wide-field inverted microscope according to Visium spatial gene expression imaging guidelines (CG000241). Raw images were stitched together using Leica application suite X (LAS X) software and exported as TIFF/PNG files with low- and high-resolution settings.

Libraries were prepared according to the Visium spatial gene expression user guide (10x genomics). Final libraries were quantified using NEBNext Library Quant Kit for Illumina (NEB) and high sensitivity D1000 TapeStation (Agilent). The number of reads required for sequencing was calculated taking into account the percentage of the tissue within each capture area (calculated using imageJ). Libraries were pooled according to the desired number of reads and sequenced on an SP 200 cycles reagent kit on a NovaSeq6000 instrument (Illumina).

#### Visium raw data processing

Raw FASTQ files and histology images were processed by sample with the Space Ranger software v1.2.1. For the lymph node, we calculated the quality control metrics using Scanpy [67] and noticed that one of the PBS samples was of low quality, as indicated by the number of UMIs per spot. We also filtered out spots with less than 4,000 UMIs and genes expressed in less than 10 spots. The number of spots after filtering was 1,092 across the remaining three lymph nodes. For the tumor data, we filtered out spots with more than 2% mitochondrial gene expression, spots with less than 10,000 UMIs and genes expressed in less than 10 spots.

### Immunofluorescence

Tissues embedded in OCT for Visium were sliced to 10 μm thick sections using a LEICA CM1950 machine and mounted on SuperFrost Plus slides (Thermo Scientific). For visualization, sections were firstly fixed by 4% Formaldehyde solution in PBS diluted from 16% Formaldehyde (Thermo Scientific) 10 minutes at room temperature. Then sections were washed by PBS three times and blocked with a blocking buffer solution (5% donkey serum, 2% BSA, 0.2% Triton X-100) for 2 h at room temperature, and incubated with primary antibody at 4°C overnight. If secondary antibody is necessary, after three times PBST (0.02% Triton X-100) washes corresponding secondary antibody was incubated at room temperature 1 hour. After three times PBST washed, DAPI (4’,6-Diamidino-2-Phenylindole, Dilactate, Biolegend) reagent was added for 10 min to detect cell nuclei. Sections were mounted with SlowFade Gold Antifade Mountant (Invitrogen, S36937) and sealed with cover-slips. Microscopic analysis was performed using a laser-scanning confocal microscope (Zeiss, LSM880). Images were acquired and processed with the same threshold settings using Imaris software (Bitplane). The primary antibodies used were: CD45 APC (1:100, 30-F11, eBioscience, 17-0451-82), CD11b Biotin (1:100, M1/70, Biolegend, 101204), CD11b PE (1:100, M1/70, eBioscience, 12-0112-83), CD64 PE (1:100, X54-5/7.1, Biolegend, 139303), Ly6C FITC (1:100, HK1.4, Biolegend, 128005), B220 (1:100, RA3-6B2, Biolegend, 103208), CD3 Biotin (1:100, 17A2, Biolegend, 100243), TCRb PE (1:100, H57-597, Biolegend, 109207), MHCII I-A/I-E FITC (1:100, M5/114.15.2, Biolegend, 107606), F4/80 APC (1:100, BM8, eBioscience, 17-4801-82), NK1.1 PE(1:100, PK136, eBioscience, 12-5941-63), CD31 APC (1:100, MEC13.3, Biolegend, 102509), IFIT3 polyclonal antibody (1:500, Proteintech, 15201-1-AP). Secondary antibody used were: Streptavidin APC (1:400, Biolegend, 405207), Goat anti-Rabbit IgG-heavy and light chain Antibody DyLight® 650 Conjugated (1:800, Bethyl, A120-101D5).

## Data Availability

The raw data discussed in this manuscript have been deposited in the National Center for Biotechnology Information’s Gene Expression Omnibus and are accessible through accession number GSE173778 (murine lymph node and tumor; spatial transcriptomics and scRNA-seq data). Processed data are available on our reproducibility repository (https://github.com/romain-lopez/DestVI-reproducibility).

## Software Availability

The code to reproduce the results in this manuscript is available on the Github repository https://github.com/romain-lopez/DestVI-reproducibility and has been deposited to Zenodo https://doi.org/10.5281/zenodo.4685952. The reference implementation of DestVI, along with accompanying tutorials is available via the scvi-tools codebase at https://scvi-tools.org/.

## Author Contributions

R.L, B.L, H.K-S, I.A and N.Y designed the study and the experiments. B.L performed the experimental procedures. H.K-S, M.K and D.P prepared Visium and scRNA-seq libraries. A.J processed single-cell RNA sequencing of the tumor data. B.L and Y.A contributed to microscopy analysis. E.D. assisted with RNA-sequencing data processing and data upload. R.L conceived the statistical model with input from B.L, H.K-S and M.I.J. R.L implemented the DestVI software and applied the software to analyze the data, with input from A.W. P.B proposed the spatially-aware extension of DestVI. I.A and N.Y supervised the work.

## Acknowledgements

We would like to acknowledge Adam Gayoso and Galen Xing for their help integrating DestVI in the scvi-tools codebase. Thanks to Zoë Steier for providing guidance on the annotation of the lymph node single-cell data. We thank Efrat Davidson for the artwork. We are grateful for insightful conversations with Assaf Weiner, Aviv Regev, Dana Pe’er, Quaid Morris, Alexis Battle, Elior Rahmani and Matthew Jones.

## Funding

NY and RL were supported by the Chan Zuckerberg Biohub. I.A. is an Eden and Steven Romick Professorial Chair, supported by Merck KGaA, Darmstadt, Germany, the Chan Zuckerberg Initiative, the Howard Hughes Medical Institute International Scholar award, the European Research Council Consolidator Grant 724471-HemTree2.0, an SCA award of the Wolfson Foundation and Family Charitable Trust, the Thompson Family Foundation, a Melanoma Research Alliance Established Investigator Award (509044), the Israel Science Foundation (703/15), the Ernest and Bonnie Beutler Research Program for Excellence in Genomic Medicine, the Helen and Martin Kimmel award for innovative investigation, the NeuroMac DFG/Transregional Collaborative Research Center Grant, an International Progressive MS Alliance/NMSS PA-1604 08459, the ISF Israel Precision Medicine Program (IPMP) 607/20 grant and an Adelis Foundation grant.

## Competing Interests Statement

N.Y. is an advisor and/or has equity in Cellarity, Celsius Therapeutics, and Rheos Medicine.

