## Supplementary Information for "Multi-resolution deconvolution of spatial transcriptomics data reveals continuous patterns of inflammation"

### Supplementary Figures

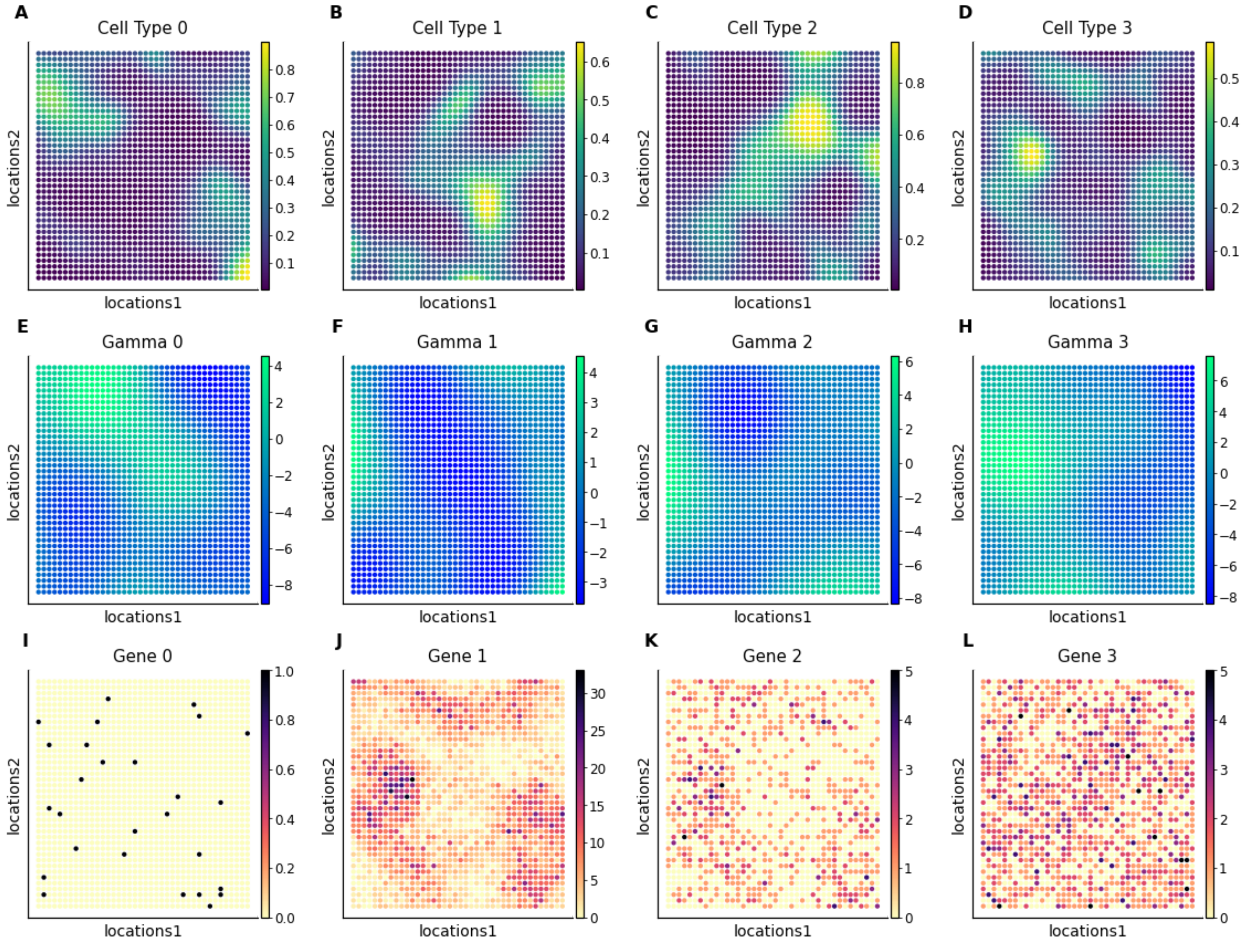


**Supplementary Figure 1.** Presentation of the simulated spatial data (each point is a spot, placed according to its spatial coordinates). **(A-D)** Simulated cell-type proportions (four out of the five cell types). **(E-H)** Simulated cell-type-specific cell states (the four dimensions for one cell type) **(I-L)** Simulated gene expression at each spot (four out of the two thousand genes).

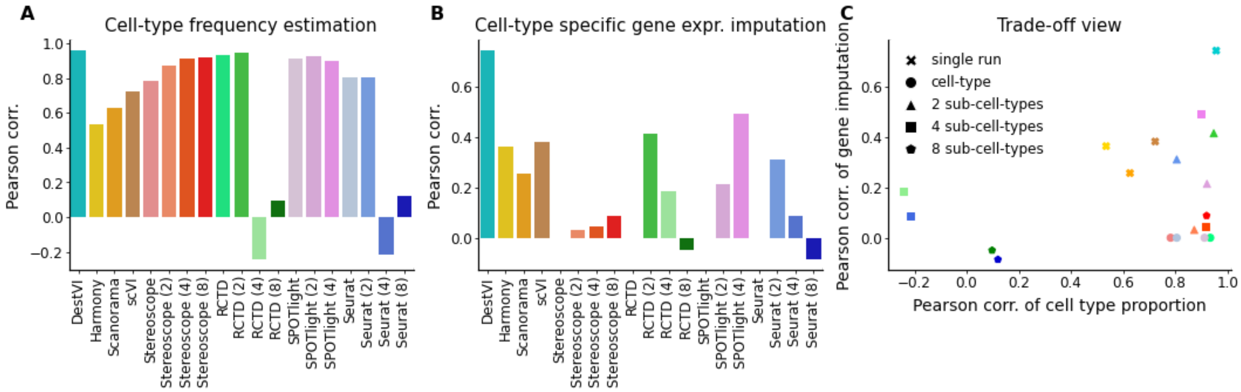


**Supplementary Figure 2.** Comparison of DestVI to competing algorithms, possibly applied to different clustering resolutions. **(A)** Pearson correlation of estimated cell-type proportions compared to ground truth for several algorithms. **(B)** Pearson correlation of estimated cell-type-specific gene expression compared to ground truth for several algorithms **(C)** Scatter plot of both metrics.

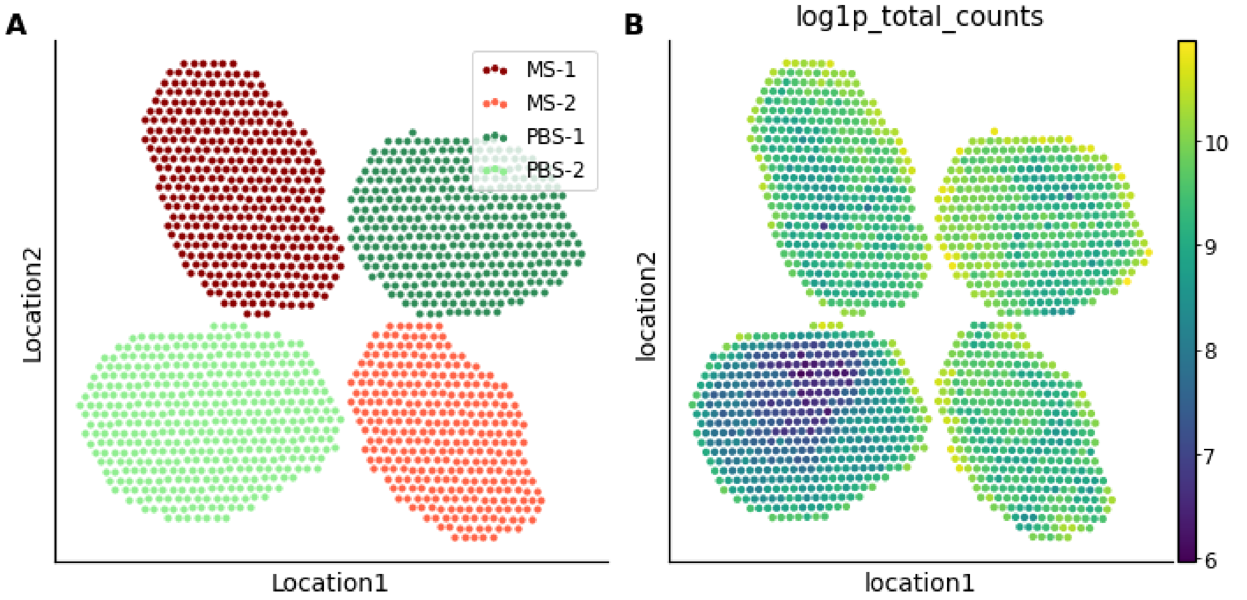


**Supplementary Figure 3.** Quality control metrics in the lymph node spatial transcriptomics data. **(A)** The four lymph node slices initially processed with 10x Visium. **(B)** The log library size as processed by scanpy on all lymph nodes, displayed based on their spatial location. The PBS-2 lymph node sample has a significantly lower number of detected UMIs per spot (magnitude of exp(4); 55x less).

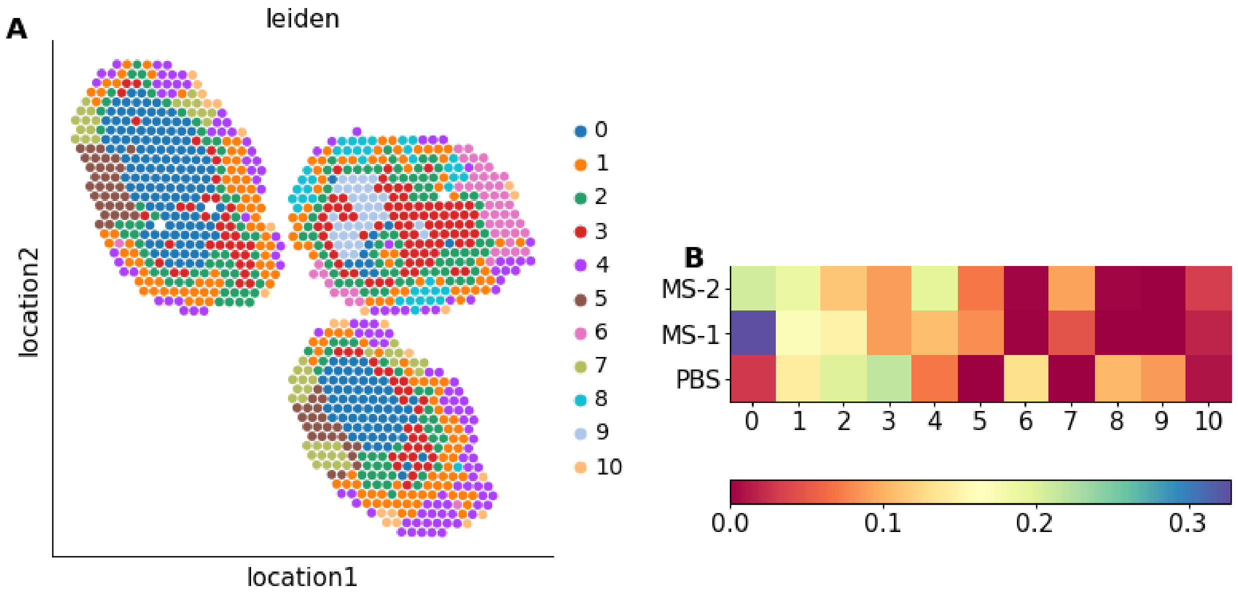


**Supplementary Figure 4.** Simple comparative analysis of the spatial lymph node data based on scanpy. **(A)** Clustering analysis: we normalized the spot counts, and proceeded to leiden clustering after principal component analysis. **(B)** Comparison of the proportion of clusters abundance across spots of the different lymph nodes.

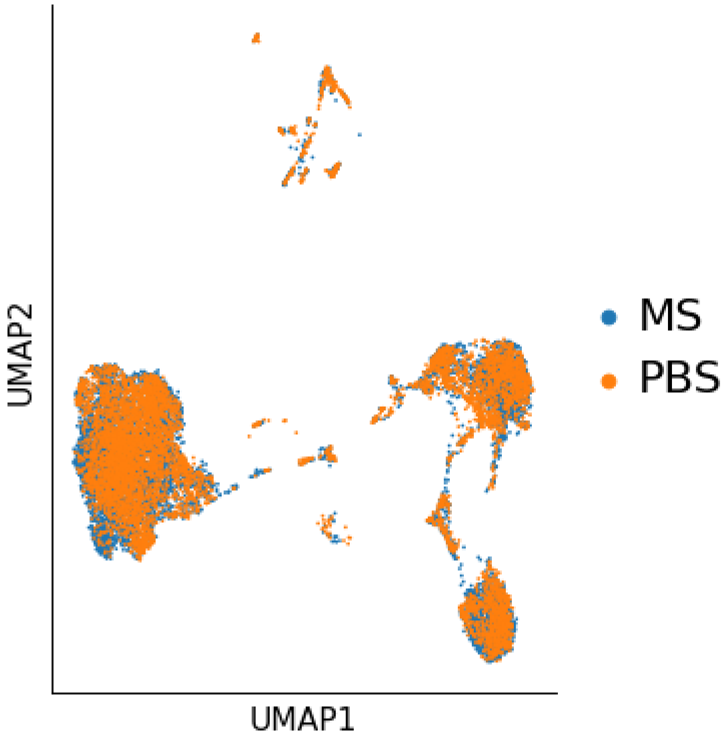


**Supplementary Figure 5.** Two-dimensional projection by UMAP, as embedded by scVI of the lymph node single-cell RNA sequencing data.

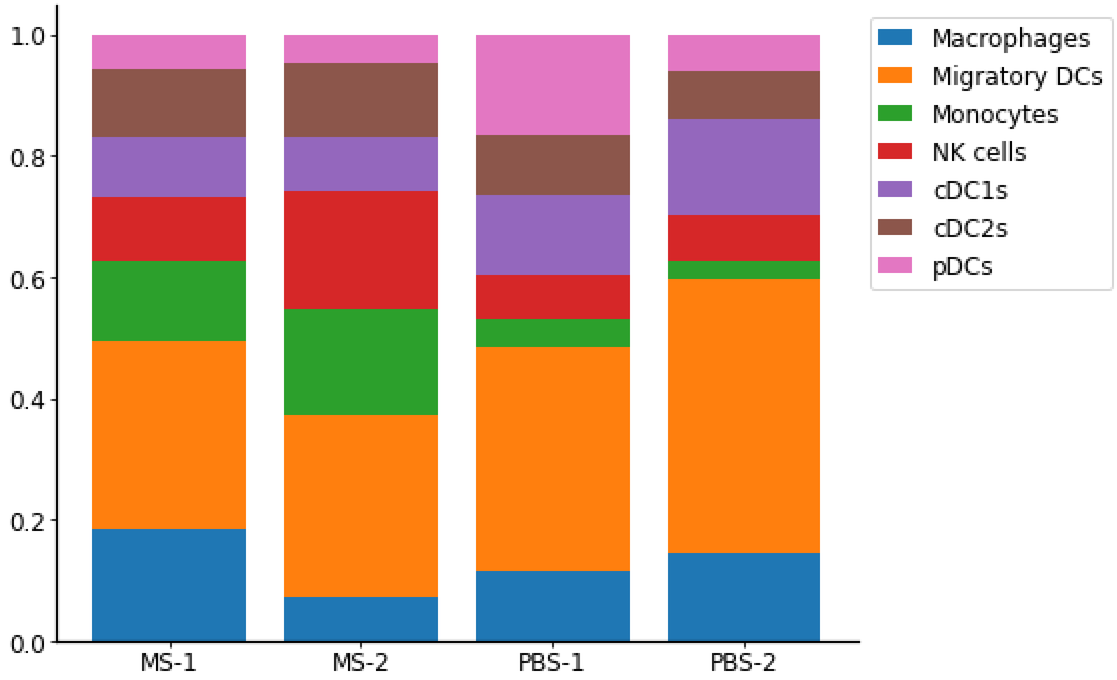


**Supplementary Figure 6.** Summary statistics of the cell type proportion for the annotated lymph node single-cell RNA sequencing data, for each lymph node.

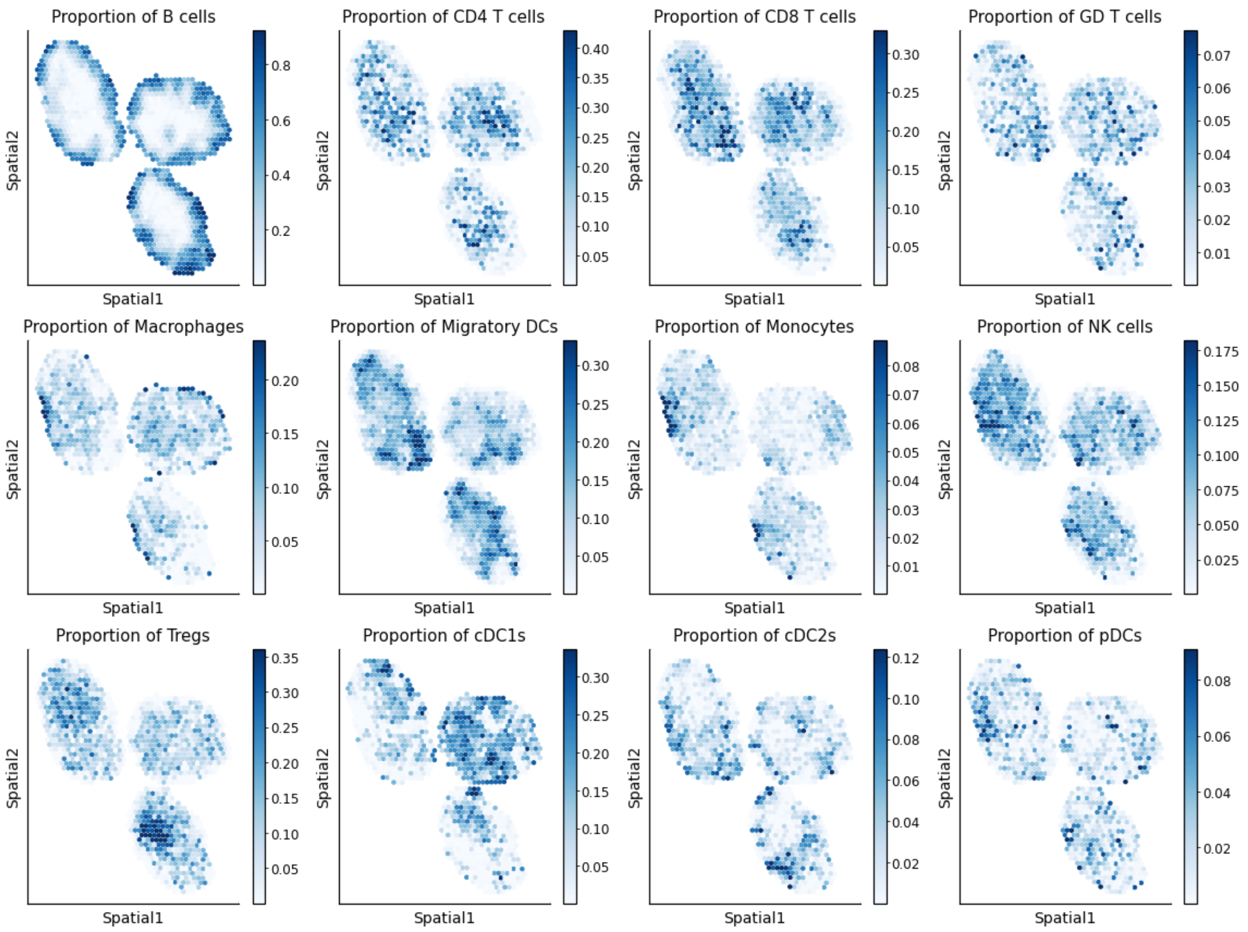


**Supplementary Figure 7.** Abundance of all cell types across all spots for the lymph node spatial transcriptomics data.

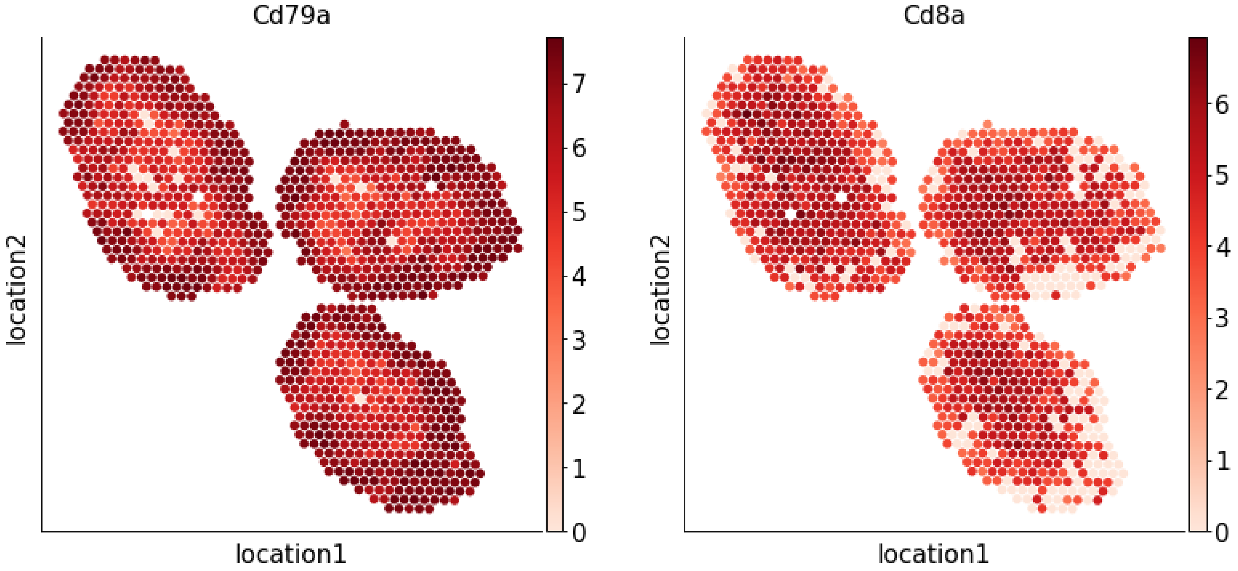


**Supplementary Figure 8.** Expression of key marker genes for the definition of the T cell compartment and the B cell area.

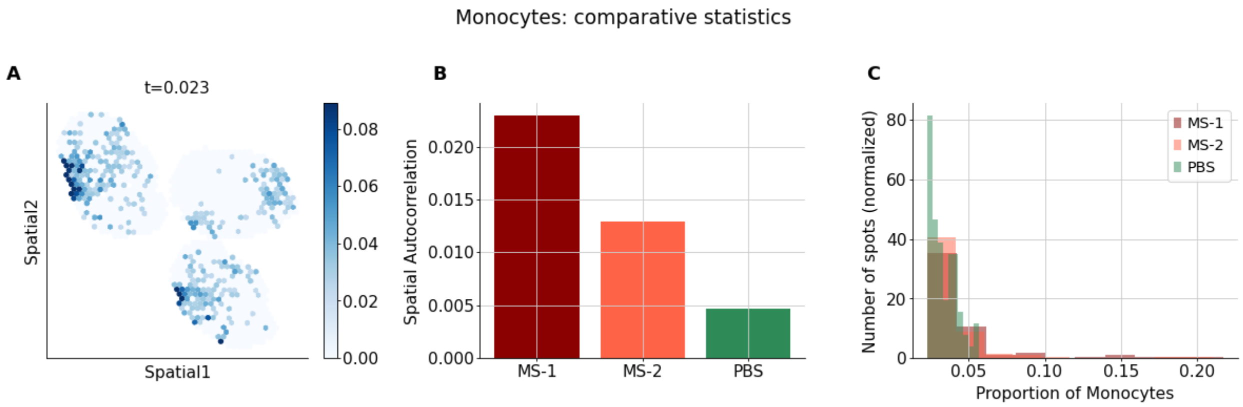


**Supplementary Figure 9.** Follow up analyses for the proportion of monocytes. **(A)** spatial distribution for the proportion of cell type, thresholded according to the characteristic point method (**Online Methods**) **(B)** Autocorrelation metric of the cell-type proportion per lymph node (Geary C) **(C)** histogram of thresholded proportions across all spots, per lymph node.

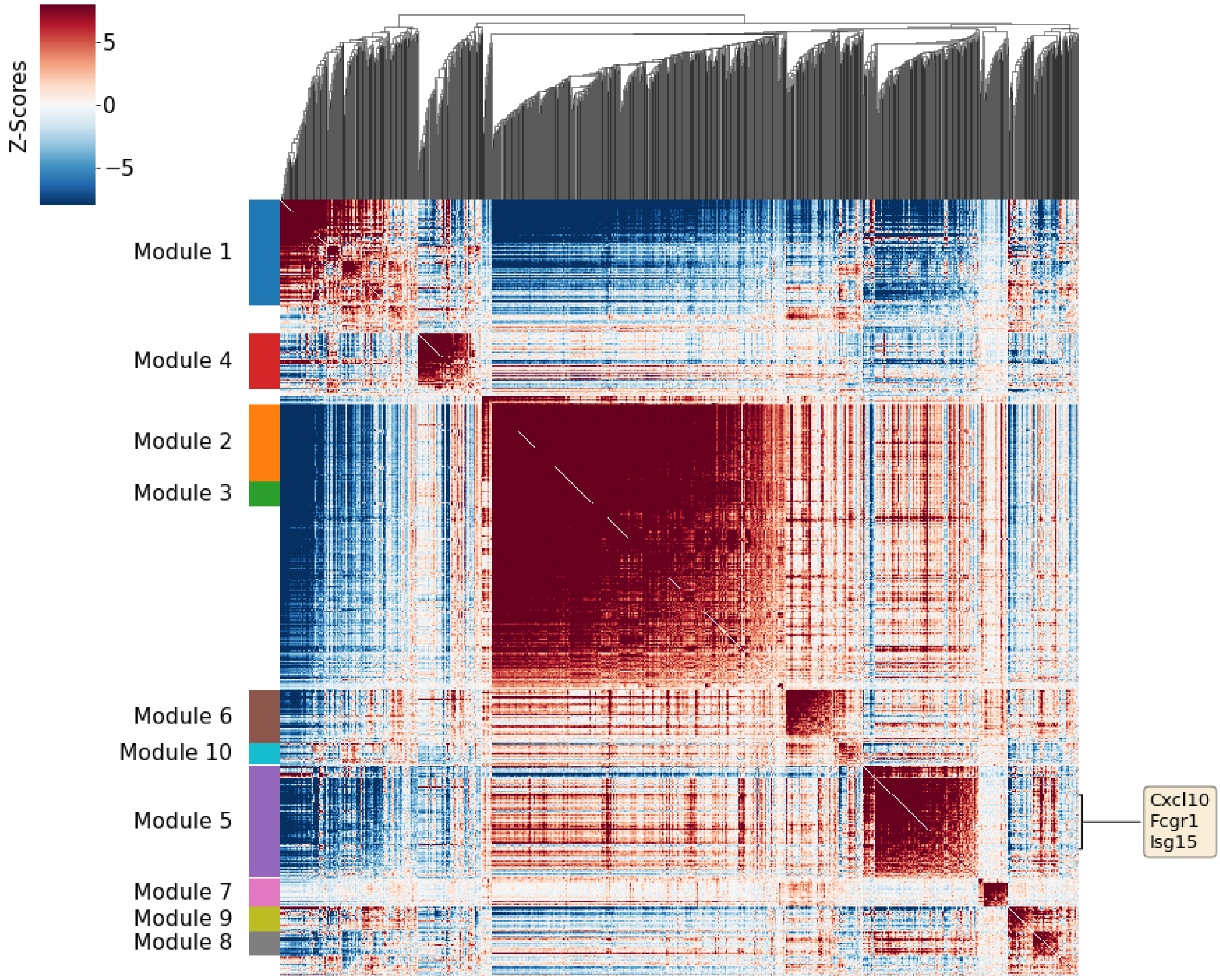


**Supplementary Figure 10.** Hotspot analysis for the monocytes: gene-gene local autocorrelation matrix, and clustering into gene modules, based on the scRNA-seq data.

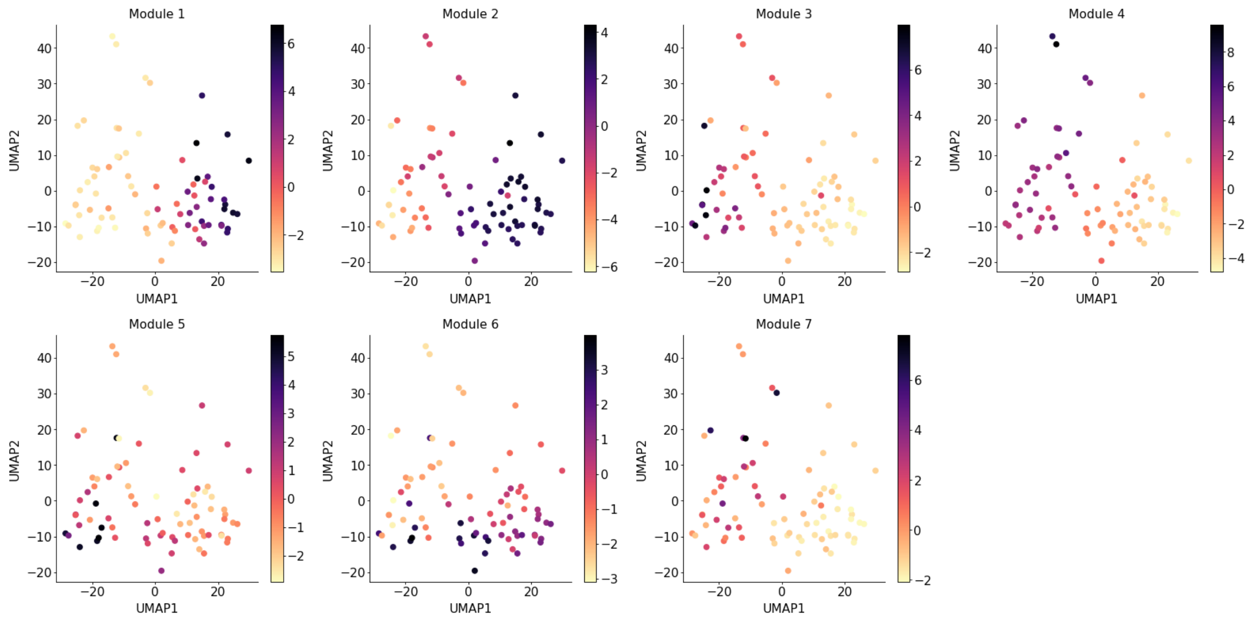


**Supplementary Figure 11.** Hotspot analysis for the monocytes: projection of every gene module onto the embedding of the single-cell data (scLVM).

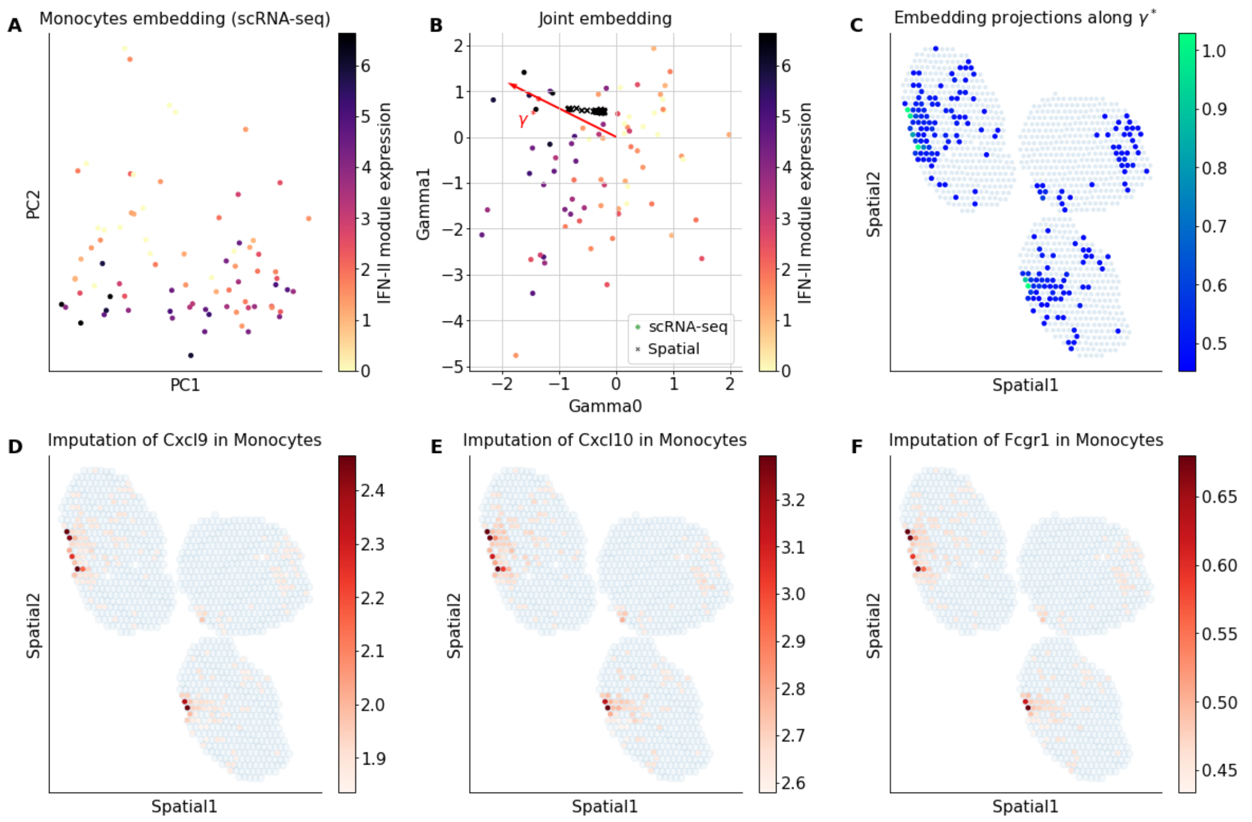


**Supplementary Figure 12.** Follow up analyses for the monocytes **(A)** Embedding of the monocytes from single-cell data, colored by expression of the selected IFN-II genes identified by Hotspot (Fcgr1, Cxcl9 and Cxcl10). **(B)** Joint embedding of the monocytes from the scRNA-seq data (circles) and the spots with high abundance of monocytes from the spatial transcriptomics data (crosses). Single-cell data is colored by normalized expression of the IFN-II marker genes. The arrow designates an interesting direction $\gamma^*$, drawn by eye-sight. **(C)** Spatial data colored by the projection of the embedding from DestVI for every spot along the direction $\gamma^*$. **(D-F)** Imputation of monocyte-specific expression of several marker genes for the spatial data

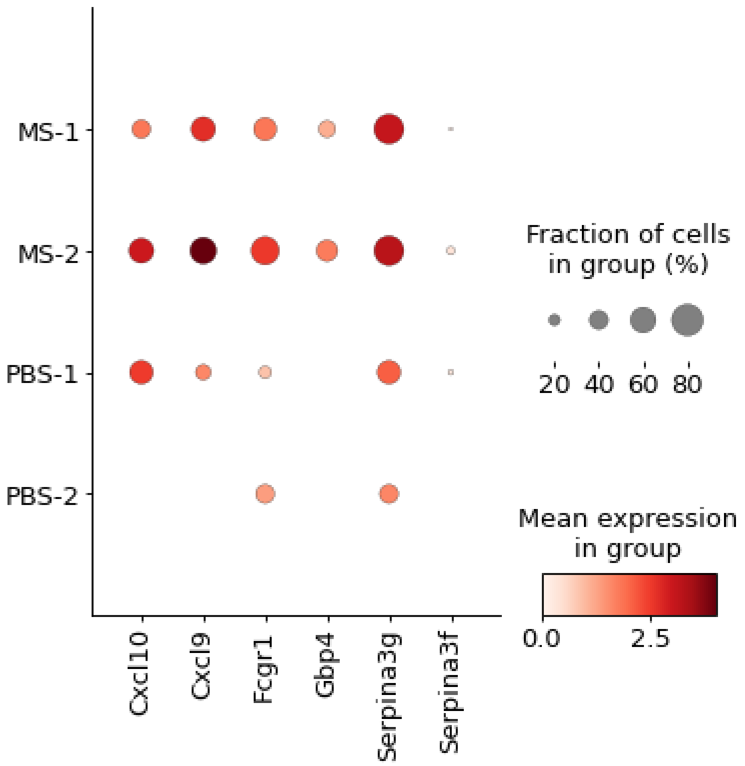


**Supplementary Figure 13.** Comparative analysis of gene expression for the monocytes of the scRNA-seq data.

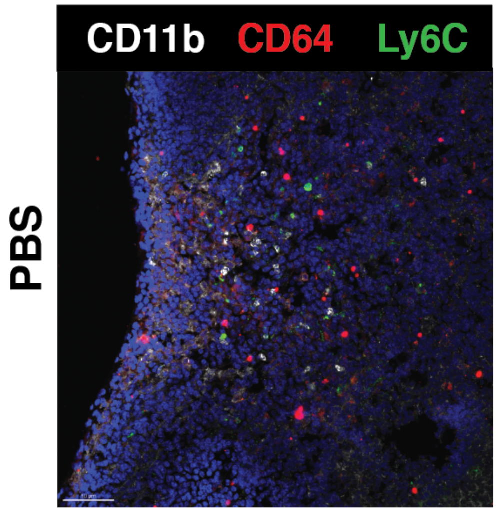


**Supplementary Figure 14.** Immunofluorescence imaging from a PBS lymph node. Scale bar, 50 µm.

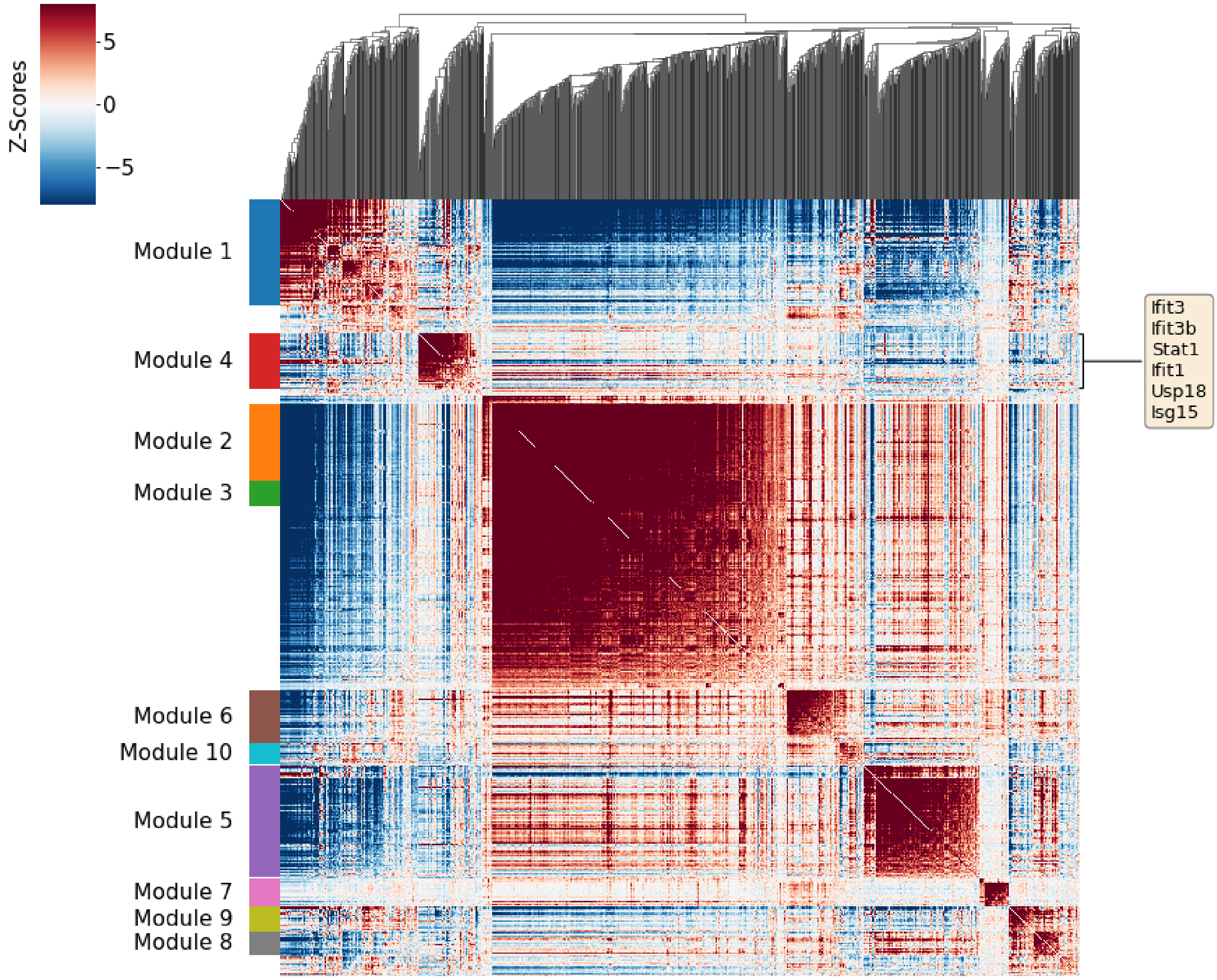


**Supplementary Figure 15.** Hotspot analysis for the B cells: gene-gene local autocorrelation matrix, and clustering into gene modules, based on the scRNA-seq data.

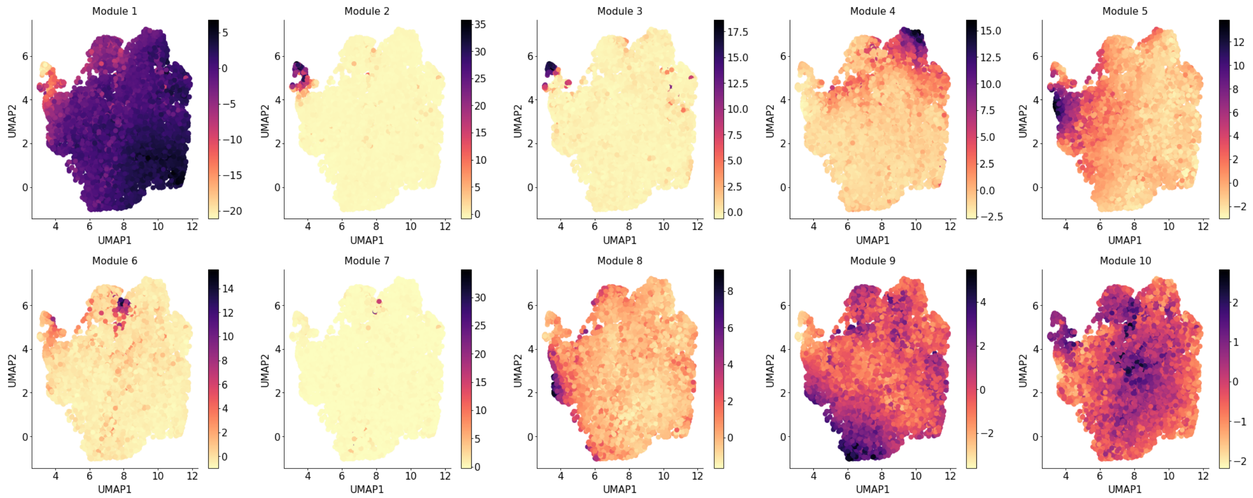


**Supplementary Figure 16.** Hotspot analysis for the B cells: projection of every gene module onto the embedding of the single-cell data (scLVM).

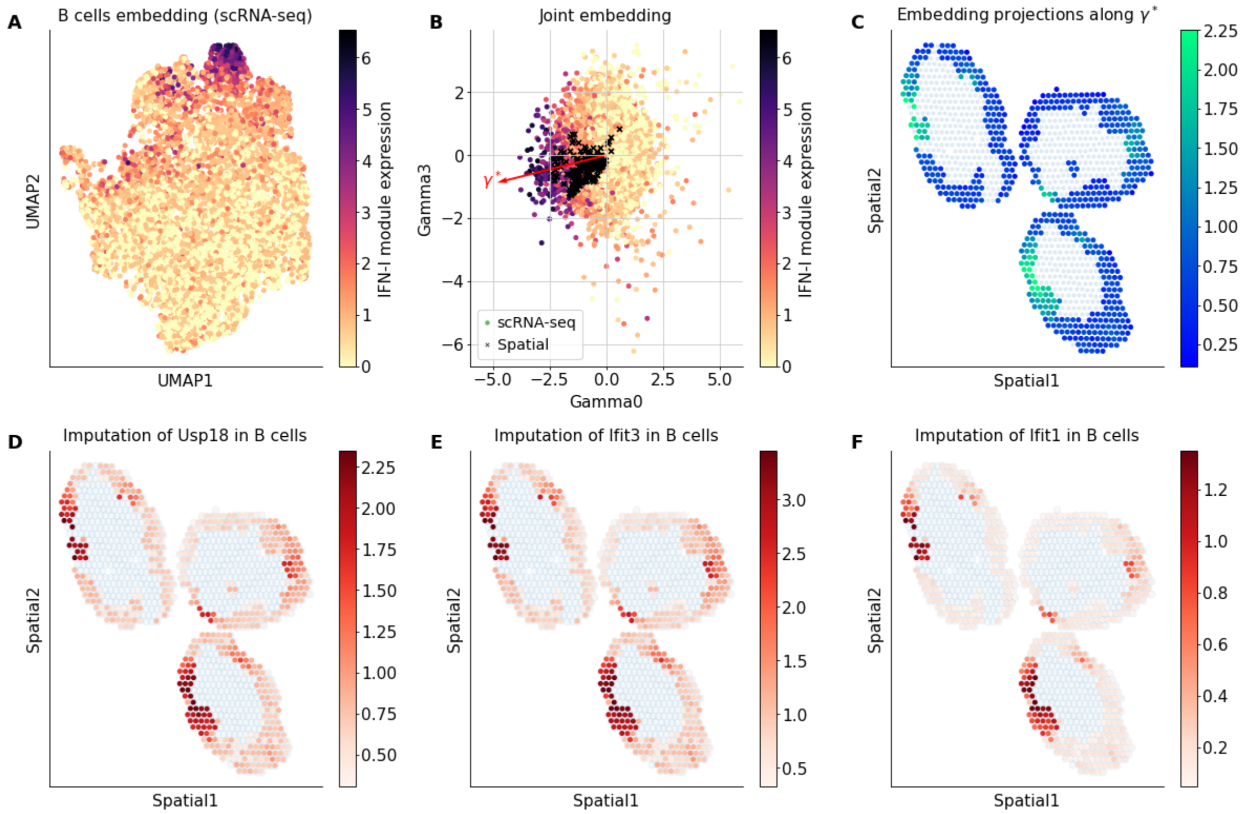


**Supplementary Figure 17.** Follow up analyses for the B cells. **(A)** Embedding of the B cells from single-cell data, colored by expression of the selected IFN-I genes identified by Hotspot (Ifit3, Ifit3b, Stat1, Ifit1, Usp18 and Isg15). **(B)** Joint embedding of the B cells from the scRNA-seq data (circles) and the spots with high abundance of B cells from the spatial transcriptomics data (crosses). Single-cell data is colored by normalized expression of the IFN-II marker genes. The arrow designates an interesting direction $\gamma^*$, drawn by eye-sight. **(C)** Spatial data colored by the projection of the embedding from DestVI for every spot along the direction $\gamma^*$. **(D-F)** Imputation of B cell-specific expression of the several marker genes on the spatial data.

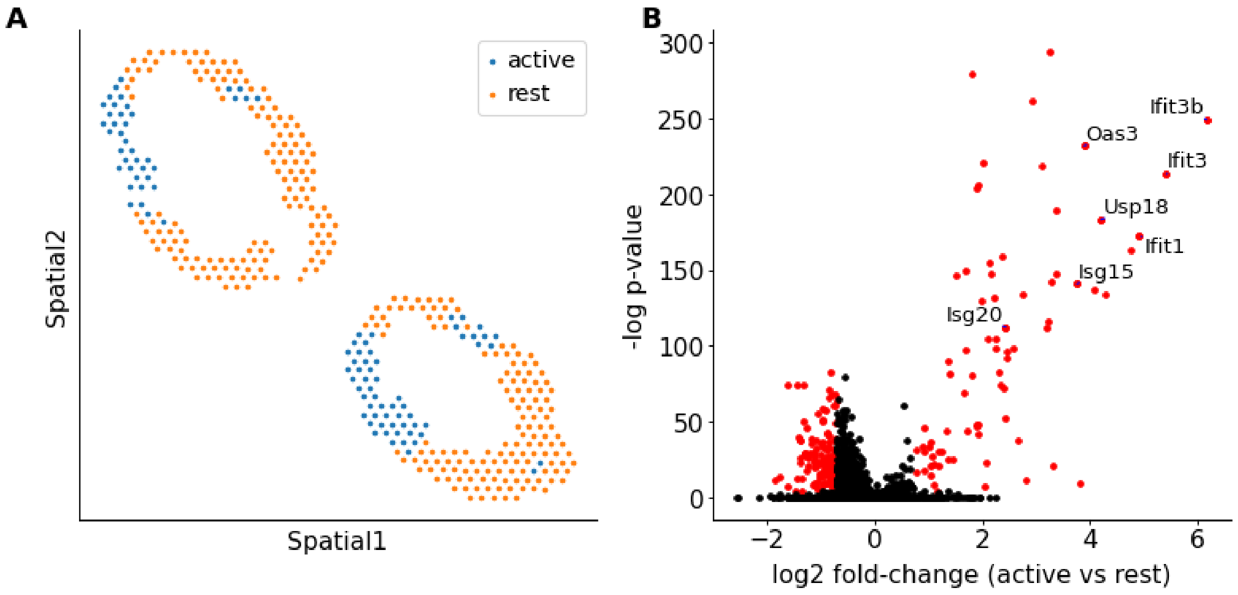


**Supplementary Figure 18.** **(A)** Definition of the active zone for differential expression analysis within MS samples. **(B)** B cell-specific differential expression analysis between MS and PBS lymph nodes (2,000 genes). Significance is calculated with our differential expression procedure (**Online Methods**).

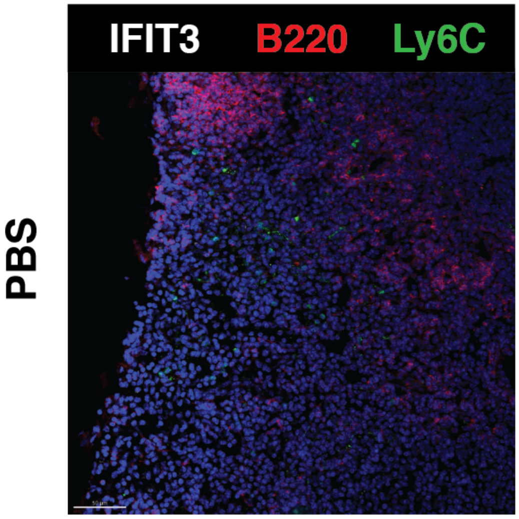


**Supplementary Figure 19.** Immunofluorescence imaging from a PBS lymph node. Scale bar, 50 µm.

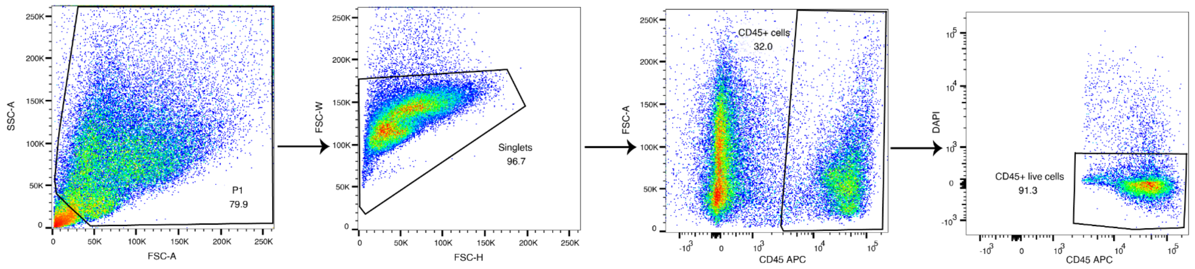


**Supplementary Figure 20.** FACS information for tumor single-cell data processing. Representative flow cytometry analysis of MCA205 tumor, showing CD45+ live cell fractions.

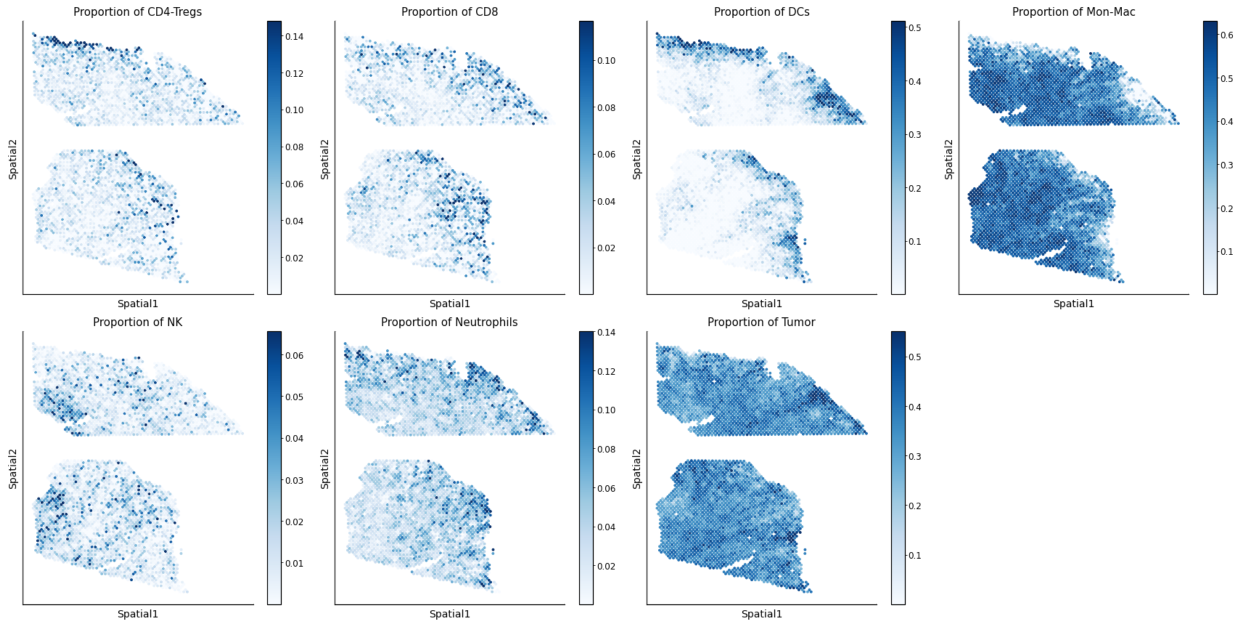


**Supplementary Figure 21.** Abundance of all cell types across all spots for the tumor spatial transcriptomics data.

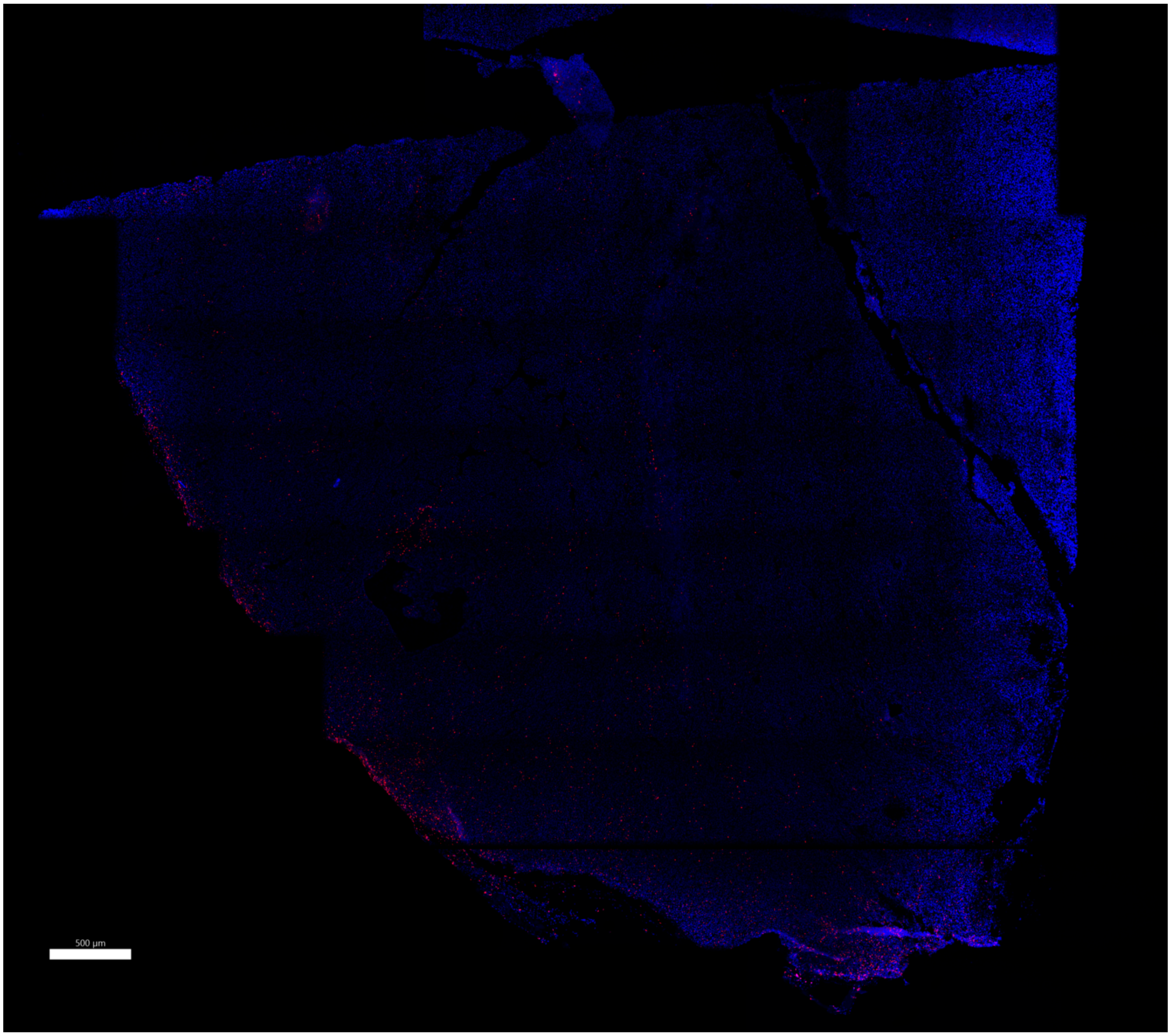


**Supplementary Figure 22.** Immunofluorescence imaging from the tumor using antibodies for TCRb. Scale bar, 500 µm.

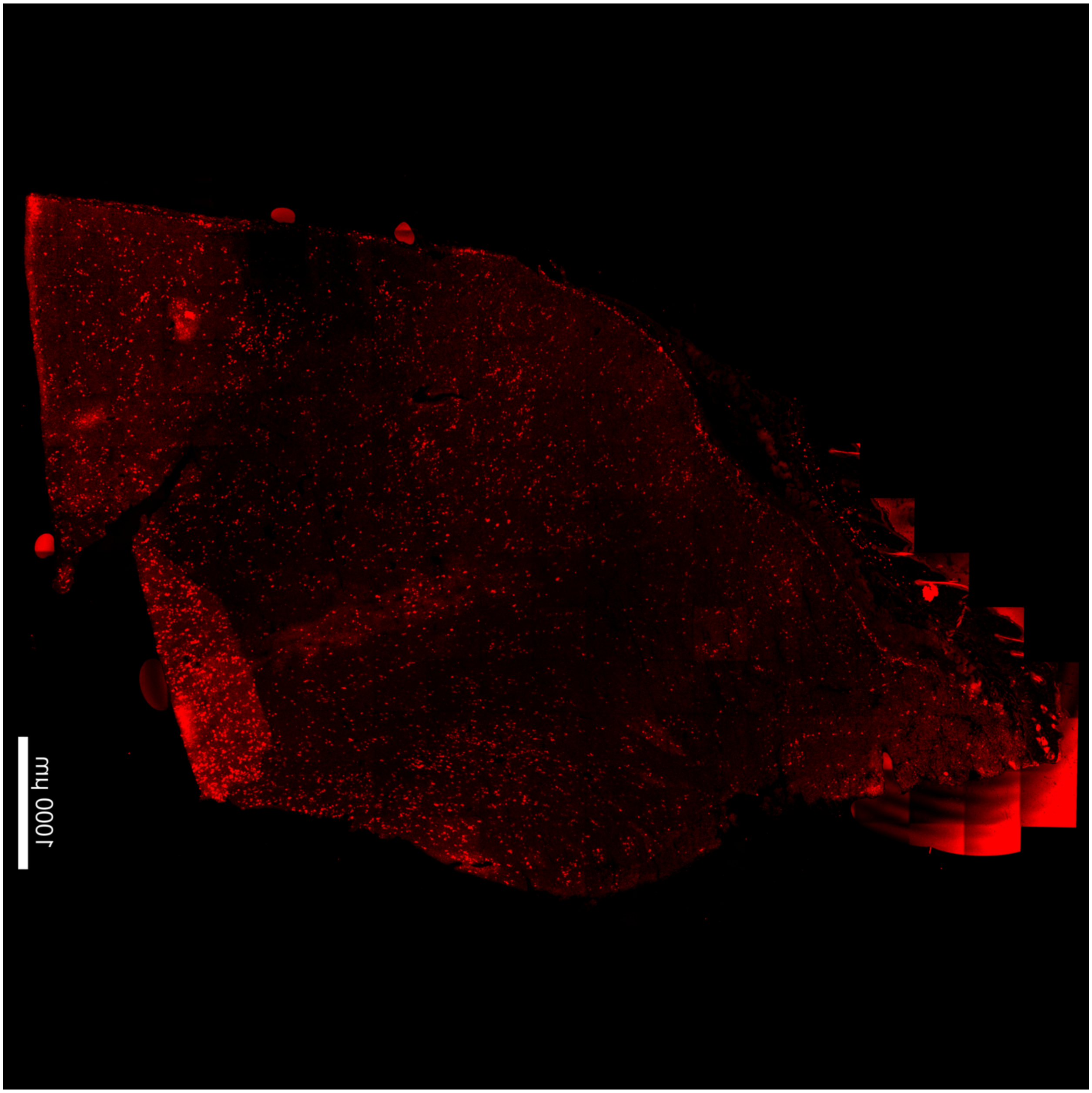


**Supplementary Figure 23.** Immunofluorescence imaging from the tumor using antibodies for NK1.1. Scale bar, 1000 µm.

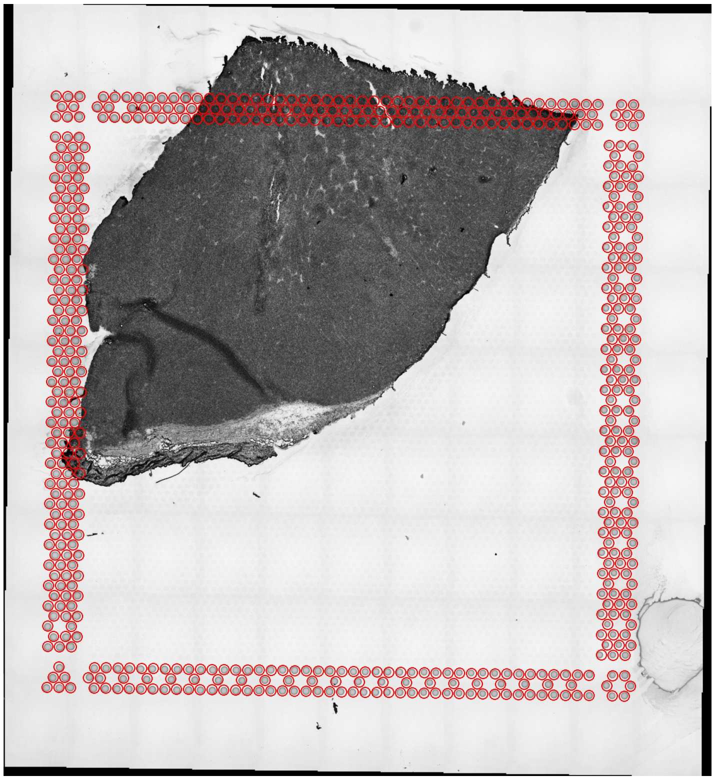

**Supplementary Figure 24**. Aligned fiducials images produced by the Space Ranger pipeline from 10x Genomics for the sample “Slide-1” presented in **Figure 4B**.

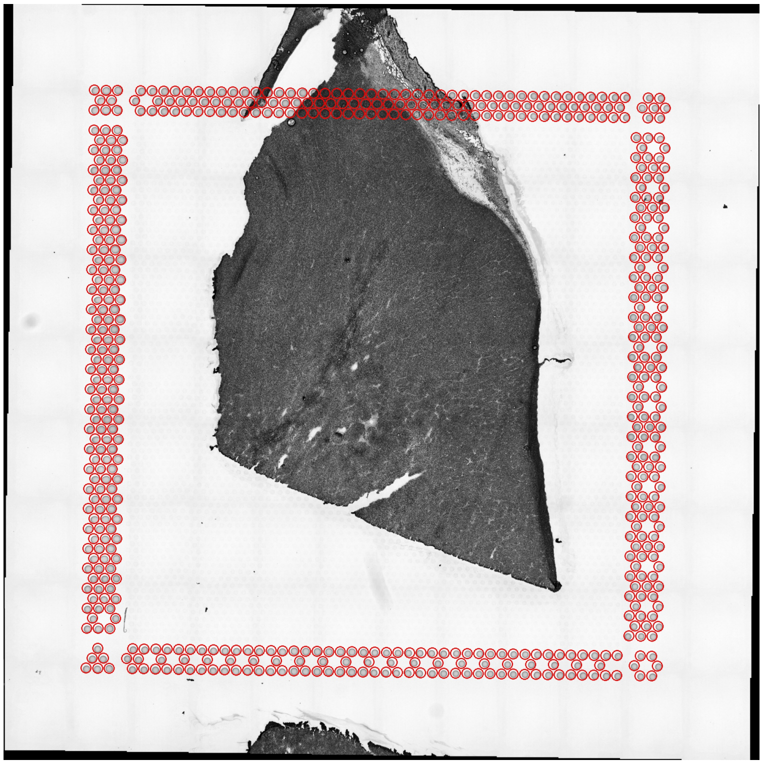

**Supplementary Figure 25.** Aligned fiducials images produced by the Space Ranger pipeline from 10x Genomics for the sample “Slide-2” presented in **Figure 4B**.

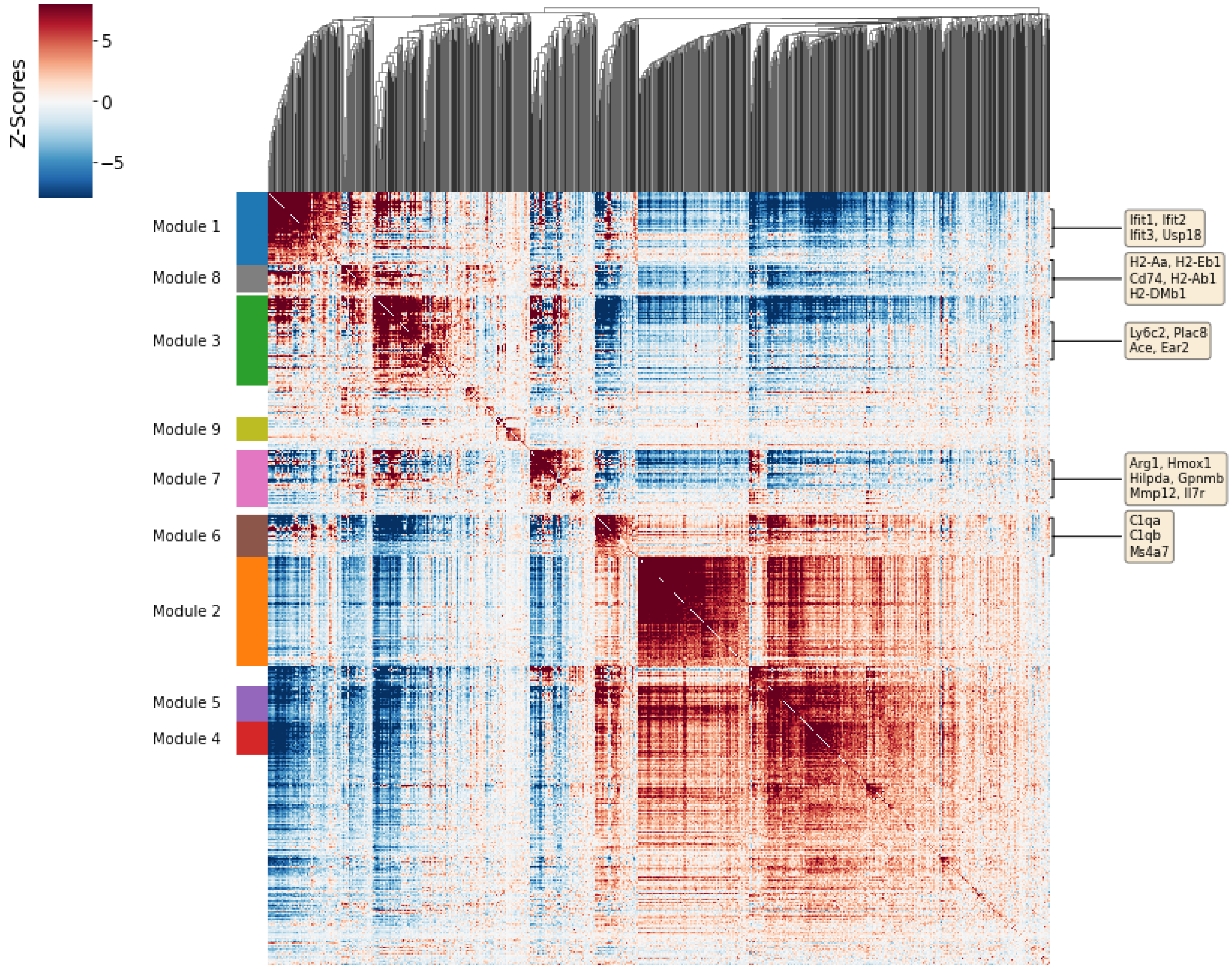


**Supplementary Figure 26.** Hotspot analysis for the Mon-Mac: gene-gene local autocorrelation matrix, and clustering into gene modules, based on the scRNA-seq data. We refer to the genes of module 1 as the Monocytes (IFN) module, to module 8 as Mon-DC, to module 3 as Monocytes (Ace), to module 7 as Mreg, and to module 6 as TAMs.

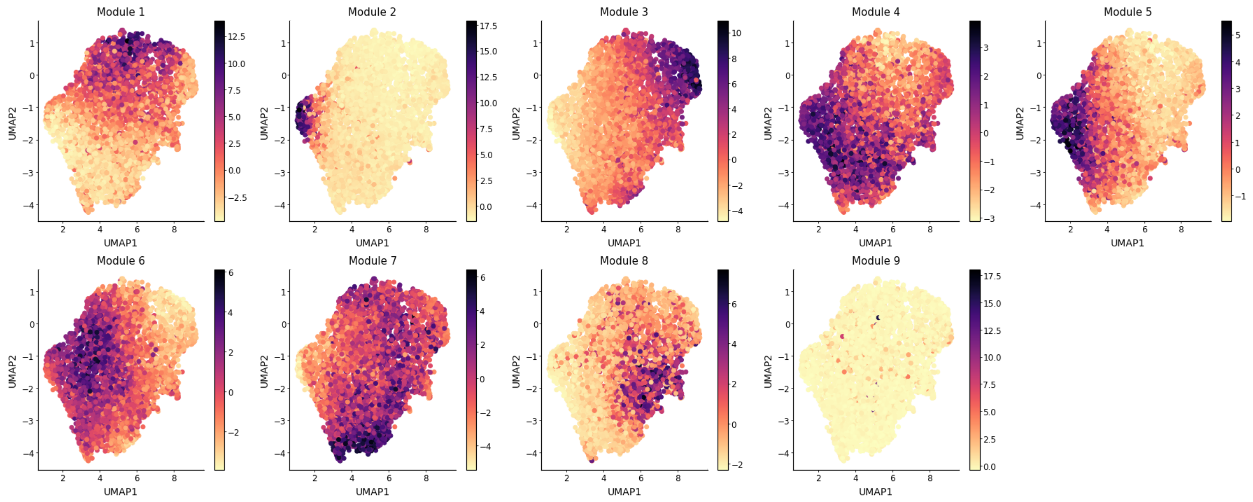


**Supplementary Figure 27.** Hotspot analysis for the Mon-Mac: projection of every gene module onto the embedding of the single-cell data (scLVM).

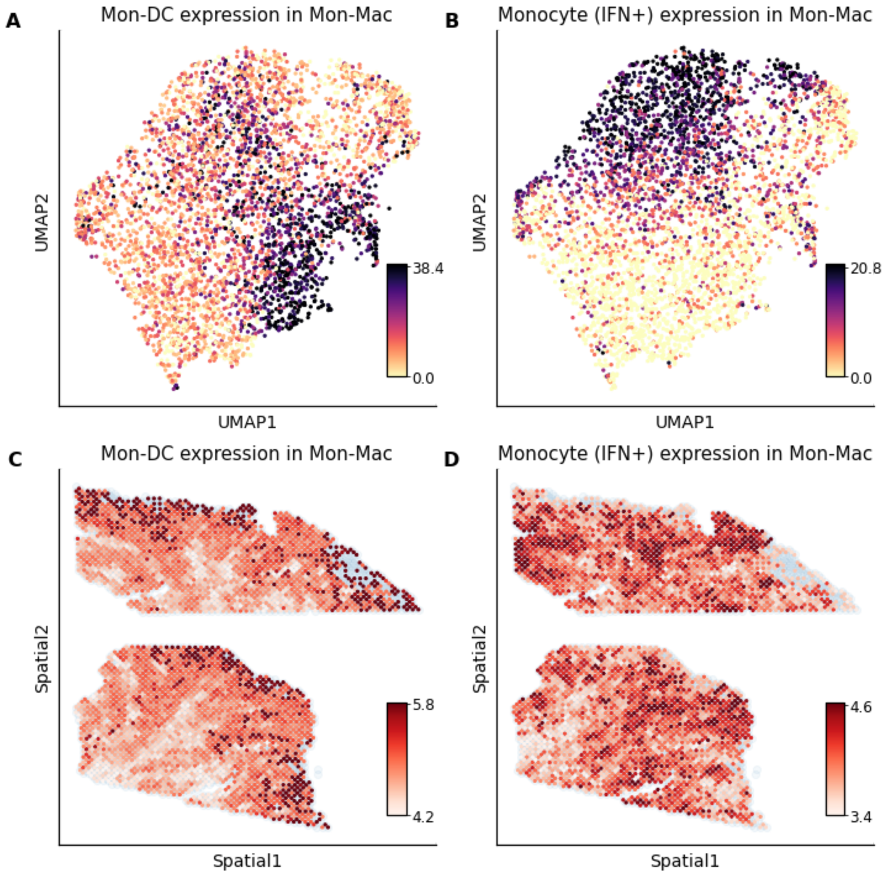


**Supplementary Figure 28. (A-B)** Visualization of selected gene expression modules inferred by Hotspot on the Mon-Mac cells from the scRNA-seq data. **(C-D)** Imputation of gene expression for those modules on the spatial dataset.

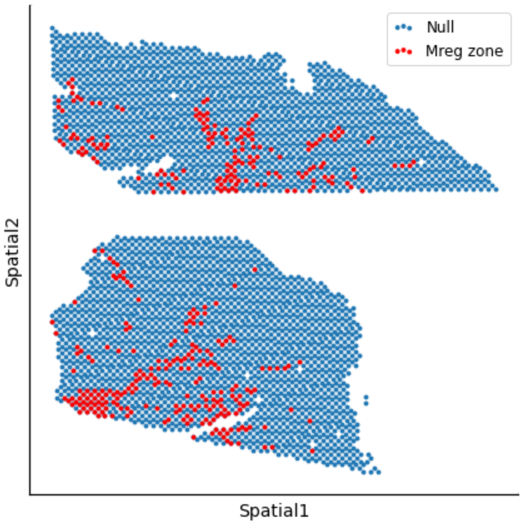


**Supplementary Figure 29.** Definition of the hypoxic zone (Mreg) for differential expression analysis.

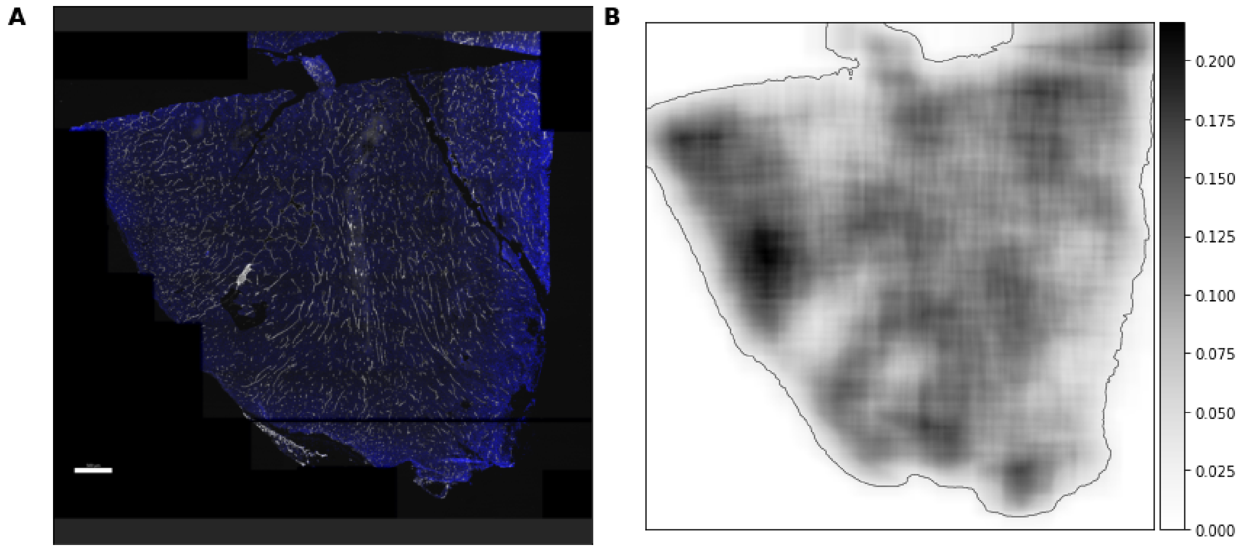


**Supplementary Figure 30.** Investigation of the hypoxic area. **(A)** Immunofluorescence imaging from the tumor using antibodies for CD31. Scale bar, 500 µm. **(B)** Percentage of blood vessels at each pixel of the image, evaluated by the proportion of pixels with high CD31 in the 150 x 150 pixels neighborhood (original image used for calculations is 1,500 x 1,500 pixels).

### Supplementary Tables Legend

**Supplementary Table 1 - Abundance of cell types in the lymph node scRNA-seq data**

**Supplementary Table 2 - Differential expression of Monocytes expression only, in spatial transcriptomics across conditions**

**Supplementary Table 3 - Differential expression of B cells expression only, in spatial transcriptomics across conditions**

**Supplementary Table 4 - Differential expression of B cells expression only, in MS lymph nodes (active area versus rest)**

**Supplementary Table 5 - Abundance of cell types in the scRNA-seq tumor data**

**Supplementary Table 6 - Differential expression of Mon-Mac expression only in spatial transcriptomics across Mreg abundant area against the rest of the tissue**

### Supplementary Note 1: DestVI report for the lymph node data

#### Step A: Automatic thresholding

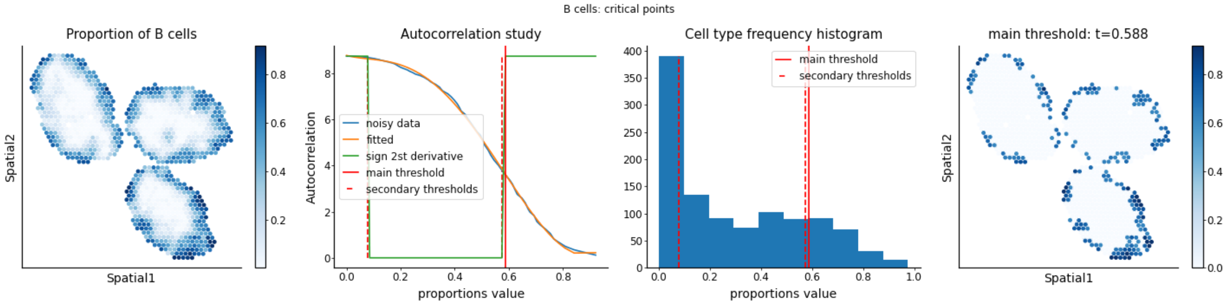


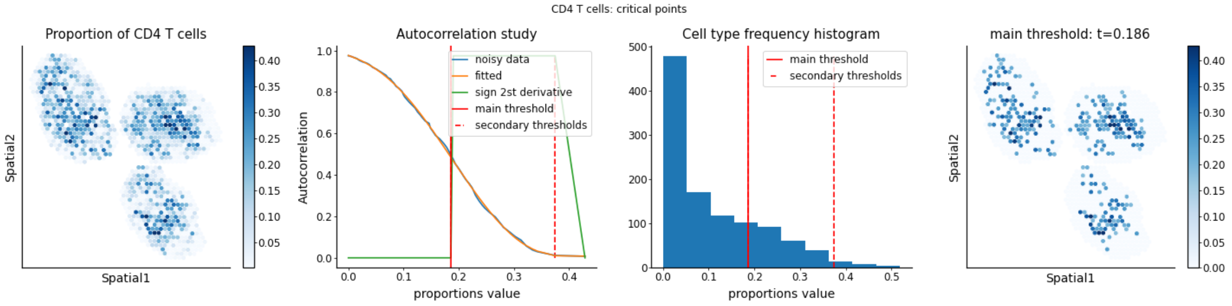


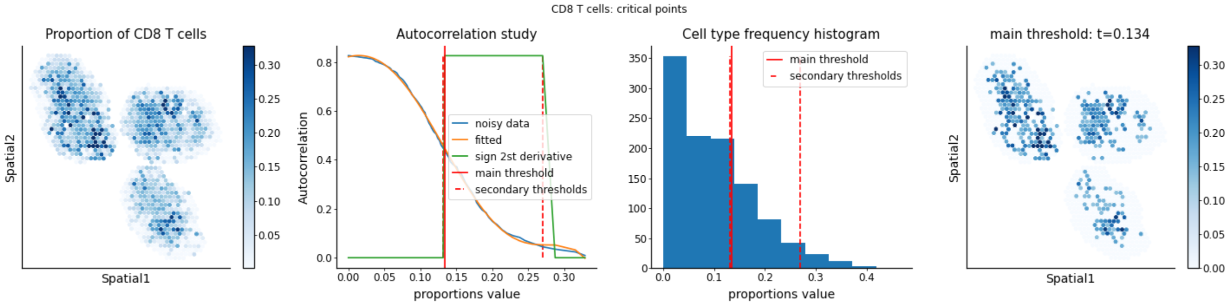

GD T cells: critical points

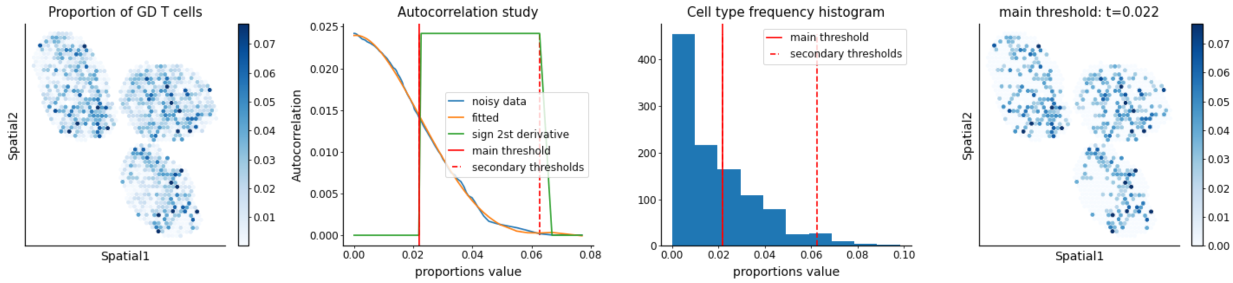


Macrophages: critical points

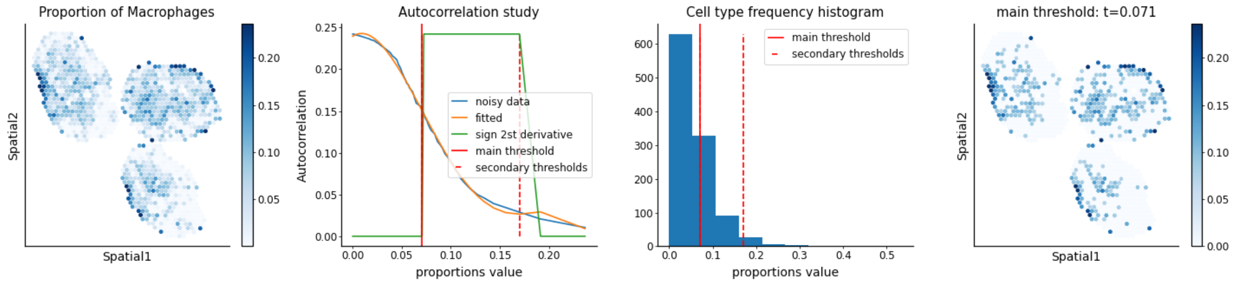


Migratory DCs: critical points

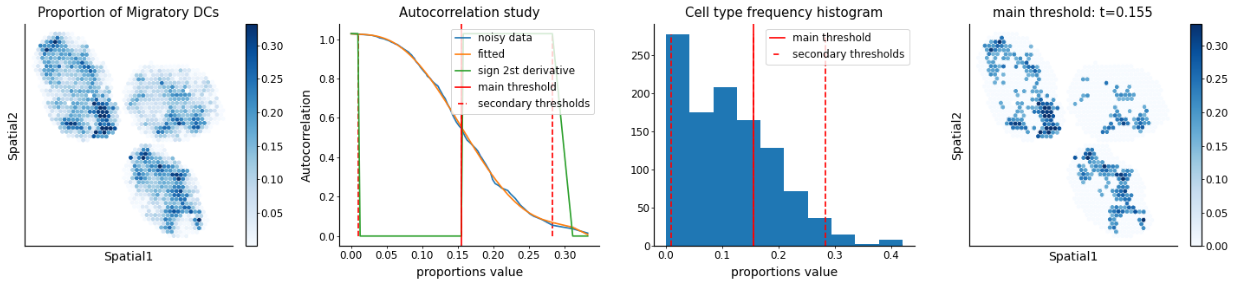


Monocytes: critical points

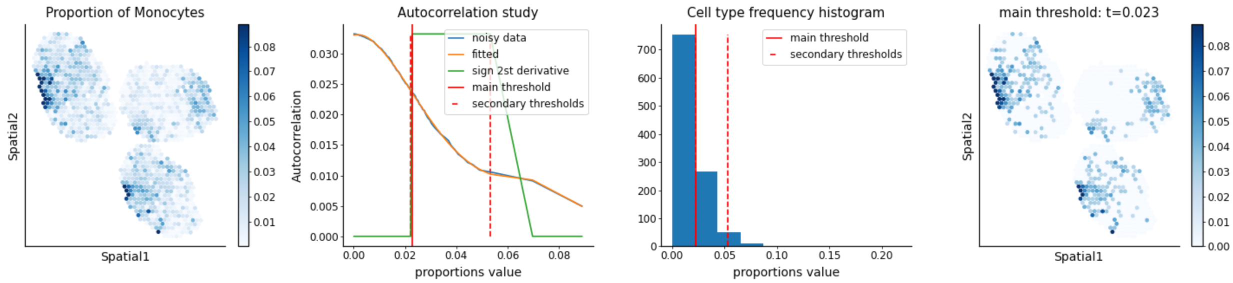

NK cells: critical points

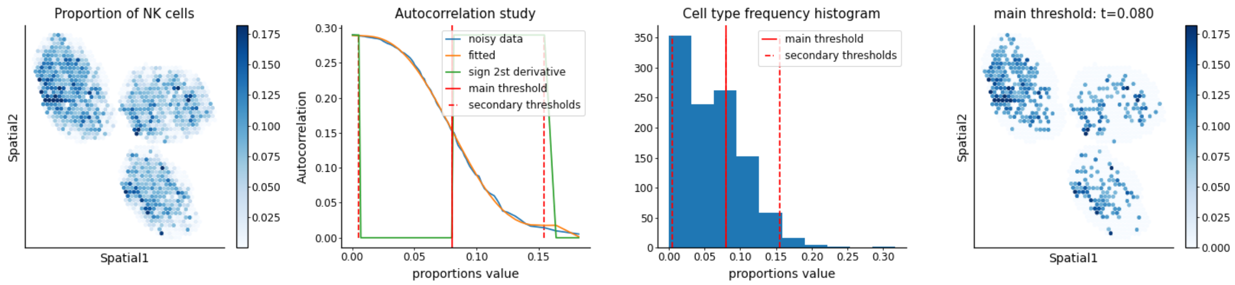


Tregs: critical points

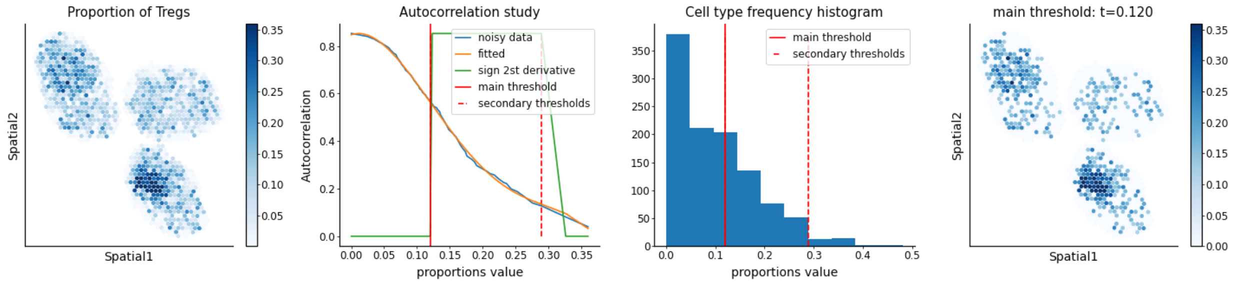


cDC1s: critical points

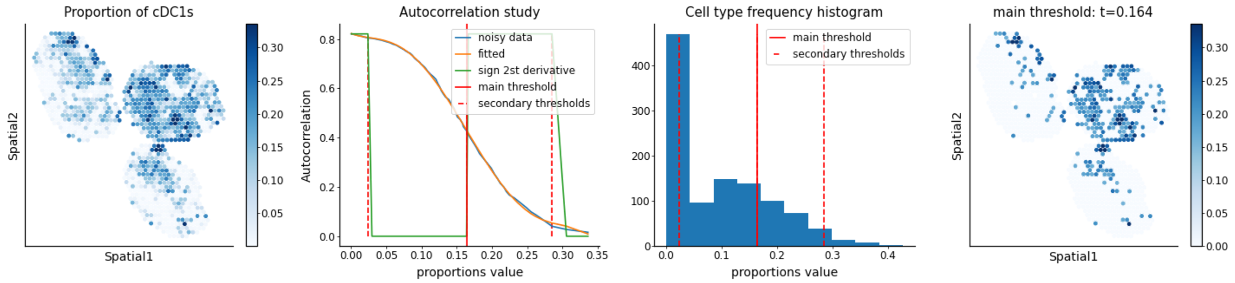

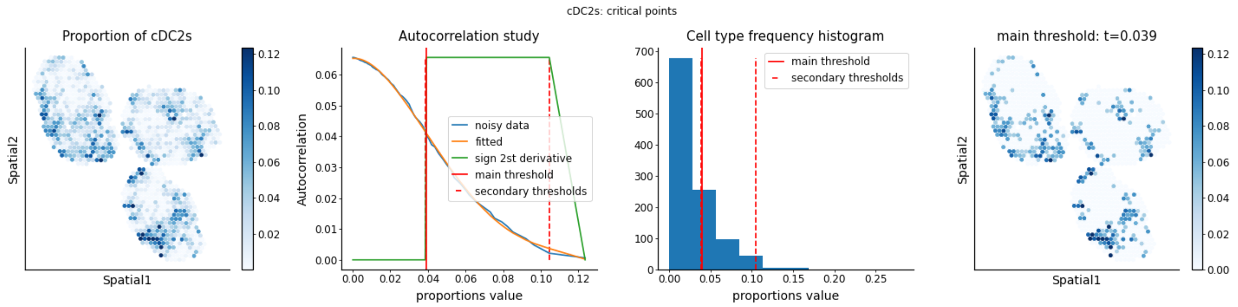


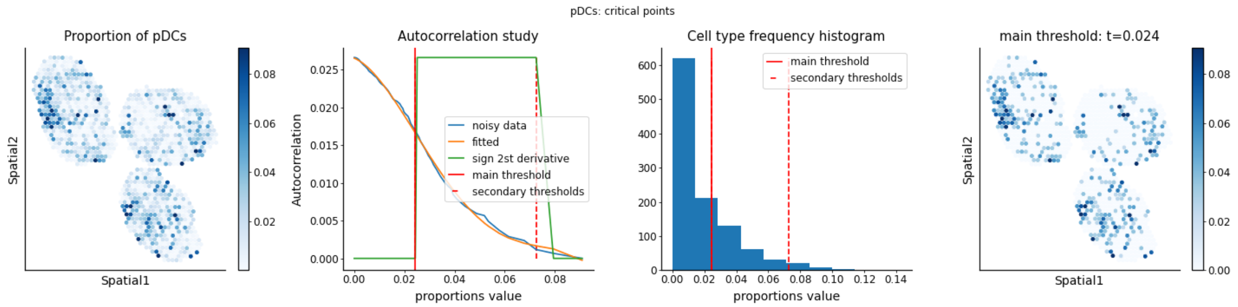


#### Step B: weighted PCA and GSEA

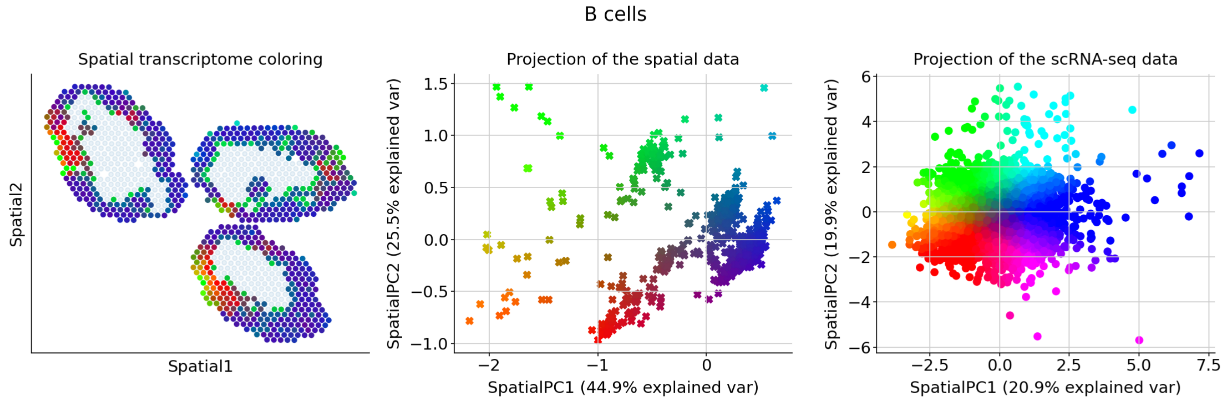


**Genes associated with SpatialPC1**

**Positively correlated**

Cldn1, Trem2, Mertk, Met, Tnnt2, Mfsd7a, Mab21l3, Ptprm, Mmp12, Syndig1l, Tarm1, Clec1b, Spint1, Cd300lb, Ear2, Pilrb1, Ppp1r3d, Itga8, Hpn, Hcar2, Ace, Ankrd35, Scn3a, Sirpb1c, Cd83, Nr4a1, Gm26532, Birc5, Bcl2a1b, Ncl, Ccna2, Npm1, Rplp0, Cks1b, Nme1, Impdh2, Mki67, Ccnb2, Ppp1r14b, Rps27l, Ube2c, Rel, Dennd4a, Tpx2, Cdca3, Dctpp1, Rps2, Ran, Hmmr, Atp5g1

FOXM1 transcription factor network*, Myc active pathway*, T cell receptor regulation of apoptosis*, Aurora B signaling*, Ran role in mitotic spindle regulation, Cyclin A/B1-associated events during G2/M transition, Cell cycle, Thymic stromal lymphopoietin (TSLP) pathway, Interleukin-11 pathway, Dilated cardiomyopathy

**Negatively correlated**

Ifit3, Slfn5, Irf7, Ifit3b, Rtp4, Stat1, Ighm, Usp18, Oasl2, Ifit1, Malat1, Cd79a, H2-Eb1, Ifit2, Zbp1, Serpina3g, Rsad2, Cd74, Hspa1b, Ifi47, Jun, H2-Aa, Igtp, H2-Ab1, Ifi27l2a, Oas3, Gbp4, Parp14, Trim30a, Isg15, Gbp7, Serpina3f, Pkib, Cmpk2, Irf1, Ly6d, Ifitm3, Rnf213, Isg20, Sdc3, Bst2, Fth1, Hist1h1c, Socs1, Ms4a4c, Igkc, Phf11b, Wfdc17, Csf1, Lgals3bp

Interferon alpha/beta signaling*, Interferon signaling*, Immune system signaling by interferons, interleukins, prolactin, and growth hormones*, Immune system*, Interferon-gamma signaling pathway*, Type II interferon signaling (interferon-gamma)*, Antiviral mechanism by interferon-stimulated genes*, Interferon alpha signaling regulation*, Prolactin activation of MAPK signaling*, Interferon-beta enhancer pathway*

##### Genes associated with SpatialPC2

**Positively correlated**

Mertk, Tnnt2, Mab21l3, Sirpb1c, Pilrb1, Ear2, Cldn1, Ptprm, Ppp1r3d, Clec1b, Cd300lb, Ankrd35, Spint1, Itga8, Scn3a, Mmp12, Hcar2, Hpn, Met, Tarm1, Ace, Syndig1l, Mfsd7a, Trem2, Rpsa, Rps2, Rplp0, Ppia, Ptma, Rpl41, Rplp1, Nme2, Rps12, Hsp90ab1, C1qbp, Rrm2, Mettl1, Npm1, Birc5, Timm8a1, Asf1b, Syce2, Ccna2, Dnph1, Cdk1, Stmn1, Ube2c, Mif, Impdh2, Cdc45

Cytoplasmic ribosomal proteins*, Influenza viral RNA transcription and replication*, Influenza infection*, Translation*, G1/S-specific transcription*, Cell cycle*, Disease*, E2F-mediated regulation of DNA replication*, Aurora B signaling*, FOXM1 transcription factor network*

**Negatively correlated**

Ifit3, Trim30a, Slfn5, Malat1, Irf7, Parp14, Stat1, Usp18, Rtp4, Fosb, Ly6a, Ifit1, Tcf4, Ifit3b, Rnf213, Rsad2, Ms4a1, Eif2ak2, Gbp7, Ifi27l2a, Pkib, Dennd4a, Zbp1, Plac8, Ifit2, Herc6, B2m, Cybb, Bcl11a, Isg15, Fos, Oasl2, Pim1, Klf4, Mpeg1, Cd86, Oas3, Junb, Ifi30, Bst2, Phf11b, Cd69, Ccnd2, Ly6d, Ly86, Vps37b, Zbtb20, Trafd1, Gm26532, Dusp1

Interferon signaling*, Interferon alpha/beta signaling*, Immune system signaling by interferons, interleukins, prolactin, and growth hormones*, Immune system*, Wnt interactions in lipid metabolism and immune response*, Interferon-gamma signaling pathway*, Type II interferon signaling (interferon-gamma)*, Antiviral mechanism by interferon-stimulated genes*, T cell receptor regulation of apoptosis*, Interleukin-3 regulation of hematopoietic cells*

CD4 T cells

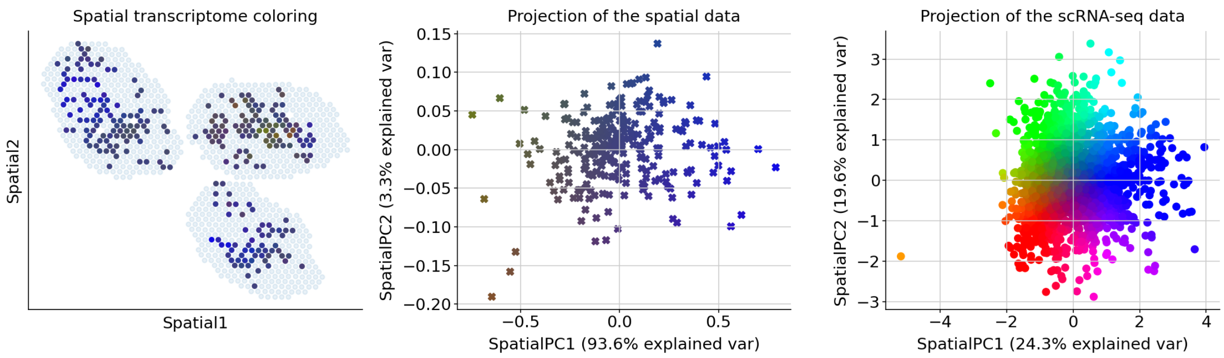


#### Genes associated with SpatialPC1

##### Positively correlated

Gpx3, Ncam1, Cblc, Tgm1, Il15, Cbr2, Cbr3, Tfec, Ccdc80, Mt2, Ccl17, Rasal2, Depdc1a, Mxd3, Ccl8, Ccl9, Gprc5c, Gpr141, Ccr1, Ccr10, Mycl, Tarm1, Mylk, Myof, Sirpb1b, Sirpb1c, Pilrb1, Rassf6, Cd14, Cd163l1, Igha, Thbd, P2ry6, Card10, Emilin2, Iqgap3, Cpne8, Erbb2, Gas7, Epha2, Tmod2, Ip6k3, Havcr1, Tmem51, Mmp12, Cox6a2, Il4, Slc7a11, Sh2d1b1, Cma1

Cytokine-cytokine receptor interaction*, Thymic stromal lymphopoietin (TSLP) pathway*, Intestinal immune network for IgA production, Binding of chemokines to chemokine receptors, Chemokine signaling pathway, Selective expression of chemokine receptors during T-cell polarization, Interleukin-7 interactions in immune response, Cytokines and inflammatory response, Arachidonic acid metabolism, Peptide G-protein coupled receptors

##### Negatively correlated

Rpsa, Rps2, Tmsb4x, Rplp0, Rplp1, Npm1, Rps12, Rpl41, Actg1, Ppia, Tagln2, Thy1, Vim, Actb, Cep55, Set, Crip1, Mif, Gm42418, S100a10, Eif4a1, Zyx, Ptma, Nme1, Gapdh, Chchd10, Cd3g, Lgals1, Dut, Spint2, Ckb, Slc25a4, Ptpn7, Adap1, Cd5, Atp1b1, Tubb5, Impdh2, Ybx1, Ddit4, Igfbp4, Tnfrsf9, Rbbp7, Ppp1r14b, Hist1h3c, Ube2c, Nme2, Mcm7, Ran, Hspe1

T cell receptor regulation of apoptosis*, Influenza infection*, Disease*, Translation*, Cytoplasmic ribosomal proteins*, Influenza viral RNA transcription and replication*, Protein metabolism*, Myc active pathway*, Activation of mRNA upon binding of the cap-binding complex and eIFs, and subsequent binding to 43S*, Nucleotide metabolism*

#### Genes associated with SpatialPC2

##### Positively correlated

Mt2, Klra6, Gpr35, Klra2, Plekhg1, Arhgap22, Gpr141, Stab2, Klra3, Klra7, Cd300c, Klra8, Styk1, Cysltr1, Sucnr1, Plcb1, Klra9, Ccr1, Klra17, Gpr4, Spry2, Gprc5c, Arpin, Guca1a, Ccl8, Gstm5, Sox13, Ccl9, Kif2c, Gramd2, Kif5a, Spint1, Gpx3, Plscr4, Klk1b27, Klra1, Spp1, Klrb1a, Klrb1b, Ccr10, Tfec, Ckap2l, Pf4, Tgm1, Thbd, Pde8b, Pde1c, Gcsam, Pcp4, Cd300a

Class A GPCRs (rhodopsin-like)*, Chemokine signaling pathway*, Binding of chemokines to chemokine receptors, Coagulation common pathway, cAMP cell motility pathway inferred from amoeba model, Eicosanoid metabolism, Kinesins, Signaling by GPCR, Fibrin clot formation (clotting cascade), GPCR ligand binding

**Negatively correlated**

Rps2, Rpsa, Nme2, Rplp0, Rplp1, Ppp1r14b, Ppia, Mettl1, C1qbp, Nme1, Ptma, Fbl, Srm, Ranbp1, Actb, Eif5a, Igfbp4, Npm1, Ppa1, Atp5g1, Gar1, Eif4a1, Prmt1, Hspa9, Mif, Nolc1, Tuba1b, Ran, Mrto4, Cct3, Shmt2, Phgdh, Uqcc2, Shmt1, Ssrp1, Prdx1, Rpl41, Nop58, Timm8a1, Psma7, Ccnd2, Actg1, Impdh2, Gnl3, Mthfd2, Pa2g4, Hspe1, Rexo2, Ly6a, Nhp2

Protein metabolism*, Disease*, T cell receptor regulation of apoptosis*, Myc active pathway*, Influenza infection*, Translation*, Cytoplasmic ribosomal proteins*, HIV infection*, Influenza viral RNA transcription and replication*, HIV factor interactions with host*

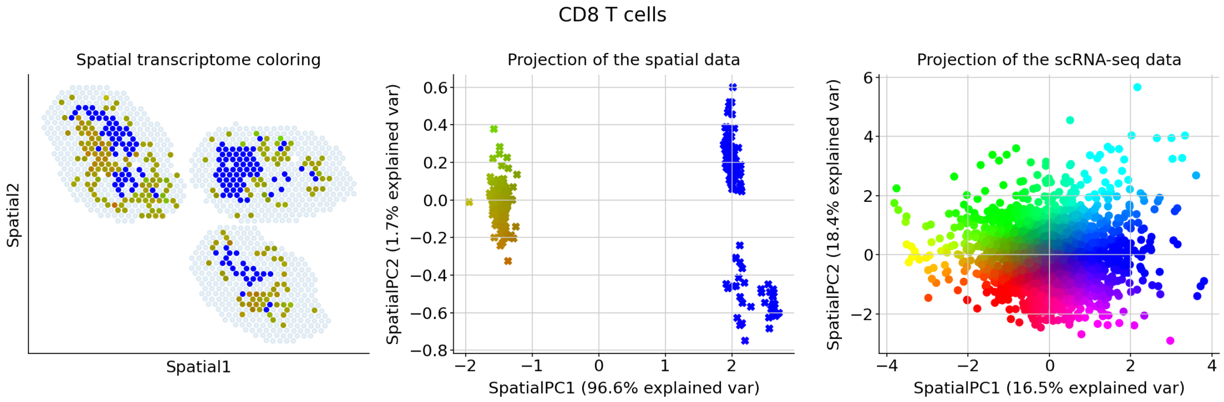


#### Genes associated with SpatialPC1

**Positively correlated**

Ccdc80, Gpr162, Cacna1b, Cd59a, Lipc, Ncam1, Fcgr4, Sirpb1b, Lmo7, Sirpb1c, Ffar2, Lrp1, Clec9a, Lrrc25, Htr7, Clec4g, Tmod2, Slc16a9, Clec4e, Fcgr1, Adam11, Il4, Adgrg6, Nuggc, Fads3, Nrxn1, Fam149a, Itga9, Gpr141, Cd300lb, Treml4, Crispld2, Itga8, Nlrp3, Cebpa, Scn3a, Sct, Gcsam, Sept3, Clec4d, Clec4a2, Tmem26, Cadm1, Ace, Tfec, Ip6k3, Mab21l3, Mmp12, Cadm3, Mfsd7a

Cell adhesion molecules (CAMs)*, Nectin/Necl trans heterodimerization*, Plexin D1 signaling, L1CAM interactions, Integrin family cell surface interactions, Dilated cardiomyopathy, Statin pathway, Adherens junction actin cytoskeletal organization, Signal transduction by L1, Urokinase-type plasminogen activator (uPA) and uPAR-mediated signaling

**Negatively correlated**

Tmsb4x, Gm42418, Ccl5, Fcer1g, Klrc2, Rplp1, Klra1, Rbpms, Klrk1, Klra7, Klre1, Klrc1, Crip1, Itga1, Ly6c2, Tcrg-C2, Tcrg-C1, Trdc, Selplg, Ncr1, Klra9, Tcrg-C4, Klra6, Anxa2, Klrb1c, Arl4d, Samd3, Pla2g7, Klrg1,

Rab7b, Lrrk1, Rpsa, Klrd1, Spry2, Ifitm10, Il1rn, Sirpa, Ifng, Fasl, Tyrobp, S100a6, Rab44, Klri2, Serpina3g, Cxcr3, Tnfaip2, Il7r, Gfra2, Cd5l, Havcr2

Natural killer cell-mediated cytotoxicity*, Leptin influence on immune response*, Immunoregulatory interactions between a lymphoid and a non-lymphoid cell*, T cell receptor regulation of apoptosis*, Ras-independent pathway in NK cell-mediated cytotoxicity*, Antigen processing and presentation*, Graft-versus-host disease*, Interleukin-2 signaling pathway*, Interleukin-1 regulation of extracellular matrix*, TNF-alpha effects on cytokine activity, cell motility, and apoptosis*

##### Genes associated with SpatialPC2

###### Positively correlated

Rab38, Tmem26, Fads3, Fam149a, Sirpb1c, Fcgr1, Fcgr4, Ffar2, Ffar4, Tlr13, Tlr11, Fndc7, Folr2, Fzd1, Ncam1, Thbs1, Gcsam, Gm12972, Gm13710, Etl4, Epha2, Nuggc, Tmod2, Cfh, Chil3, Nrxn1, Cldn1, Treml4, Clec1a, Clec1b, Clec4a2, Clec4d, Clec4e, Clec4g, Clec9a, Crispld2, Ehf, Nlrp3, Nid2, Sirpb1b, Tfec, Gm16685, Gm43916, Gpr141, Il10, Il13ra1, Il4, Ip6k3, Mab21l3, Itga8

Cell adhesion molecules (CAMs), Dendritic cells in regulating TH1 and TH2 development, Thymic stromal lymphopoietin (TSLP) pathway, Th1/Th2 differentiation pathway, Interleukin-4 signaling pathway, Asthma, Inflammatory response pathway, Cytokines and inflammatory response, Allograft rejection, Intestinal immune network for IgA production

###### Negatively correlated

Cd8b1, Stat1, Rplp1, Ifit3, Gbp2, Slfn1, Cd8a, Npc2, Epsti1, Igtp, Iigp1, Zbp1, Rtp4, Oasl2, Rplp0, Ifit1, Itm2b, B2m, Lef1, Isg15, AW112010, Saraf, Rpsa, Ly6a, Ms4a6b, Psmb9, Ifit3b, Gbp7, Cd3d, Irf7, Tspo, Ifi27l2a, Cd3g, Isg20, Ifi47, Rsad2, Herc6, Parp14, Trbc2, Rgs10, Gbp8, Art2b, Rnf213, Rpl41, Ddx60, Slfn5, Bst2, Oas3, Gbp4, Phf11b

Interferon signaling*, Interferon alpha/beta signaling*, Immune system signaling by interferons, interleukins, prolactin, and growth hormones*, Immune system*, Interferon-gamma signaling pathway*, Interleukin-12-mediated signaling events*, T helper cell surface molecules*, CTL mediated immune response against target cells*, Interleukin-17 signaling pathway*, T cell receptor signaling in naive CD8+ T cells*

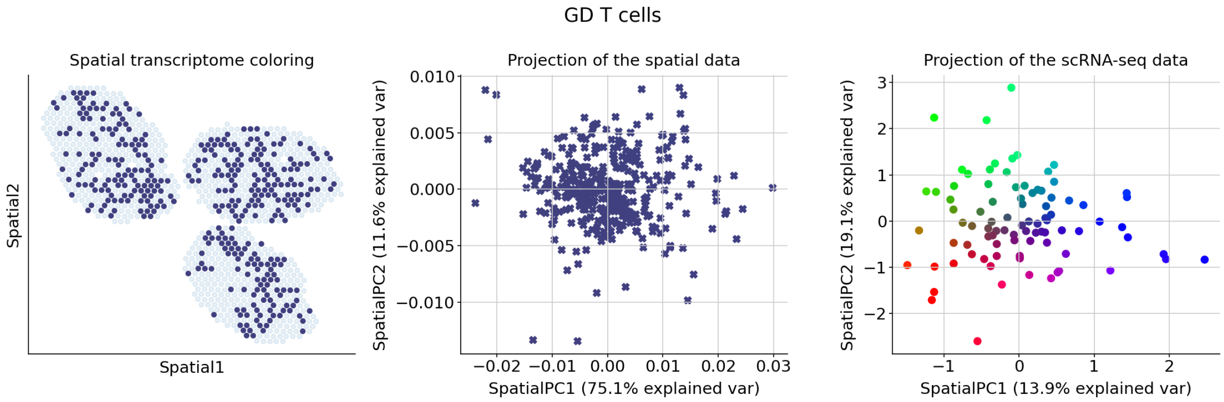

##### Genes associated with SpatialPC1

###### Positively correlated

Htra1, Clec5a, Scn3a, Cpne8, Sct, Hepacam2, Cp, Cox6a2, Col27a1, Notch3, Nostrin, Sema7a, Cmklr1, Sept3, Cma1, Clec4n, P2ry13, Clec4g, Clec4d, Serpinb8, Clec4b1, Serpinb9b, Clec4a3, Serpinf1, Serpinh1, Hfe, Clec4a2, Nlrp3, Clec4a1, Clec1b, Scin, Hcar2, Hap1, Crispld2, Gramd2, Gria3, Gtse1, Guca1a, Oaf, Dclk1, Nusap1, Gzma, Nuggc, Nuf2, Dapl1, Rsad2, Dab2, Rtn1, Nudt17, Cystm1

Extracellular matrix organization, Interleukin-9 regulation of target genes, Collagen biosynthesis and modifying enzymes, Endochondral ossification, Maturation of Notch precursor via proteolytic cleavage, NOSTRIN-mediated endothelial NOS trafficking, Pre-NOTCH Processing in the endoplasmic reticulum, Interleukin-1-beta (IL-1b) processing pathway, TSP1-induced apoptosis in microvascular endothelial cell, Gap junction degradation

###### Negatively correlated

S100a4, Fcer1g, Klrb1b, Plekho2, Serpina3g, Kit, Cd7, Dscam, Ms4a4c, Odc1, Rasl11a, Slc15a3, S100a6, Psd2, Dll1, Sema4c, Napsa, Fam110a, Anxa2, Lgals3bp, Il2rb, S100a1, Lmna, Nrgn, Bst2, Emb, Hmgcs1, Ttn, Lgals1, Tmem176a, Tyrobp, Ifngr1, Gdpd5, Psap, Fes, Tmem176b, Sdc4, Cdkn1a, Cldnd1, Cited4, Anxa1, Cdk6, Tnfsf12, Ncoa7, Cx3cr1, Spry2, Bcl2, Vegfa, Vim, Icosl

Cytokine-cytokine receptor interaction*, Leptin influence on immune response*, T cell receptor regulation of apoptosis*, Fibroblast growth factor 1*, EGFR1 pathway*, Interleukin-2 signaling pathway*, Interleukin-2/STAT5 pathway*, C-Myb transcription factor network*, Kit receptor transcriptional targets*, Prostaglandin biosynthesis and regulation*

##### Genes associated with SpatialPC2

###### Positively correlated

Depdc1a, Fcor, Fam20c, Fam83d, Fblim1, Fcgr2b, Fcgr4, Fcgrt, Fcrl5, Gpr4, Ffar4, Fgd2, Fignl1, Ngfr, Flt1, Rnd3, Mgl2, Fam149a, Fads3, Fabp4, F630028O10Rik, F13a1, F10, Exoc3l4, Rsad2, Exo1, Rtn1, Etl4, Espl1, Mgst1, Esco2, Eomes, Eno2, Rnase6, Flt3, Fn1, Gm12253, Melk, Rad51ap1, Rad54b, Rad54l, Rai14, Gpm6b, Gm7967, Gm6377, Gm5150, Gm43916, Mertk, Gm1673, Gm16685

Coagulation common pathway, Alpha-9 beta-1 integrin pathway, Homologous recombination, Fibrin clot formation (clotting cascade), RAGE pathway, Beta-1 integrin cell surface interactions, Axonal growth stimulation, Neurophilin interactions with VEGF and VEGF receptor, Extrinsic pathway, Complement and coagulation cascades

###### Negatively correlated

Ptma, Cd81, Lmo4, Ipo5, Swap70, Jak2, Plscr1, Hes1, Hsp90ab1, Cd83, Pxdc1, Rasl11a, Efnb2, Cct3, Ttn, Mapkapk2, Hspa9, Kdm2b, Chchd10, Vwa5a, Sptssa, Rnf19b, Pdia3, Cacybp, Ndufa12, Oasl2, Bzw2, Rrad, Psd2, Tgm2, Eif2ak2, Fam129c, Cks1b, Gata3, Creg1, Smarcc1, Klf4, Srm, Ppia, Sbf2, C2, Pkig, Il18bp, Dgat2, Hivep1, Il15ra, Tnfaip3, Ninj1, Ifi27l2a, Eea1

T cell receptor regulation of apoptosis, Immune system, Interleukin-2 signaling pathway, Notch signaling pathway, Toll receptor cascades, Innate immune system, TSH regulation of gene expression, Interleukin-27-mediated signaling events, Interleukin-1 regulation of extracellular matrix, Gastrin pathway

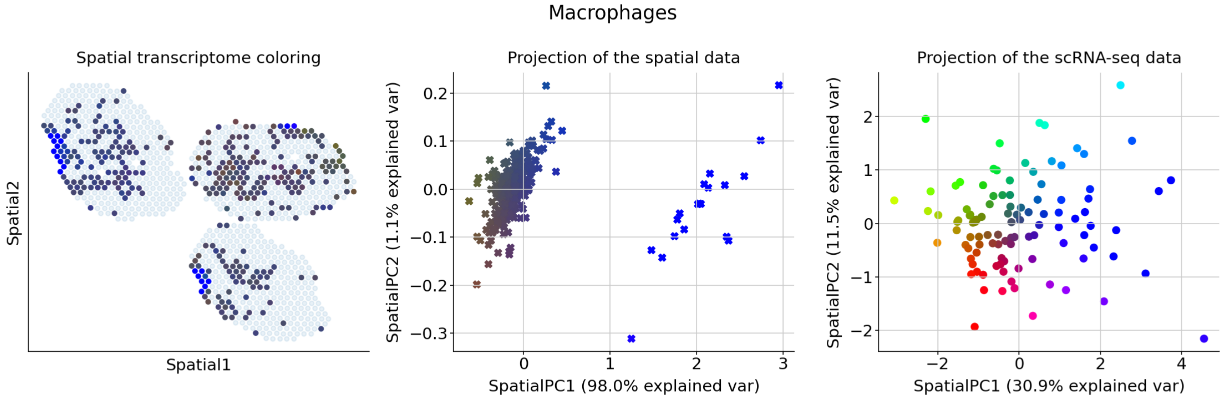


#### Genes associated with SpatialPC1

##### Positively correlated

Eme1, Gzmb, Clec1a, Penk, Slfn4, Gria3, Slpi, Pdzd4, Cldn1, Pdia5, Cited4, Smpdl3b, Smtn, Smyd1, Snai1, Pde1c, Cish, Pdcd1, Chl1, Gzma, Snx22, Chil3, Chdh, Slco3a1, Grasp, Gramd2, Phlda3, Slamf1, Slamf7, Cntnap1, Gm7967, Gm8251, Cma1, Pianp, Slc16a3, Pi16, Phf24, Pgam2, Gpr137c, Slc22a23, Clec9a, Clec4g, Gpr141, Clec4d, Clec4b1, Slc46a3, Gpr4, Pcp4, Gzmc, Sirpb1c

Interleukin-9 regulation of target genes*, Granzyme A-mediated apoptosis pathway, Prolactin regulation of apoptosis, Cell adhesion molecules (CAMs), Interleukin-12-mediated signaling events, Neurofascin interactions, Proepithelin conversion to epithelin and wound repair control, TSP1-induced apoptosis in microvascular endothelial cell, CHL1 interactions, Thymic stromal lymphopoietin (TSLP) pathway

##### Negatively correlated

Tmsb4x, Rplp0, Rps12, Rpsa, Rpl41, Rplp1, Fth1, Gpx1, Rps2, Gapdh, Ftl1, Sec61b, Cd200, Ppia, Lsp1, Tesc, Vim, Mif, Dap, S100a4, Lyz1, 1300017J02Rik, 2610528A11Rik, 2200002D01Rik, Capg, Peli2, Smim5, Dntt, Rnase6, Upb1, Cd7, Gsr, Havcr1, Bcl2a1b, Nme2, Uqcr11, Ank, Cox5a, Mt2, Iglc3, H2-Eb1, Hspa8, Nudt17, S100a6, Fscn1, Ppp1r14b, Apol10b, Cygb, Net1, Prg3

Translation*, Cytoplasmic ribosomal proteins*, Influenza viral RNA transcription and replication*, Influenza infection*, T cell receptor regulation of apoptosis*, Nucleotide metabolism*, Protein metabolism*, Free radical-induced apoptosis*, Activation of mRNA upon binding of the cap-binding complex and eIFs, and subsequent binding to 43S*, Cap-dependent translation initiation

#### Genes associated with SpatialPC2

##### Positively correlated

Pgam2, Rab44, Sox13, Eme1, Sox5, Klra6, Klra7, Klra8, Efnb3, Efna2, Ecm1, Ear2, Dll1, Dusp9, Dusp4, Klra9, Dtx1, Dscam, Dqx1, Spp1, Klrb1a, Spry2, Emilin2, Emp1, Eno2, Eomes, Slpi, Klk1, Fah, Klk1b27, F630028O10Rik, F2r, Klra1, Smpdl3b, Smtn, Smyd1, Snai1, Klra17, F10, Klra4, Exo1, Etl4, Erbb2, Snx22, Epha2, Dnm3, Dkkl1, Tcrg-C4, Cpne7, Ctla2a

MAP kinase pathway regulation through dual specificity phosphatases, EGFR1 pathway, Extrinsic prothrombin activation pathway, Acute myocardial infarction, Axon guidance, Blood clotting cascade, Neural crest differentiation, Glycolysis, Activated NOTCH1 signaling in the nucleus, Gluconeogenesis

**Negatively correlated**

Pla2g2d, Nhp2, Fabp5, Tagln2, Ndufc2, Tmem176a, Gapdh, Bcl2a1b, Tmem176b, Asb2, Traf1, Mt1, Cd63, Ncapd3, Ctsz, Clec5a, Sash1, Capg, Lyz2, Tspan13, Pdk3, Impdh2, Rps2, Cst7, Hspa8, Gpx1, Haus8, Mmp23, Eif5a, Gsap, 2900052N01Rik, Ran, Pgam1, Lyz1, Ftl1, Pla2g7, 2610528A11Rik, Phyhd1, Efhd2, Ctr9, C3, Atp1a3, Anxa3, Troap, M1ap, Aoah, Txndc17, Mcm3, Lgals3, Nudc

Lysosome vesicle biogenesis, Glycolysis, Gluconeogenesis, Ether lipid metabolism, Interleukin-5 regulation of apoptosis, Arachidonic acid metabolism, Clathrin derived vesicle budding, Glucose metabolism, Cycling of Ran in nucleocytoplasmic transport, Nucleotide metabolism

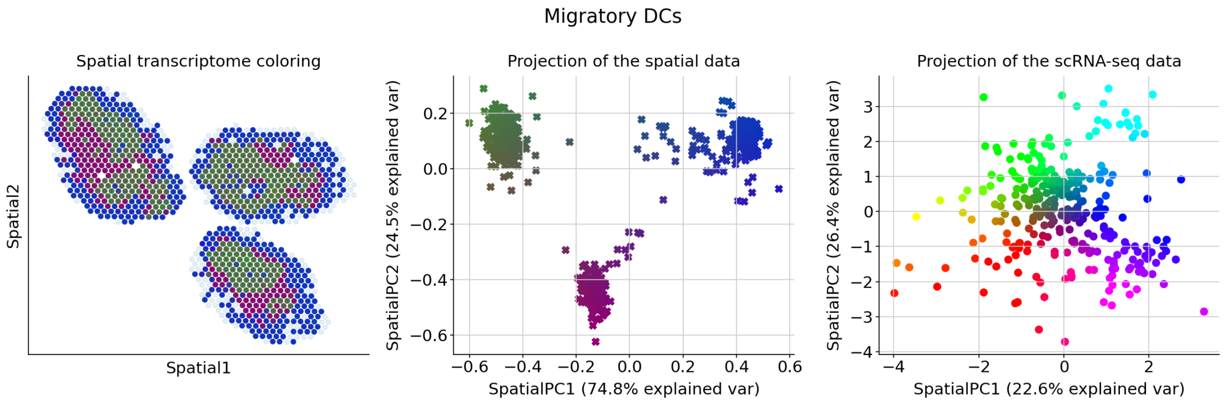


#### Genes associated with SpatialPC1

**Positively correlated**

Il4, Apoc1, Clec10a, Ifng, Pdcd1, Aatk, Ifitm6, Gcsam, Ifitm10, Penk, Pgam2, Pilrb1, Rgs13, Depdc1b, Depdc1a, Gm12972, Bpifb3, Gm13710, Pilra, Cma1, Tnfsf11, Parpbp, Anln, Galnt3, Nid2, Sapcd2, Cfh, Itga8, Itga2, Trem2, Atp1a3, Trav3-3, Bhlhe41, Ptgs1, Nuggc, Il10, Samd3, Chdh, Chil3, Sox5, Chl1, Cmklr1, Pilrb2, Folr2, Rab3il1, Gpsm2, Stc2, Prg4, Havcr1, Prkar1b

Dendritic cells in regulating TH1 and TH2 development*, Th1/Th2 differentiation pathway*, AP-1 transcription factor network*, Cytokines and inflammatory response*, Allograft rejection*, CHL1 interactions*, T cell receptor signaling pathway, Folate metabolism, Leishmaniasis, Tob role in T-cell activation

**Negatively correlated**

S100a4, Fabp5, Tmem176a, Ptms, Sec61b, Syngr2, Sub1, Tmem176b, Ybx1, Ctsz, Actg1, Cd74, Tubb5, Tmsb4x, Ifi30, Nme2, H2-Ab1, Ftl1, Calm3, Lamp1, Bri3, Gapdh, S100a6, Atox1, H2-Eb1, Marcksl1, Actb, Hnrnpa1, Avpi1, Ctss, Rpl41, Gpx1, Snrpd1, Rplp0, Calr, Nap1l1, Hspa8, Bst2, Rps12, B2m, Rpsa, Ywhae, Ppia, Srgn, Gpr183, Lsp1, Clic4, S100a11, Fscn1, Rps27l

Antigen processing and presentation*, Translation*, Phagosome*, T cell receptor regulation of apoptosis*, Influenza infection*, Antigen processing: cross presentation*, Protein metabolism*, Cytoplasmic ribosomal proteins*, Endosomal/vacuolar pathway*, Interleukin-2 signaling pathway*

##### Genes associated with SpatialPC2

###### Positively correlated

Cox4i2, Hist1h3c, Lingo4, Lipc, Coro2b, Tgm1, Cmklr1, Emp1, Cma1, Bcat1, Lrr1, Tcrg-C4, Rgs13, Bhlhe41, Bpifb3, Tbxas1, Pgam2, Ltb4r1, Tacstd2, Syndig1l, Penk, Hcar2, Havcr1, Styk1, Clec10a, Emilin2, Rassf6, Hp, Hlf, Klk1b27, Rab3il1, Klra1, E2f7, Rab44, Plcb1, Klra2, Klra4, Tnfsf11, Cryba4, Klra7, Klra8, Klra9, Klrb1a, Klrb1c, Klrc1, Klrg1, Pilrb2, Pilrb1, Atp1a3, Kntc1

Inhibition of platelet activation by aspirin, Eicosanoid metabolism, G-protein signaling through tubby proteins, Cytochrome P450 metabolism of eicosanoids, Gastric acid secretion, Cardiac muscle contraction, Activation of PKC through G-protein coupled receptors, Insulin secretion regulation by fatty acids bound to GPR40 (FFAR1), Salivary secretion, Phospholipase C signaling pathway

###### Negatively correlated

Malat1, Neat1, Birc2, Slco5a1, Ankrd33b, Adam23, Aebp2, Tmcc3, Ccdc88a, Klf6, Dennd4a, Lrrk1, Swap70, Sbno2, Mical3, Fnbp1l, Tbc1d4, Chka, Adam11, Kdm2b, Atp11a, Nrgn, Ciita, Arhgap31, Gm15987, Zeb2, Etv3, Traf1, Cacnb3, Marveld1, Atp2b1, Col27a1, Rel, Lyst, Sbf2, Frmd4a, Zfp36, Nav1, Marcks, Plce1, Birc3, Icosl, Sned1, Nrxn1, Tmpo, Trim35, Tns1, Plxnc1, Il1r1, Zbtb20

CD40/CD40L signaling*, TNF-alpha signaling pathway*, Keratinocyte differentiation, HIV-1 Nef as negative effector of Fas and TNF, Apoptosis modulation and signaling, Small cell lung cancer, Mitochondrial role in apoptotic signaling, Tumor necrosis factor (TNF) pathway, Apoptosis, Induction of apoptosis through DR3 and DR4/5 death receptors

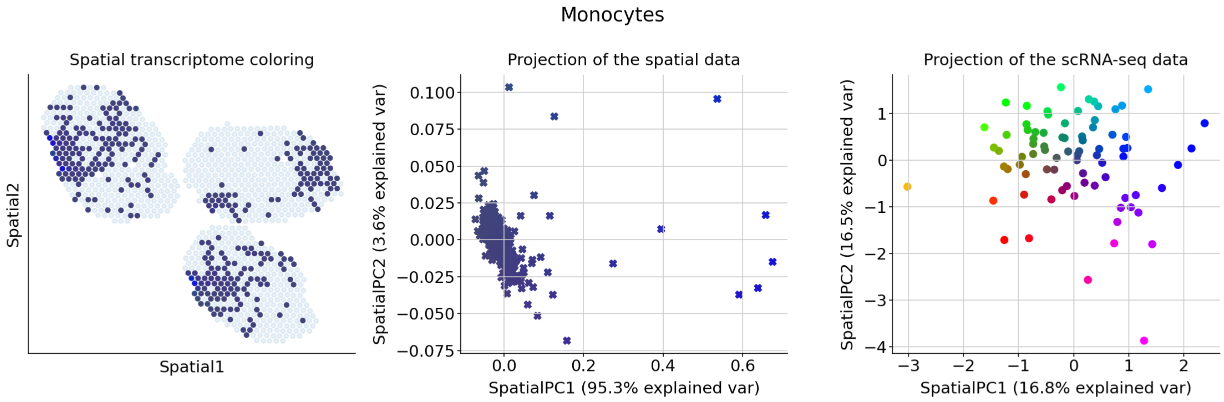

##### Genes associated with SpatialPC1

###### Positively correlated

Ighg3, Hmmr, Eme1, Emp1, Il1rl1, Il2ra, Eno2, Cd163l1, Cd160, Eomes, Il4, Stab2, Insm1, Stc2, Stil, Epha2, Ccr4, Strip2, Ptprm, Cd209d, Mdk, Nebl, Cd247, Ect2, Socs2, Rad51, Nefh, Sox13, Efnb3, Sox5, Spag5, Neil3, Cd5l, Spc25, Cd59a, Nek2, Cd300c, Melk, Il18r1, Spp1, Iqgap3, Styk1, Ptprk, Ptprf, Tcrg-C1, Mxd3, Tcrg-C4, Cblc, Hist1h2ag, Capsl

Interleukin-12-mediated signaling events*, Jak-STAT signaling pathway*, Tob role in T-cell activation*, Th1/Th2 differentiation pathway*, Selective expression of chemokine receptors during T-cell polarization*, Interleukin-12/STAT4 pathway*, Regulation of NFAT transcription factors*, Type I interferon (interferon-alpha/beta) pathway*, Hyaluronan metabolism, CD8/T cell receptor downstream pathway

###### Negatively correlated

Ppia, Ptma, Atox1, Smc2, Rps2, S100a4, Ifngr1, Cdk14, Spdl1, Mcm10, Tuba1c, Cbx5, Chaf1a, Cdk4, Eif4ebp1, Eif4a1, Rplp1, Tuba1a, Pira2, Exo1, Dbn1, Rad51ap1, Kntc1, Ube2t, Dscc1, Cenpp, Odc1, St8sia4, Cd1d1, Hnrnpab, Ptpn22, Ndufc2, Shmt1, Rexo2, Tacc3, Rrm1, Dhfr, Cbfa2t3, Hmgb1, Ncapg, Uqcc2, Il1b, Nr4a3, Pip4k2a, Cd6, Fut7, Rplp0, Sptssa, Eif1ax, Plxnc1

Translation*, Myc active pathway*, Activation of mRNA upon binding of the cap-binding complex and eIFs, and subsequent binding to 43S*, Protein metabolism*, Cap-dependent translation initiation*, Cell cycle*, Aurora B signaling*, Cytoplasmic ribosomal proteins*, Folate and pterine metabolism*, Translation factors*

##### Genes associated with SpatialPC2

###### Positively correlated

Hba-a1, Lefty1, Ryr3, Lipc, Samd3, Lingo4, Sapcd2, Etl4, Ccne1, Ccna2, Scin, Scn3a, Ccl22, Sema4c, Klri2, Ccl17, F2r, Sept3, Ccdc80, Fads3, Lcn4, Fah, Laptm4b, Serpinb9b, Serpine2, Cblc, Ccne2, Ccr10, Esco2, Ccr4, Rassf6, Ly6k, Cd5l, Eomes, Cd59a, Ly6c1, Epha2, Rgs13, Rgs16, Rgs9, Cd300c, Lta, Cd247, Cd209d, Lrrc32, Rorc, Lrr1, Cd163l1, Cd160, Erbb2

Binding of chemokines to chemokine receptors*, G0 and early G1 pathway*, Peptide G-protein coupled receptors*, Cytokine-cytokine receptor interaction, G alpha q pathway, Cyclin A-Cdk2-associated events at S phase entry, E2F transcription factor network, Chemokine signaling pathway, Prostate cancer, Signaling events mediated by PRL

###### Negatively correlated

Prkar1b, Gfpt1, Gfra2, Zc3h12c, Dusp5, Fosb, Chst2, Ly75, Il10, Anln, Hsd11b1, Tnfrsf8, Egr1, Ddx60, Hsph1, Fam20c, Stmn1, Kcnk6, Tmem26, Gpr157, Pfkfb3, Slc6a6, Sirpa, Adgrg6, Gpm6b, Sdc3, Cd274, Tox, Tnfaip3, Dusp1, Nuak1, Ccnd2, Ehf, B2m, Prg3, Pmaip1, Rai14, Marcksl1, Art2b, Tgm1, Susd2, Tspan3, Cd81, Trpm2, Rgs1, Gm42418, Marcks, Mmp23, Inpp4b, Pvr

BDNF signaling pathway*, T cell receptor regulation of apoptosis*, AP-1 transcription factor network*, Malaria, CD40L signaling pathway, TNFR2 signaling pathway, Interleukin-2 signaling pathway, TSH regulation of gene expression, Ovarian infertility genes, Stathmin and breast cancer resistance to antimicrotubule agents

NK cells

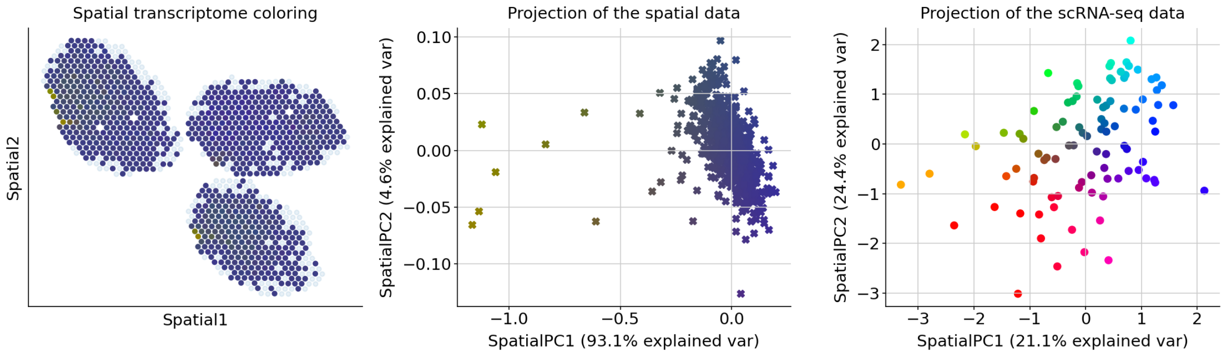


##### Genes associated with SpatialPC1

###### Positively correlated

1300017J02Rik, Gprc5c, Gm5150, Gpm6b, Gpr157, Rem1, Gpr162, Gpr35, Gpr4, Gpx3, Prg3, Rbm47, Gramd2, Gria3, Rassf6, Rasl11a, Rasal2, Gsta4, Gm43916, Rgs13, Gm1673, Rgs9, Rpl39l, Gatm, Gbp2b, Rorc, Gcsam, Gda, Gdpd1, Gfra2, Gm10851, Gm12253, Rnf144b, Gm12972, Gm13710, Rnd3, Gm16685, Gstm5, Gtse1, Guca1a, Hlx, Ptafr, Psrc1, Hmgn3, Hp, Psd2, Hpgd, Hpn, Hpse, Hs3st1

Glutathione metabolism, Visual signal transduction: cones, Visual signal transduction: rods, Glutathione conjugation, Phototransduction, Heparan sulfate/heparin glycosaminoglycan (HS-GAG) metabolism, Metapathway biotransformation, G alpha q pathway, Phase II of biological oxidations: conjugation, Creatine metabolism

###### Negatively correlated

Rsad2, Tcrg-C4, Usp18, Igtp, Il2ra, Isg15, Ifit1, Il7r, Cxcl10, Ifi44, Rnf19b, Ifi204, Gpr183, Cxcr6, Ckb, Klk8, Irf7, Fas, Ly6a, Parp12, Cmpk2, Il6st, Ifit3, Sema4c, Zbp1, Herc6, Klrc1, Myo6, Iigp1, Gbp2, Oas3, Maf, Hopx, Cd82, Xcl1, Dscam, Cd3e, Gbp5, Ifih1, Socs3, Ikzf2, Cish, Tmem176b, Isg20, Socs1, Foxp3, Lilrb4a, Thy1, Batf, Furin

Immune system signaling by interferons, interleukins, prolactin, and growth hormones*, Interferon alpha/beta signaling*, Immune system*, Interferon signaling*, Interferon-gamma signaling pathway*, Interleukin-2 signaling pathway*, Leptin influence on immune response*, Thymic stromal lymphopoietin (TSLP) pathway*, Cytokine-cytokine receptor interaction*, Interleukin-4 regulation of apoptosis*

##### Genes associated with SpatialPC2

###### Positively correlated

1300017J02Rik, Htr7, Hpgd, Hpn, Sash1, Hpse, Sapcd2, Hs3st1, Htra1, Igfbp7, I830077J02Rik, Ryr3, Ifitm6, Ifnlr1, Igf1, Igfbp4, Hp, Hmgn3, Hlx, Hist1h3c, Scin, Scn3a, Hfe, Sct, Hepacam2, Hcar2, Hbegf, Hba-a1, Havcr2, Havcr1, Hap1, Hacd4, Sept3, Rtn1, Igha, Guca1a, Rnd3, Itga8, Itga9, Itgb8, Rnf144b, Izumo1r, Jag1, Jchain, Ighg3, Kcnj10, Kcnj2, Kcnk6, Rgs9, Zmynd15

Integrin family cell surface interactions*, Dilated cardiomyopathy*, Ghrelin-mediated regulation of food intake and energy homeostasis, Insulin-like growth factor (IGF) activity regulation by insulin-like growth factor binding proteins (IGFBPs), Arrhythmogenic right ventricular cardiomyopathy (ARVC), ECM-receptor interaction, Integrin cell surface interactions, Visual signal transduction: cones, Visual signal transduction: rods, Plexin D1 signaling

**Negatively correlated**

Hsp90ab1, Odc1, Cct3, Nhp2, Eif5a, Hnrnpab, Rsl1d1, Kdm6b, Ftsj3, Fosl2, Hnrnpa1, Nr4a1, Smco4, Tnfrsf21, Nop58, Bzw2, C1qbp, Gm26532, Tagap, Tuba1b, Srm, Dtl, Mrto4, Nop16, Eif4a1, Nfkbiz, Ncl, Gadd45b, Ndufab1, Shmt1, Nfkbid, Ipo5, Hells, Vegfa, Prmt1, Uqcc2, Timeless, Mettl1, Apex1, Ran, Hdgf, Dkc1, Ifng, Cd69, Tex2, Mybbp1a, Hsph1, Smc2, Psma7, Npm1

Myc active pathway*, T cell receptor regulation of apoptosis*, Telomere extension by telomerase*, Aurora B signaling*, Polyamine metabolism, Telomerase regulation, Interleukin-2 signaling pathway, Hypertrophy pathway, Urea cycle and metabolism of amino groups, Amino acid metabolism

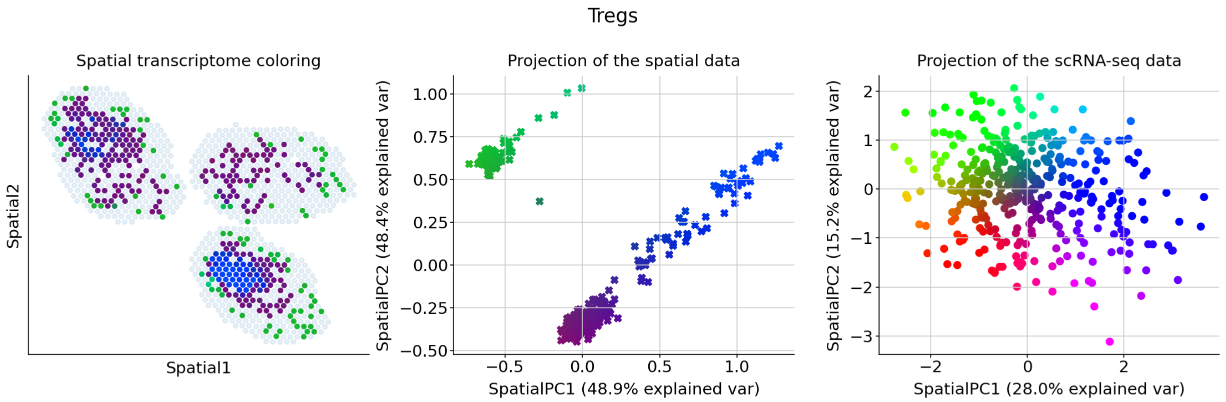


#### Genes associated with SpatialPC1

**Positively correlated**

Emilin2, Myof, Npl, Notch3, Arpin, Nlrp3, Nid1, Nek2, Cdh2, Ncr1, Mylk, Nrxn1, Arhgef40, Arhgef10l, Arhgap28, Mycl, Stap2, Msr1, Ms4a8a, Ms4a7, Cd5l, Nuggc, Thbd, Cd300lb, Pdzd4, Snx22, Pdia5, Pde8b, Cd300c, Pcp4, Pbx1, Sox13, Papss2, Nupr1, Cd300ld, Cd302, P2ry6, P2ry13, Cd33, Cd36, Nxpe4, Spic, Mreg, Strip2, Mrc1, Lrr1, Tbc1d9, Mab21l3, Ltc4s, Cldn1

Nucleotide-like (purinergic) G-protein coupled receptors, Cell adhesion molecules (CAMs), Phagosome, Maturation of Notch precursor via proteolytic cleavage, Pre-NOTCH Processing in the endoplasmic reticulum, Interleukin-1-beta (IL-1b) processing pathway, Antigen processing: cross presentation, Cell junction organization, Cross-presentation of particulate exogenous antigens (phagosomes), TSP1-induced apoptosis in microvascular endothelial cell

**Negatively correlated**

Tnfrsf4, Il2ra, Ikzf2, Ikzf4, Foxp3, Izumo1r, Tnfrsf18, Igf1r, Tiam1, Tnfrsf9, Lrrc32, Dusp1, Swap70, Cd81, Ptma, Hivep3, Gpr83, Ecm1, Malat1, Cd83, Itgb8, Arhgap20, Ift80, Tbc1d4, Kdm6b, Snx18, Junb, Mical3, Cd4, Cish, Chchd10, Dst, Gnl3, Rel, Smc4, Mettl1, Rgs10, Myo1e, Rgs2, Plagl1, Lta, Gm42418, Pim1, Eea1, Mrps6, Gsta4, Gsto1, Spred1, Nfkbia, Myc

T cell receptor regulation of apoptosis*, Thymic stromal lymphopoietin (TSLP) pathway*, Interleukin-9 regulation of target genes*, CD8/T cell receptor downstream pathway*, Interleukin-2/STAT5 pathway*, TNFR2 signaling pathway*, Interleukin-2 signaling pathway*, Jak-STAT signaling pathway*, Antigen-activated B-cell receptor generation of second messengers*, Wnt interactions in lipid metabolism and immune response*

##### Genes associated with SpatialPC2

**Positively correlated**

Smim5, Slc1a3, Mgll, Mgl2, Mfsd7a, Cd163l1, Cd14, Sh2d1b1, Met, Mertk, Mefv, Fcgr3, Siglec1, Fcgr4, Sirpb1b, Sirpb1c, Fcrl5, Ska1, Ffar2, Mcemp1, Ffar4, Slamf8, Slamf9, Slc11a1, Slc16a9, Mgst1, Serpinh1, Fcgr1, Sct, Msr1, Cd36, Sash1, Cd33, Cd302, Cd300ld, Ms4a8a, Ms4a7, Scn3a, Cd300lb, Cd300c, Mmp19, Fam149a, Fblim1, Cd209d, Sept3, Mreg, Mrc1, Mmp23, Cd209a, Serpinb8

Phagosome, Hematopoietic cell lineage, Malaria, Acyl chain remodeling of diacylglycerol and triacylglycerol, Antigen processing: cross presentation, Cross-presentation of particulate exogenous antigens (phagosomes), TSP1-induced apoptosis in microvascular endothelial cell, Sema4D-mediated inhibition of cell attachment and migration, IKK complex recruitment mediated by RIP1, Hormone-sensitive lipase (HSL)-mediated triacylglycerol hydrolysis

**Negatively correlated**

Tcf7, Rps12, Pip4k2a, Dapl1, Themis, Rplp0, Rpl41, Itgal, Cd8b1, Cd8a, Dap, Evi2a, Slc25a4, Gapdh, Nsg2, Cd9, Tnfsf11, Ppic, Atpif1, Rpsa, Pgk1, Cd200, Ptpn22, Crtam, Cd160, Ccnb2, Nkg7, Gpm6b, Smtn, Tbc1d4, Actn1, Xcl1, Fyn, Cd3d, Sh2d1a, Scd2, Ddt, Rgcc, Msrb1, Zyx, Txn1, 2310001H17Rik, Mxd4, Thy1, Cst7, Ube2c, Serpinb6b, Lag3, Casp3, Vdac1

T helper cell surface molecules*, Immunoregulatory interactions between a lymphoid and a non-lymphoid cell*, Interleukin-2 signaling pathway*, Interleukin-4 regulation of apoptosis*, T cell receptor regulation of apoptosis*, Adaptive immune system*, Cytoplasmic ribosomal proteins*, TSP1-induced apoptosis in microvascular endothelial cell*, Influenza viral RNA transcription and replication*, T cell receptor signaling in naive CD8+ T cells*

cDC1s

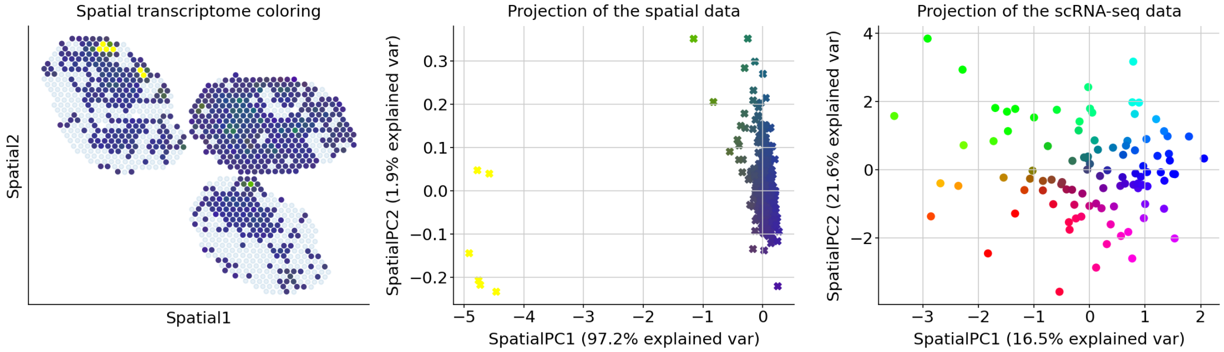


#### Genes associated with SpatialPC1

##### Positively correlated

Itgb8, Spon1, Klra17, Klra1, Klk1b27, Csf3r, Sned1, P2ry13, Kdr, Kcnj10, Kcnc1, Izumo1r, Sox13, Cd160, Sox5, Prkar1b, Sparc, Palld, Itga9, Atp2a1, Klra2, Atp2b4, Klrc1, Crmp1, Lcn4, Crtam, Ppp1r3d, Cd163l1, Cryba4, Klrg1, Klrb1c, Klra4, Klrb1a, Klra9, Klra8, Klra7, Slfn4, Slpi, Klra6, Itga2, Plagl1, Lifr, Il7r, Ifng, Aqp9, Ifitm10, Ptprm, Tcrg-C1, Tcrg-C2, Tcrg-C4

Integrin family cell surface interactions*, Jak-STAT signaling pathway, Leptin influence on immune response, Immunoregulatory interactions between a lymphoid and a non-lymphoid cell, Reduction of cytosolic calcium levels, Cytokine-cytokine receptor interaction, Arrhythmogenic right ventricular cardiomyopathy (ARVC), Platelet calcium homeostasis, ECM-receptor interaction, Integrin cell surface interactions

##### Negatively correlated

Stmn1, Ube2c, Birc5, Aurkb, Top2a, Cdca8, Nusap1, Kif11, Hmgb2, Cks1b, Mki67, Ccnf, Ptma, Pbk, Spc24, Hist1h4d, Tk1, Hist1h2ae, Bub1b, Plk1, Cdk1, Prc1, Incenp, Tubb4b, Cenpl, Ndc80, Ccnb1, Cit, Aurka, Rrm2, Tacc3, Cdca3, Kif15, Ttk, Cdc25b, Esco2, Rfc5, Cks2, Asf1b, Cdca2, Bub1, Cenpm, Cdc20, Smc2, Lockd, Hjurp, Hist1h3c, Mxd3, Melk, Fbxo5

Cell cycle*, Polo-like kinase 1 (PLK1) pathway*, M phase pathway*, DNA replication*, Aurora B signaling*, APC/C-mediated degradation of cell cycle proteins*, Phosphorylation of Emi1*, FOXM1 transcription factor network*, APC/C activator regulation between G1/S and early anaphase*, APC/C- and Cdc20-mediated degradation of Nek2A*

#### Genes associated with SpatialPC2

##### Positively correlated

Gsta4, Thbd, Klrc1, Nid1, Gm12253, Klrg1, Gm12972, Cd160, Ngfr, Eomes, Adgre4, Enpp2, Cald1, Serpinb9b, Thbs1, Gm14085, Notch3, Serpinh1, Gm1673, Nebl, Neb, Ar, Lcn4, Ccr4, Aqp9, Clec4g, Tdrp, Ndnf, Tcrg-C4, Tcrg-C2, Cdkn2a, Klrb1c, Sparc, Klk1b27, Gdpd1, Ryr3, Actg2, Cma1, Dclk1, Actn2, Nrxn1, Samd3, Hcar2, Cadm3, Etl4, Hbegf, Cd163l1, Tmem26, Klrb1a, Klra1

Muscle contraction*, ID regulation of gene expression, Response to elevated platelet cytosolic calcium, Smooth muscle contraction, Senescence and autophagy, Immunoregulatory interactions between a lymphoid and a non-lymphoid cell, FRA pathway, Striated muscle contraction, Bladder cancer, Gastrin pathway

**Negatively correlated**

Psap, Ece1, Cd74, Rrbp1, Gns, H2-Ab1, Poglut1, Hsp90b1, Adam8, Slamf7, Mpeg1, Havcr2, Neat1, Ifngr1, Clec9a, Anpep, Procr, Selplg, Csf2rb, Xcr1, Slc8b1, Zfp516, Fgl2, Myadm, Sbno2, H2-Eb1, Camk1d, Pdia3, Ppt1, Trpm2, Herpud1, Lyn, Flt3, Ahr, Nabp1, Tlr11, P2ry6, Tbc1d8, Itm2c, Ly75, Nr4a1, Rp2, Tbc1d9, P2ry14, Naga, Pik3cb, Sdc3, Zfp366, Icam1, Ehbp1l1

Interleukin-2 signaling pathway*, Alpha-M beta-2 integrin signaling, Interleukin-3, interleukin-5, and GM-CSF signaling, Lysosome, Adhesion and diapedesis of granulocytes, Immune system signaling by interferons, interleukins, prolactin, and growth hormones, Nucleotide-like (purinergic) G-protein coupled receptors, Immune system, Protein processing in the endoplasmic reticulum, Cells and molecules involved in local acute inflammatory response

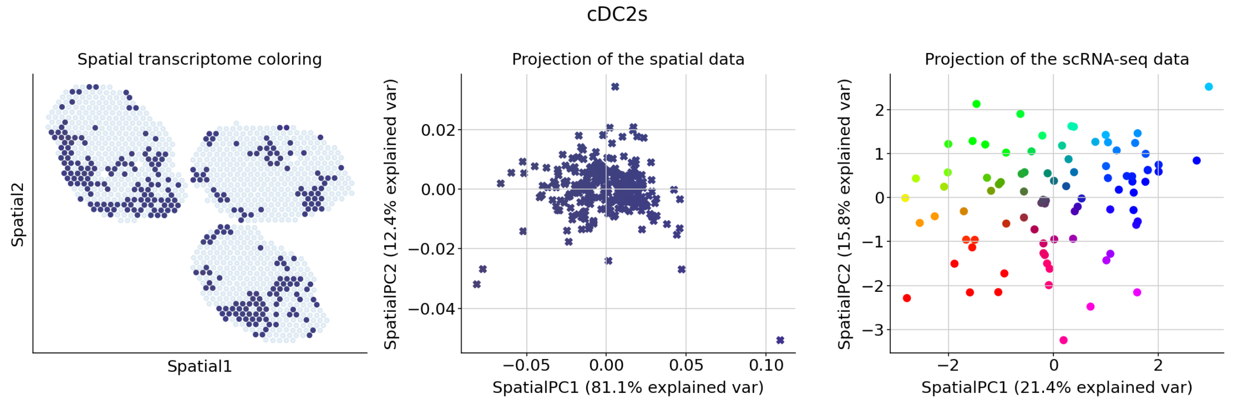


#### Genes associated with SpatialPC1

**Positively correlated**

Tcrg-C4, Fasl, Fam83d, Arl4d, Lingo4, Lmo7, Prg4, Tmem26, Etl4, Prg2, Ltc4s, Asns, Clec4e, Cald1, Tlr3, Erbb2, Ppef2, Enpp2, Popdc2, Mab21l3, Map2, Mdk, Cma1, Arhgap20, Fblim1, Plekhg1, Tnfrsf8, Ptpn13, Klra4, Klra6, Klra7, Klra8, Klra9, Klrb1c, Klrc1, Klrc2, Ptgs1, Stab1, Slc16a9, Cbr2, Slc1a3, Tnnt2, Psd2, Cited4, Apoc1, Apol10b, Laptm4b, Tnfrsf9, Plscr4, Mgll

Eicosanoid biosynthesis, Ras-independent pathway in NK cell-mediated cytotoxicity, T cell receptor regulation of apoptosis, Muscle contraction, Arachidonic acid metabolism, GRB7 events in ERBB2 signaling, Acyl chain remodeling of diacylglycerol and triacylglycerol, TGF-beta regulation of extracellular matrix, Adherens junction cell adhesion, Antigen processing and presentation

**Negatively correlated**

Malat1, Sirpa, Fyb, Gm42418, St8sia6, Trps1, Tnfrsf1b, Inpp4b, Tcf7, Gda, Cd5, Myof, Pld4, Ddhd1, Neat1, Ptpn22, Lilrb4a, Nfkbiz, Anxa3, Txk, Tspan33, Epb41l2, Stat1, Pdlim4, Pla2g7, Cxcl16, Cd247, Birc3, Sdc4,

Skap1, H2-Eb1, Lck, Trac, Gbp8, Lcp2, Dnase1l3, Ly75, Fam49a, Itgax, Hist1h1c, Rel, Tarm1, Cdk14, Ehf, Cd3d, Ptafr, Cd28, Ms4a4b, Gbp4, Lef1

Generation of second messenger molecules*, Interleukin-2 signaling pathway*, T cell receptor signaling in naive CD4+ T cells*, T cell activation co-stimulatory signal*, T cell receptor signaling in naive CD8+ T cells*, Costimulation by the CD28 family*, PD-1 signaling*, HIV-induced T cell apoptosis*, Interleukin-5 regulation of apoptosis*, Lck and Fyn tyrosine kinases in initiation of T cell receptor activation*

##### Genes associated with SpatialPC2

###### Positively correlated

Gdpd1, Klrc1, Mdk, Efnb3, Lingo4, Dntt, Tnfrsf9, Tnfrsf8, Ncr1, Ncam1, Ccr10, Dnm3, Mgst1, Dll1, Plekhg1, Klrc2, Hpn, Cd163l1, Tmem26, Prg2, Mgll, Hp, Sox13, Enpp2, Tnnt2, Klrb1c, Klra7, Cald1, Trem2, Trdc, Crmp1, Erbb2, Plscr4, Gpx3, Igfbp7, Nebl, Klra8, Klra9, Gramd2, Neb, Cbr2, Ifitm10, Ccl12, Ccl17, Rorc, Src, Gsta4, Ptgs1, Cma1, Klrg1

Axon guidance*, Muscle contraction, Glutathione metabolism, Developmental biology, Semaphorin interactions, Ras-independent pathway in NK cell-mediated cytotoxicity, L1CAM interactions, Glutathione conjugation, Recycling pathway of cell adhesion molecule L1, Interleukin-7 interactions in immune response

###### Negatively correlated

Pld4, Ppt2, Tgfbi, Il1rn, Fbxo5, Lst1, Lyz2, Ube2c, Dlgap5, Tspan33, Birc5, Cdca3, Cd63, Hmox1, Pla2g7, Plxnd1, Bub1b, Pltp, Clec4a3, Mmp12, Cdk1, Clec4a1, Slc7a11, Mxd3, Cd81, Gas2l3, Gngt2, Sat1, Psap, E2f2, Emb, Nuf2, Hist1h2ae, Ccl6, Hck, Nr1h3, Depdc1a, Ccnb2, Tpi1, Slamf8, Sirpb1b, Siva1, Ear2, Fcgr3, Pid1, Mki67, Kif4, Cxcl16, Ckap2, Myof

Cell cycle*, APC/C activator regulation between G1/S and early anaphase*, APC/C- and Cdc20-mediated degradation of Nek2A*, APC/C-mediated degradation of cell cycle proteins*, Cyclin B2-mediated events*, Phosphorylation of Emi1*, M phase pathway*, FOXM1 transcription factor network*, DNA replication*, Polo-like kinase 1 (PLK1) pathway*

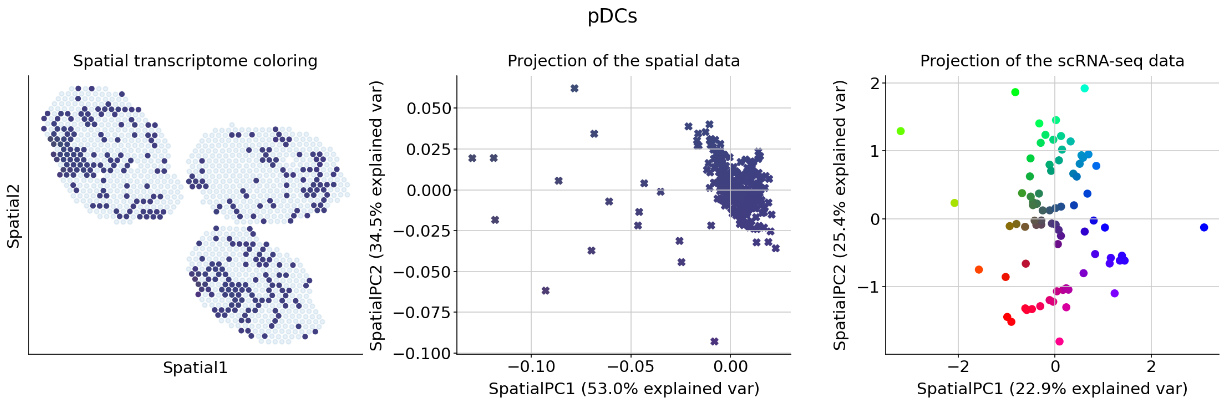


##### Genes associated with SpatialPC1

**Positively correlated**

1300017J02Rik, Hist1h2ag, Htr7, Samd3, Hpn, Hpgd, Sapcd2, Hp, Sash1, Hmmr, Hmgb3, Hlf, Hk3, Scin, Hist1h3c, Hist1h2bj, Hepacam2, Enpp2, Hcar2, Hbegf, Hbb-bt, Hap1, H1fx, Gzmm, Gzmc, Gzmb, Gzma, Gtse1, Serpinb8, Gsta4, Serpinb9b, Serpine2, Htra1, Ryr3, Ifit3b, Ifitm1, Jag1, Itgb8, Itgae, Rgs16, Itga9, Rgs9, Itga8, Itga2, Itga1, Iqgap3, Il4, Rnf144b, Il1rl1, Il1r1

Integrin family cell surface interactions*, ECM-receptor interaction*, Integrin cell surface interactions*, Actin cytoskeleton regulation*, Integrin-mediated cell adhesion*, Plexin D1 signaling*, Arrhythmogenic right ventricular cardiomyopathy (ARVC)*, Dilated cardiomyopathy*, Arf6 integrin-mediated signaling pathway*, Interleukin-12-mediated signaling events*

**Negatively correlated**

Cybb, Ifi204, Grn, Cyb561a3, Lair1, Mpeg1, Tcf4, Xbp1, B2m, Melk, Cbr3, Metrnl, Trpm2, Cd14, Tnfsf12, Gpr141, Guca1a, E2f2, Ccl17, Tgm2, Klra2, Bambi, 1700025G04Rik, 2610528A11Rik, Prdm1, Fcgr3, Il1b, Msr1, Rtn1, 1600010M07Rik, Cd300lb, Ffar2, Slc7a11, Pid1, Pilrb2, Cd300ld, Ip6k3, Tmem51, Sirpb1b, Sirpb1c, C3, Treml4, Tgfbi, Thbs1, Csf1r, Celf4, Clec4n, Ms4a7, Clec4a1, Clec4a2

Beta-1 integrin cell surface interactions*, Phagosome*, Wnt interactions in lipid metabolism and immune response, Hematopoietic cell lineage, Fibroblast growth factor 1, Integrin beta-2 pathway, Cytokine-cytokine receptor interaction, Bladder cancer, Beta-3 integrin cell surface interactions, Type II interferon signaling (interferon-gamma)

##### Genes associated with SpatialPC2

**Positively correlated**

1300017J02Rik, Ifitm10, Il18bp, Il10, Ikzf4, Ighg2b, Igha, Igfbp4, Igf1, Serpinb8, Ifnlr1, Serpinb9b, Serpine2, Ifng, Ifitm6, Ifitm1, Fignl1, Ifit3b, Htra1, Htr7, Hpn, Hpgd, Hp, Hmmr, Hmgb3, Hlf, Hk3, Sidt1, Siglec1, Hist1h3c, Il18r1, Il18rap, Il1r1, Il1rl1, Klra3, Klra1, Kif4, Kif2c, Kif20a, Kif18b, Kif14, Ryr3, Kif11, Khdc1a, Zmynd15, Kcnj2, Samd3, Kcnj10, Sapcd2, Sash1

Interleukin-12-mediated signaling events*, Th1/Th2 differentiation pathway*, Kinesins*, Cytokine-cytokine receptor interaction*, Interleukin-23-mediated signaling events*, Interleukin-12/STAT4 pathway*, Ghrelin-mediated regulation of food intake and energy homeostasis, Dendritic cells in regulating TH1 and TH2 development, Insulin-like growth factor (IGF) activity regulation by insulin-like growth factor binding proteins (IGFBPs), Hypertrophy pathway

**Negatively correlated**

Actb, Isg15, Slfn5, Phf11b, Oasl2, Ifi44, Ifitm3, Usp18, Tagln2, Sdc3, Atp5g3, Cmpk2, Psme2, Parp14, Rasa4, Pkib, Actg1, Atp6v0c, Iigp1, Cd7, Ifi47, Evi2a, Fam129a, Pfkp, Dnajc7, Lgals1, Hnrnpab, Irf7, Calm1, Gm42418, Casp3, Acer3, Rtp4, Prkcd, Plac8, Clec9a, Crip1, Atp5g1, Uqcrq, Rsad2, Atad5, AY036118, Skap2, Trafd1, Eif2ak2, Zbp1, Taldo1, Ass1, Rnh1, Cd68

Interleukin-2 signaling pathway*, Interferon signaling*, Interferon alpha/beta signaling*, Myometrial relaxation and contraction pathways*, Alzheimer's disease*, Immune system signaling by interferons, interleukins, prolactin, and growth hormones*, Type II interferon signaling (interferon-gamma), Parkinson's disease, Vibrio cholerae infection, Oxidative phosphorylation

### Supplementary Note 2: DestVI report for the tumor data

#### Step A: automatic thresholding

CD4-Tregs: critical points

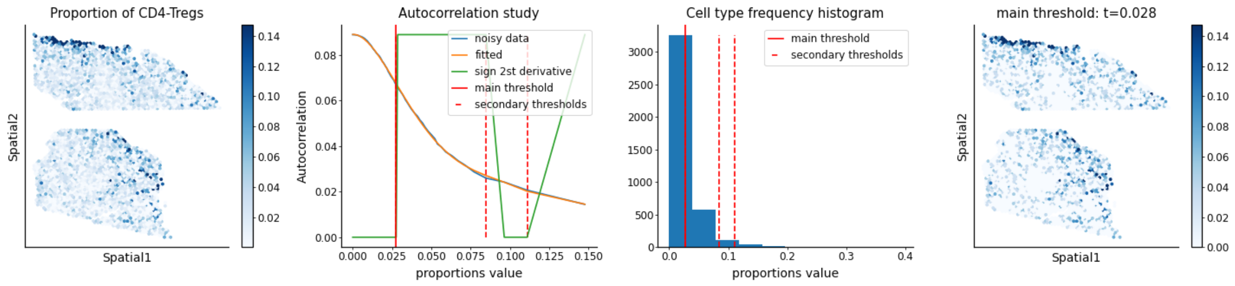


CD8: critical points

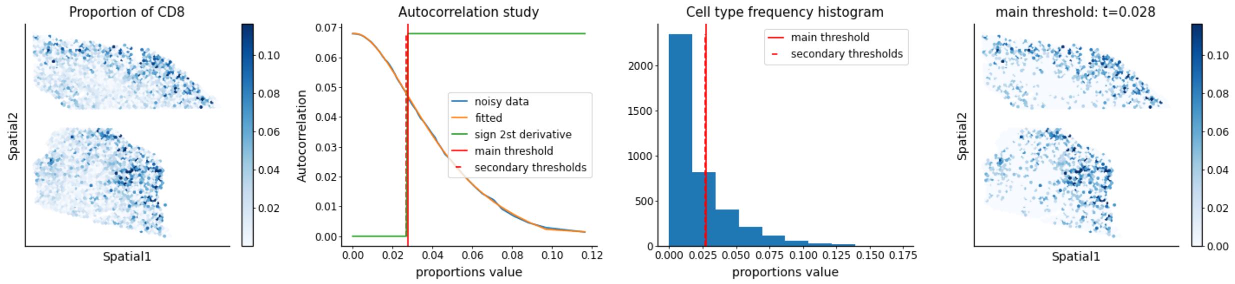


DCs: critical points

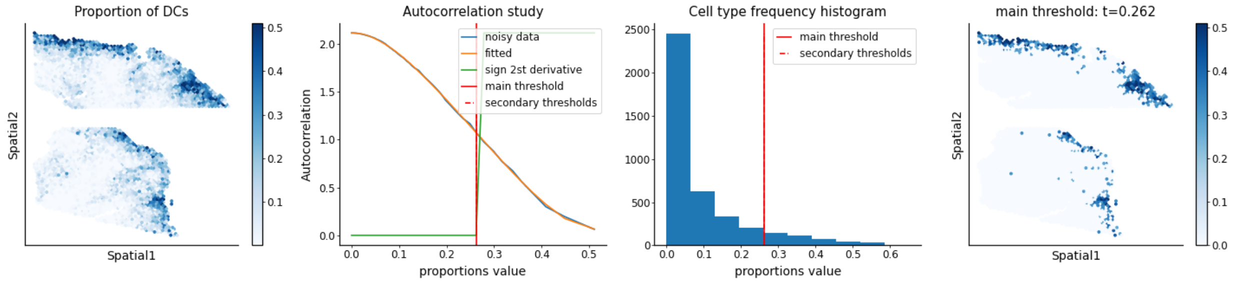


Mon-Mac: critical points

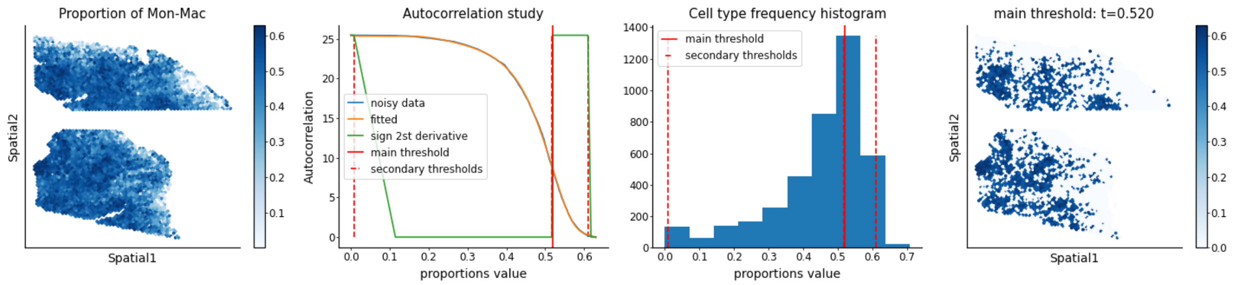

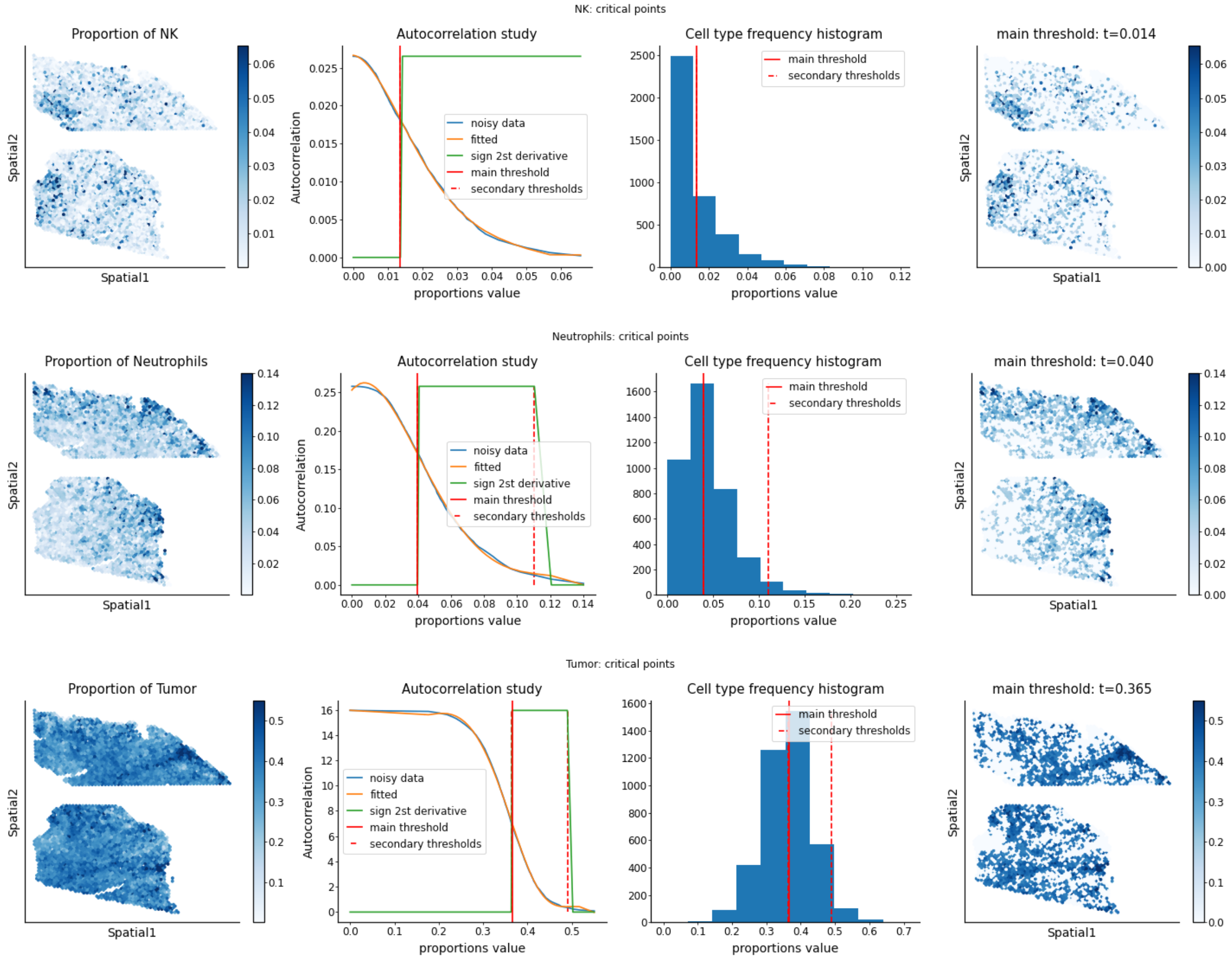


### Step B: weighted PCA and GSEA

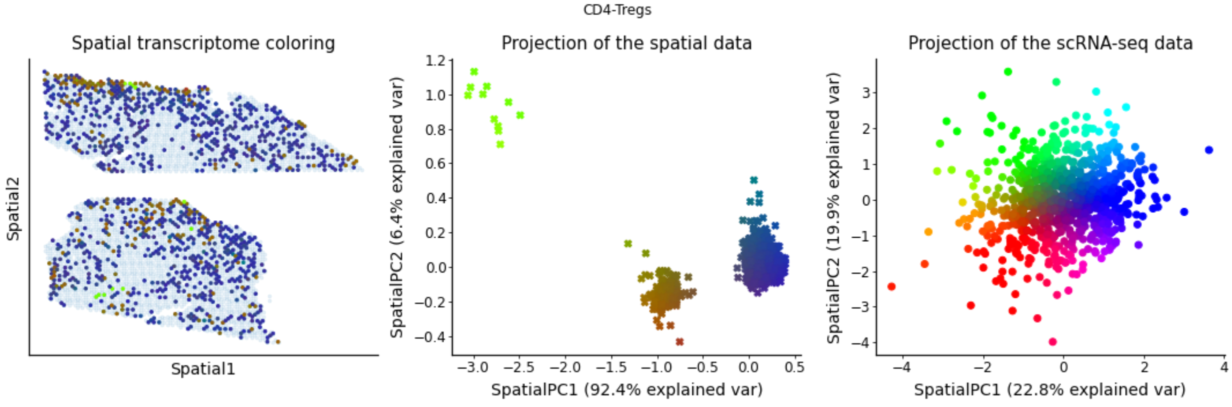


**Genes associated with SpatialPC1**

**Positively**

Fgf7, Runx2os2, Gli3, Glrp1, Gm10244, Gm10501, Rslcan18, Gm12689, Gm12708, Gm13056, Gm13199, Gm13391, Gm13470, Gm13496, Gm13713, Rps13, Gm14010, Gm14154, Gm14207, Gm14636, Gm15056, Gm15518, Gm15952, Gjc1, Gjb4, Gm16096, S100a9, Scx, Sctr, Gal, Galnt18, Scrn1, Gas6, Gata2, Schip1, Scd1, Scamp5, Sbsn, Saxo2, Gchfr, Gcsam, Gdf15, Samd4, Samd14, Gemin4, Saa3, Ghr, S100g, Gm15956, Rnft2

Gap junction assembly, Gap junction trafficking and regulation, BDNF signaling pathway, Endochondral ossification, Electric transmission across gap junctions, FGFR2b ligand binding and activation, Tetrahydrobiopterin (BH4) biosynthesis, recycling, salvage and regulation, Gamma-carboxylation, transport, and amino-terminal cleavage of proteins, Glypican-3 pathway, Prolactin receptor signaling pathway

**Negatively**

Nhp2, Rpl4, Rpl8, Serbp1, Banf1, Cks1b, Npm1, Dut, Hsp90ab1, Hmgb2, Atp5a1, Hint1, Hnrnpab, Stmn1, Birc5, Cxcr6, Pdlim1, Hist1h2ao, Rgcc, Cdca3, Cct6a, Smc2, Ranbp1, Ncl, Cct5, Cct8, Snrpa1, Eef1d, Snrpd1, Cenpf, Cct7, Dkc1, Lsm4, C1qbp, Hspe1, Fkbp4, Dpy30, Ccnb2, Ybx3, Trbc2, Cd7, Adh5, Hspd1, Hist1h2ae, Tyms, Mrpl12, Eif2s2, Gars, Cd9, Cct3

Folding of actin by CCT/TriC*, Cooperation of prefoldin and TriC/CCT in actin and tubulin folding*, Association of TriC/CCT with target proteins during biosynthesis*, Protein metabolism*, Aurora B signaling*, Protein folding*, FOXM1 transcription factor network*, Cell cycle*, Myc active pathway*, Pyrimidine biosynthesis*

**Genes associated with SpatialPC2**

**Positively**

Rnf183, Scx, Cxcl1, Kcng2, Nupr1l, Sectm1a, Cxcl14, Sdsl, Cxcl5, Kcne3, Gm44053, Sctr, Sgcd, Cxcr2, Scrn1, Mirt2, Gm42517, Cyp1b1, D130017N08Rik, Schip1, D130020L05Rik, Scd1, Gm44275, Sema3a, Sema6a, Ctsf, Colec12, Gm8773, Copz2, Gm826, M1ap, Gm807, Serpine1, Creb3l3, Gm568, Kcnh2, Serpinb2, Nuak1, Cryab, Cspg5, Cstad, Gm4890, Sept5, Cth, Gm45051, Scamp5, Sbsn, Saxo2, Rslcan18, Dmrta1

TGF-beta regulation of extracellular matrix*, Senescence and autophagy*, Dissolution of fibrin clot*, Binding of chemokines to chemokine receptors, Fibrinolysis pathway, Oncostatin M, Chemokine signaling pathway, Blood clotting cascade, TSH regulation of gene expression, Fibroblast growth factor 1

**Negatively**

Gm42418, Gm26917, AY036118, Serpinb9, Sdf4, Tnfrsf4, Tg, Nfkbia, Tnfrsf18, Rel, Lgals7, Hilpda, Mir155hg, Serpinb6b, Gadd45b, Trac, Tnfrsf9, Cd200, Dusp2, Cd2, Il5, Zap70, Gm19585, Arap2, Cdkn1a, Icos, Dusp4, Csf2, Sh2d1a, Ms4a4b, Rilpl2, Lck, Tnfsf11, Cd6, Cyp51, Gimap5, Cct8, Ak2, Gimap6, Sdcbp2, Bhlhe40, Cd4, Zbtb32, Ifitm1, Eprs, Rgs16, Traf1, Camk4, Acot7, Sqstm1

T cell receptor signaling pathway*, T cell receptor regulation of apoptosis*, Immune system*, Interleukin-2 signaling pathway*, PD-1 signaling*, Primary immunodeficiency*, Generation of second messenger molecules*, Adaptive immune system*, Lck and Fyn tyrosine kinases in initiation of T cell receptor activation*, Hematopoiesis regulation by cytokines*

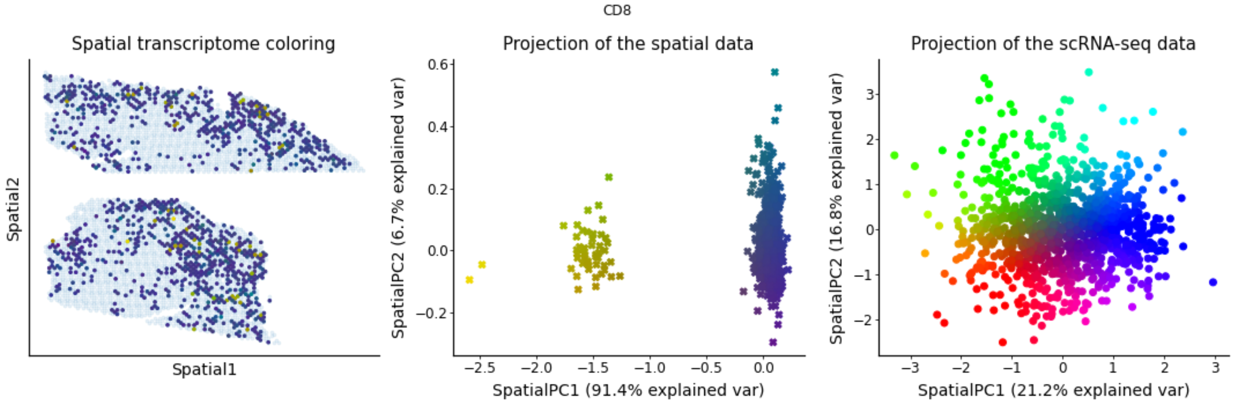


##### Genes associated with SpatialPC1

###### Positively

Prss27, Plpp7, Cyp1b1, Pou3f1, Postn, Cyp2r1, Cyp7b1, D330050G23Rik, D430040D24Rik, Dach1, Plxna4os1, Plxna4, Plpp3, Ppbp, Pllp, Ddr2, Plet1, Plekha4, Plcl1, Platr3, Plagl1, Pla2r1, Dhrs9, Pla2g4f, Pparg, Ppef2, Pkia, Ctsf, Ptn, Ptk2, Ptgs2, Creb3l3, Crispld2, Cryba4, Csgalnact1, Prtn3, Prss50, Prr15, Prkg2, Cxcr2, Prelp, Prdm5, Cxadr, Ppp1r27, Ppp1r14c, Ppp1r14a, Ppp1r13l, Cxcl14, Cxcl3, Cxcl5

Chemokine signaling pathway*, Binding of chemokines to chemokine receptors*, Phase I of biological oxidations: functionalization of compounds*, TGF-beta regulation of extracellular matrix*, Interleukin-7 interactions in immune response*, Interleukin-1 regulation of extracellular matrix*, Biological oxidations, Cytochrome P450 pathway, Cytokine-cytokine receptor interaction, Cytochrome P450 metabolism of endogenous sterols

###### Negatively

Isg20, Ifit3, Usp18, Bst2, Ly6a, Gzmb, Rsad2, Ifit1, Ifit1bl1, Igtp, Gbp2, Irgm1, Ifit3b, Jaml, Ifi208, Ifit2, Ifih1, Slfn5, Phf11c, Cxcl10, Cmpk2, Plac8, Cd8a, Ifi214, Tnp2, Iigp1, Socs1, Slamf7, Tnfrsf9, Cd69, Samd9l, Hspa5, Mcm2, AW112010, Tuba1b, Cxcr6, Fen1, Ly6c2, Mcm6, Ctla2a, Il12rb1, Spc24, Ccrl2, Pmepa1, Sh2d2a, Top2a, Lig1, Cd8b1, Lag3, Ctsw

Interferon alpha/beta signaling*, Interferon signaling*, DNA strand elongation*, Immune system signaling by interferons, interleukins, prolactin, and growth hormones*, Interleukin-12-mediated signaling events*, Interleukin-4 regulation of apoptosis*, Immune system*, DNA replication*, S phase*, Type II interferon signaling (interferon-gamma)*

##### Genes associated with SpatialPC2

###### Positively

Nrbp2, Dhx58os, Slc9a4, Gpha2, Gpc6, Nrep, Ccdc151, Ccdc142os, Aplnr, Slc6a17, Slc6a13, Ccdc106, Gng11, Slc4a3, Nuak1, Slc40a1, Slc39a2, Cbr2, Apol6, Gfra2, Anxa8, Npy, Gpr137c, Snhg18, Ankrd33b,

Ankrd35, Nid2, Sncg, Gdf3, Gpr153, Gpnmb, Npl, Ccl11, Ano1, Smim5, Nos2, Gpihbp1, Smad6, Antxr1, Obsl1, Slc27a3, Gnao1, Efna5, Shroom4, Shroom3, Capns2, Gm9530, Pak3, Paqr5, Parva

SLC-mediated transmembrane transport, Transmembrane transport of small molecules, Metal ion solute carrier family (SLC) transporters, Transport of glucose and other sugars, bile salts and organic acids, metal ions and amine compounds, G-protein activation, Inhibition of insulin secretion by adrenaline/noradrenaline, Oncostatin M, Peptide G-protein coupled receptors, G alpha (i) signaling events, Opioid signaling

**Negatively**

Gzma, Ifit1, Gzmb, Ifit3, Slfn5, Ccl5, Serpinb9, Igtp, Rsad2, Ifit1bl1, Ifit3b, Mxd1, Socs1, Ly6c2, Serpinb6b, Usp18, Ifi208, Ccnd2, Ifih1, Bst2, Gbp2, Isg20, AW112010, Oasl1, Klra9, Klrk1, Cmpk2, Samd9l, Tcrg-C4, Iigp1, Klre1, Ifi214, Klrb1c, Tcrg-C2, Ccrl2, Gm34342, Ly6c1, Kcnj8, Fasl, Klra1, Tmem140, Arhgef5, Cish, Sult2b1, Klra5, Dgat1, Irgm1, Jaml, Ifit2, Zfp628

Interferon alpha/beta signaling*, Interferon signaling*, Immune system signaling by interferons, interleukins, prolactin, and growth hormones*, Interleukin-9 regulation of target genes*, Immune system*, Granzyme A-mediated apoptosis pathway*, Interleukin-4 regulation of apoptosis, Interleukin-12-mediated signaling events, Interleukin-2 signaling pathway, Interferon alpha signaling regulation

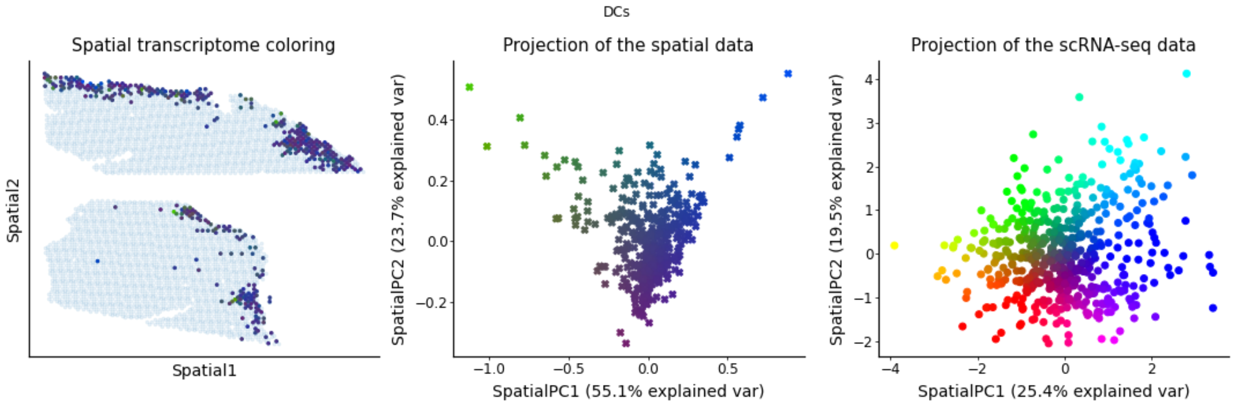


#### Genes associated with SpatialPC1

**Positively**

Gm10505, Ddx43, Dixdc1, Armcx4, Dhx58os, Lrrc73, Serping1, Lurap1, Serpine2, Lurap1l, Arnt2, Serpinb2, Lyzl4, Arsg, Ly6g5b, Asb11, Serp2, Dcbld1, Sept5, Asb4, Lyve1, Dapl1, Loxl4, Sgcd, Dlg5, Sgip1, Lcn4, Arhgef5, Skap1, Dnmt3l, Six2, Lgals7, Sit1, Lhx6, Shroom4, Shroom3, Shox2, Lif, Shisa4, Shf, Lmcd1, Dnah8, Dmrta1, Lmntd2, Dmpk, Sema6a, Dach1, Lcn2, Cspg5, B230312C02Rik

Dissolution of fibrin clot, Dermatan sulfate biosynthesis, Glycosaminoglycan metabolism, Arylsulfatase activation, Chondroitin sulfate/dermatan sulfate degradation, Hyaluronan metabolism, Prolactin regulation of apoptosis, Interleukin-1 regulation of extracellular matrix, Coagulation intrinsic pathway, Fibrinolysis pathway

**Negatively**

Ifit3, Ifit1, Ifit2, Rsad2, Isg20, Ifit3b, AA467197, Phf11d, Ly6a, H2-Ab1, Cxcl10, Axl, H2-DMb1, Ifit1bl1, Cd74, H2-Aa, Tgm2, Samd9l, Igtp, H2-Eb1, Irgm1, Cd72, Oasl1, Pnp, Usp18, Cmpk2, Ifih1, Gbp2, Ifi208, Cd14, Ccnd2, Ifi44, Slfn5, Il1rn, Sectm1a, Ifi205, Spint1, Socs1, AU020206, Slfn4, Slfn9, Iigp1, Cdkn1a, Clec4n, Ccrl2, Slamf9, Ctsd, Rilpl1, AW112010, Hexb

Interferon alpha/beta signaling*, Interferon signaling*, Immune system signaling by interferons, interleukins, prolactin, and growth hormones*, Immune system*, Type II interferon signaling (interferon-gamma), Cyclins and cell cycle regulation, Interferon alpha signaling regulation, Leptin influence on immune response, Cyclin D-associated events in G1, Interleukin-4 regulation of apoptosis

##### Genes associated with SpatialPC2

**Positively**

Rbp2, Fam160a1, Endou, Enpp2, Prickle1, Prf1, Prelp, Prdm5, Eomes, Epas1, Ppp1r14c, Epb41l4a, Epb41l5, Ephb3, Ephx3, Epn2, Epor, Ereg, Etv1, Evc, Evc2, Pou6f1, Eya2, F2rl1, F2rl3, Pomc, F630040K05Rik, Prm3, Prox2, En1, Ecm2, E2f7, Ptprn, Ptprk, Ptprg, Ptpn5, Eaf2, Ptpn13, Ptpdc1, Ebf3, Ptn, Echdc2, Eda2r, Emx2, Efcab9, Efhc1, Efnb3, Ehd3, Prx, Elmo3

Type I interferon (interferon-alpha/beta) pathway*, Jak-STAT signaling pathway*, Ephrin receptor B forward pathway, TGF-beta regulation of extracellular matrix, Syndecan 3 pathway, Steroid hormone, vitamin A, and vitamin D metabolism, Type 1 diabetes mellitus, Peptide G-protein coupled receptors, CD8/T cell receptor downstream pathway, Glucocorticoid biosynthesis

**Negatively**

H2-DMb1, H2-Ab1, Rpl4, Lgals1, Cd209a, Rpl8, Cd74, Ckb, Naaa, Anxa1, Wfdc17, Ranbp1, Ccl6, Itgb7, Lyz2, Calr, Nop10, Clec10a, Nhp2, H2-Aa, Eif5a, Selenop, Apex1, Cd7, Ighm, H2-Eb1, Mgst3, Eif4ebp1, Ccl9, Egr1, Hnrnpab, Ramp1, Nop16, Hint1, Ifitm6, Abca9, Cyb5a, Sell, Plac8, Kmo, Bcl11a, Cdkn2d, Parvg, Cd209d, Ank, H13, Nop58, Atp5a1, Mrc1, Mcm6

Protein metabolism, Interleukin-9 regulation of target genes, T cell receptor regulation of apoptosis, Adaptive immune system, Influenza infection, Translation, EGFR1 pathway, Translation factors, Immune system, G1 to S cell cycle control

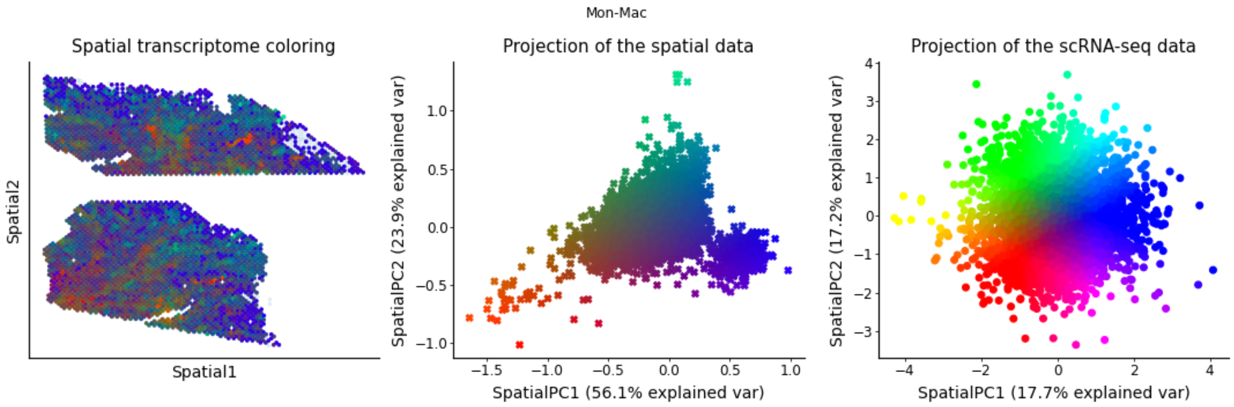


**Genes associated with SpatialPC1**

**Positively**

Perp, Utf1, Klra1, Serp2, Klra5, Dapl1, Gramd2, Tmem150c, Apol10b, B930095G15Rik, Lcn4, Ypel4, Chrd, Chst2, Nefh, Lurap1, Gpha2, Tff1, Gng3, Fam71b, Cd163l1, Fgf21, Gjb4, Ptprg, Hemgn, Upk1a, A630023P12Rik, Scin, Arnt2, Dmrta1, Pdzd4, Ppp1r14c, Il5, Kcnj8, Ninj2, Endou, 1700001O22Rik, Sema6d, Aqp3, Gzmf, Bmp7, Cfap69, Dzip1, Lancl3, Nckap5, Extl1, Platr22, 2700069I18Rik, Gm11714, Sv2c

Inwardly rectifying potassium channels, ALK in cardiac myocytes, BMP receptor signaling, Aquaporin-mediated transport, Endochondral ossification, TGF-beta regulation of skeletal system development, Eosinophils in the chemokine network of allergy, Potassium channels, Passive transport by aquaporins, ALK2 pathway

**Negatively**

Gm42418, Gm26917, Thbs1, Pf4, AY036118, Cd93, F13a1, Abca1, Ctsl, Ms4a7, Emp1, Cxcl3, Apoe, Adam8, Spp1, Hilpda, Ctsd, Folr2, Mafb, Nceh1, Ppbp, Rnase4, Pmp22, Ninj1, Syngr1, Fcrls, Arg1, Gpnmb, Mxd1, Stab1, Ccl9, Fabp5, Rgs1, Cxcl2, Ccl7, Tcf4, Ms4a14, Tspan3, Ccl2, Tmem37, Ecm1, Gstm1, Filip1l, Ctla2b, Cbr2, Olfml3, Ddah2, Rsad2, Sox4, Hmox1

Binding of chemokines to chemokine receptors*, Interleukin-4 regulation of apoptosis*, Interleukin-1 regulation of extracellular matrix*, Chemokine signaling pathway*, Peptide G-protein coupled receptors*, Oncostatin M*, Alpha-9 beta-1 integrin pathway*, Cytokine-cytokine receptor interaction*, Response to elevated platelet cytosolic calcium*, Interleukin-7 interactions in immune response*

**Genes associated with SpatialPC2**

**Positively**

Lurap1, Arnt2, Nrtn, 2700069I18Rik, Fgf21, Klra1, Klra5, Mid2, Fam71b, Chst2, Lancl3, Tmem150c, Dzip1, Lcn4, Extl1, Serp2, Ninj2, B930095G15Rik, Nefh, Ypel4, Aqp3, Nckap5, Ncald, Nbl1, Il5, Naalad2, D630039A03Rik, Upk1a, Dmrta1, Shroom3, Muc20, Utf1, Apol10b, Gjb4, Gm11714, Dapl1, Slc6a17,

Ppp1r14c, Cd79a, Gzmf, Scin, Ephb3, Gramd2, Gpha2, 1700001O22Rik, Gng3, Cd163l1, Ptprg, A630023P12Rik, Endou

Activation of kainate receptors upon glutamate binding, Aquaporin-mediated transport, Eosinophils in the chemokine network of allergy, Activation of calcium-permeable kainate receptor, Passive transport by aquaporins, Glycosaminoglycan biosynthesis: keratan sulfate, Hematopoiesis regulation by cytokines, Dendritic cells in regulating TH1 and TH2 development, Gap junction assembly, GATA3-mediated activation of Th2 cytokine expression

**Negatively**

Ccl6, Lyz2, Pf4, Ccl9, F13a1, Fn1, Cd14, Cxcl2, Thbs1, Lyz1, Cd93, Hp, Ctsl, Cxcl3, Ctsd, Mt1, Ppbp, Clec4n, Rpl4, Ninj1, Hilpda, Lmna, Vat1, Rpl8, Spp1, Fam20c, Cd36, Igfbp4, Selenop, Pdlim1, Cbr2, Akr1b8, Emp1, Nhp2, Folr2, Mcemp1, Sdc1, Mmp8, Mrc1, Cd9, Areg, Ear2, Gja1, Slc7a11, Igfbp6, Clec4e, Abca1, Gcsh, Klf2, Tspan3

Response to elevated platelet cytosolic calcium*, Beta-1 integrin cell surface interactions*, Platelet activation, signaling and aggregation*, ECM-receptor interaction*, FSH regulation of apoptosis*, Beta-3 integrin cell surface interactions*, Interleukin-4 regulation of apoptosis*, Binding of chemokines to chemokine receptors*, Hemostasis pathway*, Alpha-9 beta-1 integrin pathway*

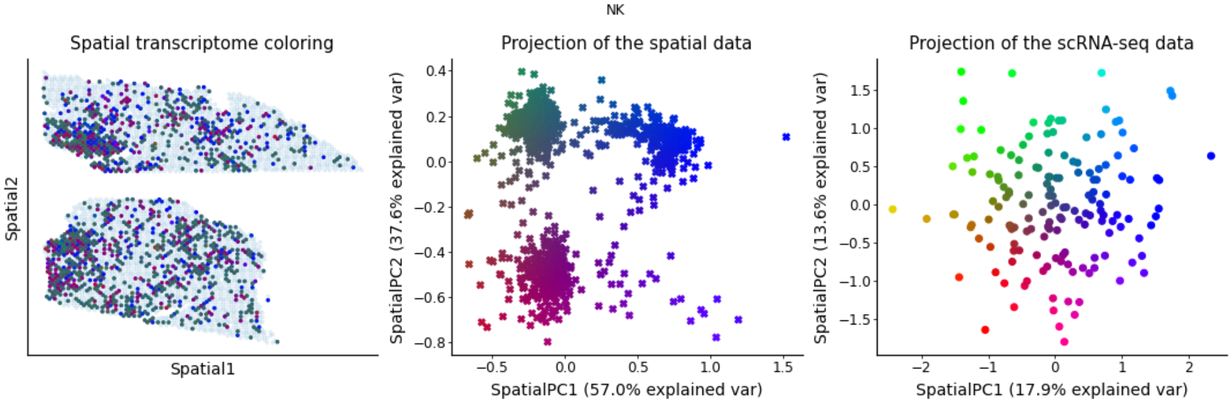


**Genes associated with SpatialPC1**

**Positively**

Zscan20, Ccno, Ksr1, Kyat1, Kynu, Ccr9, Lactb, Lad1, Ccr3, Lamc2, Lancl3, Laptm4b, Ccr10, Ccnf, Car7, Ccne2, Lca5l, Ccne1, Lcn2, Ccnb2, Lhx6, Lhx8, Ccna2, Lins1, Lipg, Lix1l, Kremen2, Kntc1, Kmo, Cd101, Cd209a, Kif11, Kif15, Kif18a, Kif20a, Kif20b, Kif21a, Kif2c, Kif4, Kifc1, Kitl, Klc3, Klhdc8b, Cd200, Klhl15, Klhl23, Klhl32, Klhl35, Klk1, Klra17

Kinesins*, Factors involved in megakaryocyte development and platelet production*, MHC class II antigen presentation*, Cell cycle*, G0 and early G1 pathway*, M phase pathway*, FOXM1 transcription factor network*, Hemostasis pathway*, DNA replication*, p53 activity regulation*

**Negatively**

Gzmc, Syngr1, Mgst3, Gtf2i, Ccl12, Eif3b, Tbc1d7, Axl, Thbs1, Emp1, Fbxo36, Cdh13, Exoc3l4, Eml1, Src, Sgce, Lhfp, Crabp1, Cald1, Prdx4, Nt5dc2, Dut, Mt1, Srm, H1f0, Tcf4, Gja1, Rhoc, Cnn3, Mmp9, Arl2, Apex1, Ranbp1, Tmem176b, Mcm4, Anxa1, Fam216a, Cd81, 9530082P21Rik, Stab1, Mvk, C1qc, Ccl7, Kcnk5, Sdr39u1, Tppp3, Scd2, Mcm5, Desi2, Gmds

Unwinding of DNA, CDK regulation of DNA replication, ID regulation of gene expression, Angiogenesis, Signaling events mediated by PRL, Gap junction trafficking and regulation, Activation of the pre-replicative complex, DNA strand elongation, Syndecan 4 pathway, CXCR4 signaling pathway

**Genes associated with SpatialPC2**

**Positively**

Zscan20, Npy, Nr2f1, Nr2f2, Fam3b, Fam229b, Fam20c, Nrbp2, Nrep, Nrg1, Nrn1, Fam187b, Fam178b, Nrtn, Nsg2, Fam174b, Fam161a, Nthl1, Fam160a1, Fam129c, Fahd1, Nr2c1, Fam71b, Fah, Npl, Nlgn2, Fbxo32, Fbxo2, Fbxl12os, Nnmt, Nnt, Fbn1, Fbln2, Fat1, Fam83h, Fam83g, Fam83e, Fam83d, Fam83a, Nos2, Nos3, Noxo1, Fam71e1, Npdc1, Nuak1, Nudt12, Msantd1, Extl1, Oscp1, Evc2

Nicotinate and nicotinamide metabolism*, Nitric oxide effects, Mechanism of gene regulation by peroxisome proliferators via PPAR-alpha, Ghrelin pathway, Nitric oxide stimulation of guanylate cyclase, Latent infection of Homo sapiens with Mycobacterium tuberculosis, Nuclear receptors, Interleukin-4 regulation of apoptosis, Nuclear receptor transcription pathway, Arginine and proline metabolism

**Negatively**

Ptprcap, AY036118, Atad2, Gm26917, Flt3l, Gm42418, Selenoh, Klra7, Cd7, Ick, S1pr5, Banf1, Dedd2, Asf1b, Ncr1, Ctla2b, Kcnj8, Nkg7, Rpa2, Ppp2r3a, Gm15283, Klhl11, G0s2, Icam2, Cd72, Grap, Herpud1, Tecpr2, Tmem121, Thbs3, Lcmt2, Lyrm7, Krba1, Tab1, Arhgap27os1, Cdyl2, Samd3, Dtymk, Cd9, Gzmk, Ears2, Vamp5, Cct3, Dut, Adh5, Cst7, Slc29a1, Smpd5, Hdc, Elp2

Histidine metabolism, Tyrosine metabolism, Diurnally regulated genes with circadian orthologs, 2-LTR circle formation, APOBEC3G-mediated resistance to HIV-1 infection, Histidine catabolism, Flap intermediate removal from the telomere lagging strand (C-strand), Nucleotide metabolism, Pyrimidine biosynthesis, Homologous DNA pairing and strand exchange

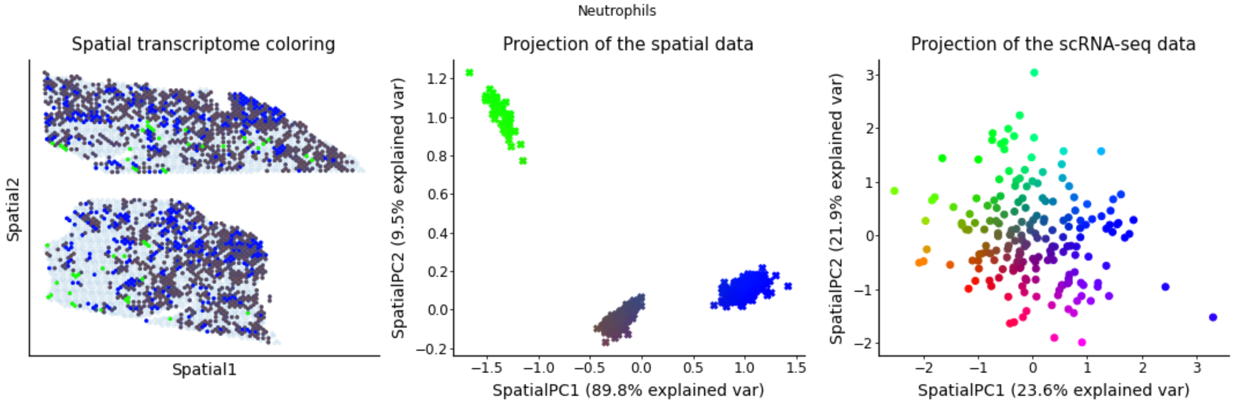

**Genes associated with SpatialPC1**

**Positively**

Zscan20, Loxl4, Cxcl1, Cxadr, Lrig1, Lrp8, Lrr1, Lrrc24, Lrrc32, Lrrc51, Lrrc73, Cx3cl1, Cwc27, Cul7, Cttn, Ltbp2, Ltbp4, Ctsw, Lurap1, Lurap1l, Ctsf, Ly6c1, Ctla4, Cstad, Ly6g5b, Cst6, Csrnp2, Lpar1, Loxl1, Lax1, Lncpint, Lbx1, Lca5l, Lck, Lclat1, Lcmt2, D130020L05Rik, Lcn4, D130017N08Rik, Cysltr2, Lgals7, Cyp7b1, Lhx6, Lhx8, Cyp4f17, Lig1, Cyp2r1, Lins1, Cyp1b1, Cxxc5

Cytochrome P450 pathway, Cytochrome P450 metabolism of endogenous sterols, Phase I of biological oxidations: functionalization of compounds, T cell activation co-stimulatory signal, CTLA4 inhibitory signaling, Adaptive immune system, Biological oxidations, Binding of chemokines to chemokine receptors, Steroid hormone biosynthesis, Tryptophan metabolism

**Negatively**

Ccl6, Thbs1, Hilpda, Bgn, Gm26917, Snrpa1, Ankrd33b, Gata2, Mtmr7, Rrs1, Rab30, Gramd1c, Mmp23, Plxdc2, Reps2, Gng11, Paip2b, Fosl1, Ciart, Rgs1, Rae1, Apoe, Mcpt8, Spp1, Ndrg1, Nedd4, Rit1, Tceal9, Cd63, Slc40a1, Gm42418, Avpi1, Msrb3, Id2, Tppp3, Itgb3, Il7r, Ankrd1, Parva, Tada2a, 9330020H09Rik, Mad2l1, Fgfr1, Timp1, Snhg6, Rcn1, Trmu, Mt1, Cd93, Vwa1

Response to elevated platelet cytosolic calcium, Platelet activation, signaling and aggregation, Beta-3 integrin cell surface interactions, BDNF signaling pathway, HIF-1 transcriptional activity in hypoxia, AP-1 transcription factor network, Osteoclast signaling, Hemostasis pathway, ECM-receptor interaction, Integrin cell surface interactions

**Genes associated with SpatialPC2**

**Positively**

Zscan20, Hspb1, Hoxc9, Hpgd, Hpgds, Hrh1, Bcl11b, Hsd17b7, Hsf2bp, Bcl11a, Hspa12b, Bcdin3d, Bcat1, Bcas1, Bcar1, Hspb6, Hoxb3, Hspb7, Bard1, Hspg2, Htra1, Htra4, Bank1, Bace2, Icam2, Ick, Icos, BC055324, Id3, BC051142, Hoxc10, Hoxa9, BC049352, Hlf, Hist1h3e, Hist1h3g, Hist1h3h, Hist1h4b, Hist1h4c, Hist1h4d, Bend5, Hist2h2ab, Hist2h2ac, Hist2h2bb, Hist3h2ba, Hist4h4, Hk1os, Bdnf, Hoxa7, Bdkrb2

Systemic lupus erythematosus*, Type II interferon signaling (interferon-gamma)*, VEGF, hypoxia, and angiogenesis, Amyloids, RAGE pathway, Integrin cell surface interactions, BRCA1-dependent ubiquitin ligase activity, Valine, leucine and isoleucine biosynthesis, Adhesion and diapedesis of lymphocytes, Stress induction of HSP regulation

**Negatively**

Cxcl2, Ccrl2, Cd14, Ccl3, Isg20, Il1a, Il1rn, Ccl4, Ier3, Rsad2, Ifit1, Usp18, Cdkn1a, Ninj1, Ifit3, Slfn4, Clec4n, Cxcl3, Marcksl1, Ifit3b, Tgm2, Slfn5, Spp1, Ifi208, Irgm1, Thbs1, Il1r2, Ddx27, Chac1, Ankrd50, Chaf1a, Dhfr, Gm14636, 4732465J04Rik, Ddah2, Ifit1bl1, Kbtbd11, Cd70, Pnp, Ifit2, Rnase6, Cdc34, Cd9, Cd74, Gbp2, Sod2, Acod1, Rasip1, E230032D23Rik, Tmem70

TNF-alpha effects on cytokine activity, cell motility, and apoptosis*, Interleukin-1 regulation of extracellular matrix*, Interferon alpha/beta signaling*, Immune system signaling by interferons, interleukins, prolactin, and growth hormones*, Interleukin-4 regulation of apoptosis*, Immune system*, Thymic stromal lymphopoietin (TSLP) pathway*, Interferon signaling*, Cytokine-cytokine receptor interaction*, Binding of chemokines to chemokine receptors*

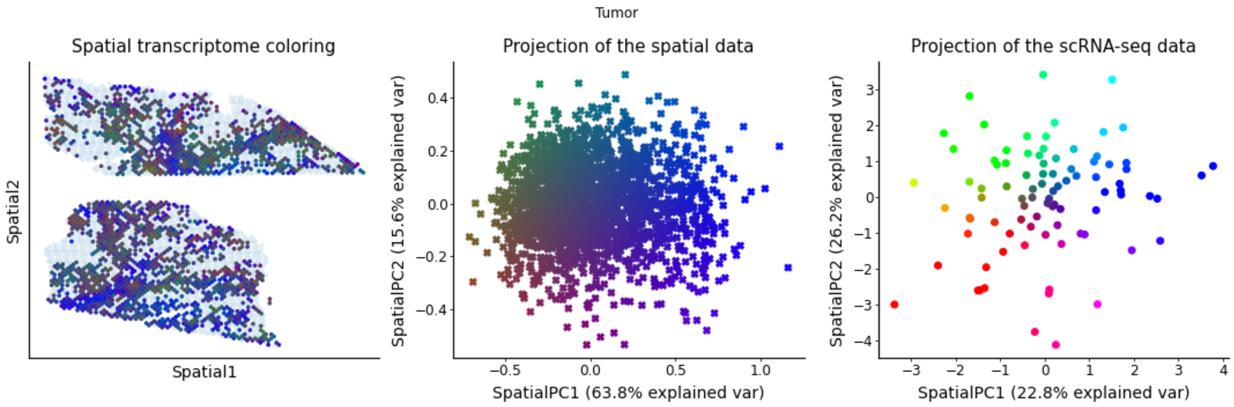


**Genes associated with SpatialPC1**

**Positively**

Zscan20, Gbp2b, Lad1, Gal, Kynu, Krtcap3, Sit1, Siglech, Siglecg, Krt83, Gcsam, Fxyd7, Gdf15, Gemin4, Gfi1, Sh2d2a, Sh2d1b1, Sh2d1a, Sgip1, Ggt1, Plxna4os1, Slc13a3, Gimap1os, Flywch2, Fcna, Slc43a3, Lat, Ffar2, Slc39a2, Flicr, Flt1, Flt4, Slc2a3, Slc15a2, Foxj1, Foxp3, Fpr1, Plxdc2, Plxna4, Slc22a18, Fut7, Fxyd2, Gimap1, Gimap4, Fbxw10, Klrb1b, Gm13203, Gm13391, Gm13546, Gm14010

Transport of glucose and other sugars, bile salts and organic acids, metal ions and amine compounds, Signaling by VEGF, SLC-mediated transmembrane transport, Transmembrane transport of small molecules, VEGF, hypoxia, and angiogenesis, Actions of nitric oxide in the heart, Selenium pathway, Sodium-coupled sulphate, di- and tri-carboxylate transporters, Neurophilin interactions with VEGF and VEGF receptor, Signaling events mediated by VEGFR1 and VEGFR2

**Negatively**

Gstm2, Tbrg1, Anxa1, Cavin3, Apbb2, Usp18, Rcn3, Gnb4, Tcf4, Enpp4, Zbtb20, Prrx1, Tmem176b, Hoxc9, Cyb5r3, Fstl1, H1f0, Hspa4l, Cd9, Col6a1, Sdc2, Rhoc, Aebp1, Igtp, Cand1, Sparc, Plscr2, Gbp5, Nrp2, Mt2, Phf11d, Irgm1, Acat2, Efemp2, Gbp2, Apol9a, Ifit3, Tubg1, Maged1, Gstm1, Mmaa, Fkbp9, AW112010, Iigp1, Cdh13, Ccnd2, Shisa4, Hspa1b, Mvk, Clpp

Interferon alpha/beta signaling, Terpenoid backbone biosynthesis, Interferon signaling, Immune system signaling by interferons, interleukins, prolactin, and growth hormones, Glutathione metabolism, Neurophilin interactions with VEGF and VEGF receptor, TGF-beta regulation of extracellular matrix, Platelet activation, signaling and aggregation, G alpha (12/13) signaling events, Nuclear beta-catenin signaling and target gene transcription regulation

**Genes associated with SpatialPC2**

**Positively**

Zscan20, Gm8013, Gm26935, Gm27042, Gm28347, Gm28935, Gm29156, Sh2d2a, Sh2d1b1, Sh2d1a, Sgip1, Gm29243, Gm29361, Gm3716, Gm37170, Gm42997, Gm44053, Gm44867, Gm5129, Serpine2, Gm568, Siglecg, Siglech, Sit1, Gm17745, Gm16283, Gm16287, Gm16315, Gm16556, Gm1673, Gm16845, Slc22a18, Gm17455, Gm17767, Gm26902, Slc15a2, Slc13a3, Gm19585, Gm20069, Gm20627, Gm20632, Gm26542, Gm26571, Serpinb9b, Gm807, Gm16096, Gm826, Gstp3, Gstt2, Gzmc

Sodium-coupled sulphate, di- and tri-carboxylate transporters, Organic cation transport, Transport of glucose and other sugars, bile salts and organic acids, metal ions and amine compounds, SLC-mediated transmembrane transport, Bile salt and organic anion SLC transporters, Organic cation/anion/zwitterion transport, Oxidative stress, Transmembrane transport of small molecules, Amino acid and oligopeptide SLC transporters, Glutathione metabolism

**Negatively**

Gm42418, AY036118, Gm26917, Spin4, Dkc1, Hist1h1e, Col7a1, 4930581F22Rik, Adh7, Enpp2, Schip1, Armc6, Zfhx4, Cby1, Epb41l4a, Cwc27, Bicd1, Rpusd3, Zfp948, Zeb1, Smc1a, Gzf1, Cdc42bpa, Miga2, Ktn1, Rrs1, Dnajc9, Tmem47, Emp2, Stamos, Fbln2, Anp32b, Cald1, Inpp4b, Huwe1, 1700001O22Rik, Gm11131, Appbp2os, Cdh2, Nol12, F730043M19Rik, Orc1, Morc4, Nexn, Mrps35, Ppid, Plod2, Cenpe, Cdc25a, Rbbp7

E2F-enabled inhibition of pre-replication complex formation, G1/S-specific transcription, Cell cycle, Chromosome maintenance, E2F-mediated regulation of DNA replication, G2/M checkpoints, Mitotic prometaphase, ATM pathway, Collagen biosynthesis and modifying enzymes, G1 to S cell cycle control

### Supplementary Note 3: a spatially-regularized DestVI

#### Adding spot locations as side information in DestVI

While DestVI is specifically designed around spatial transcriptomics data, it does not directly leverage the spatial covariates $\lambda$ to predict cell-type proportions $\pi$. In principle, modeling spatial variations at the spot level with Gaussian processes is feasible. Gaussian processes have been applied in other contexts of spatial transcriptomics analyses (e.g., SpatialDE [75], SVCA [76]), but not for this particular problem. There are two possible explanations for this. The first reason is purely technical: building a Gaussian process model outside of basic regression settings with Gaussian likelihood is hard [58]. The second reason is related to our knowledge of the smoothness of the spatial distribution of cells. Indeed, the idea that cells next to each other should have similar proportions is valid, but a bit simplistic. Indeed, many tissues are highly structured, organized in discrete layers rather than smooth transitions.

We have therefore designed a simple extension of DestVI, in which we enforce smoothness for the cell-type proportions in the tissue. Because, spatial transcriptomics simulation framework explicitly models cell-type proportions as spatially smooth, this data is a natural starting point to understand how to extend DestVI to encourage information sharing between parameters in neighboring spots in a scenario where the influence of spatial covariates is controlled. To do so, we introduce a regularized objective function $L'(\lambda)$,

$$L'(\lambda) := L + \sum_{s,s' \in N(s)} \lambda ||\beta_s - \beta_{s'}||^2,$$

Where $L$ is the original objective function of DestVI, $\lambda$ is a regularization strength hyperparameter and $N(s)$ denotes the set of neighboring spots of $s$ (in the experiments, the 4-nearest neighbors). We note that each contribution $\lambda||\beta_s - \beta_{s'}||^2$ to this penalization shrinks the cell-type abundance $\beta_s$ of every spot $s$ toward the average of its neighbors. Consequently, the penalization aims at encoding a prior assumption that the cell-type proportions in adjacent spots should be related. To be beneficial, the regularization procedure implicitly assumes some knowledge on the spatial structure, represented by $\lambda$. In particular, a too strong regularization could be detrimental in contexts where cell proportions in some areas of the studied tissue are very loosely correlated.

#### Searching for hyperparameters with held-out genes results on simulations

We propose an empirical solution to determine the prior strength $\lambda$ based on held-out data. Although a natural approach to select hyperparameters consists in splitting the spots for spatial cross-validation, this is not applicable here because the parameters influences the cell-type abundance $\beta$, which are spot-specific. As an alternative, we propose to split the genes into a training gene set $G_{train}$ and a testing gene set $G_{test}$. For this splitting to be representative of the different gene modules, we cluster genes based on Hotspot applied to the single-cell data and randomly assign genes to $G_{train}$ and $G_{test}$, such that these sets approximately have equal size.

For a candidate regularization strength $\lambda$, we operate as follows. First, we train DestVI using the regularized objective function $L'(\lambda)$ on the training counts $X_{G_{train}}$. During this step, we learn the generative model parameters (the cell type abundance parameters $\beta$, and the biases $\alpha_{G_{train}}$) as well as the inference network parameters. Second, with $\beta$ fixed, we fit the gene-specific parameters of the model $\alpha_{G_{test}}$ from observations $X_{G_{test}}$. We report the reconstruction error after training, on the set of held-out genes.

#### Results on simulations

We report the held-out reconstruction error, as well as performance for cell-type proportion prediction for DestVI, and the spatially-aware variant for different strength parameters. We noticed that the performance for gene expression imputation is stable (unreported).

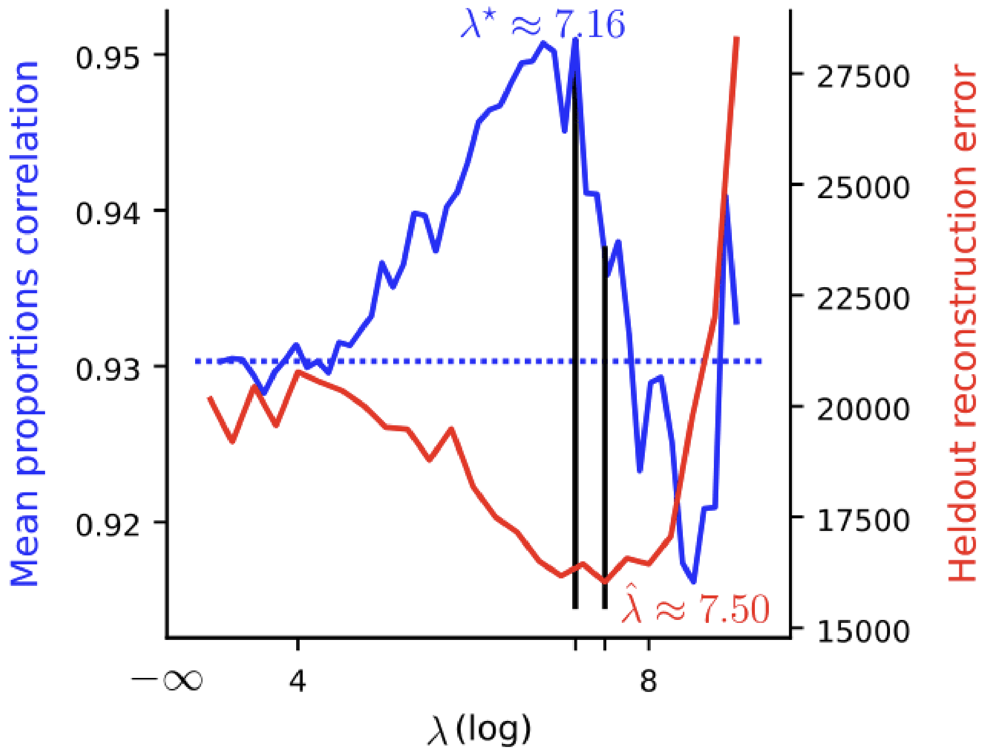


*Figure: Spatial prior strength selection. Held-out reconstruction used for selection (red) with ground-truth average Spearman correlation of proportions with ground-truth (blue). The* $\lambda$ *selection relies on a grid of* 25 *values on a logarithmic scale, ranging from* $10^{3}$ *to* $10^{10}$ *The dashed blue line corresponds to the mean proportion correlation without spatial prior* $(\lambda = 0)$. $\hat{\lambda}$ *and* $\lambda^{*}$ *denote the associated optima.*

We first observe that a broad range of regularization improves proportions prediction over the regular DestVI model (increase of correlation from 0.93 to 0.95), but too high values decrease performance. Our procedure picks a prior strength with an improved correlation over the baseline, and that closely matches the optimal $\lambda$ value. Therefore, our described procedure helps select a prior that improves the proportions prediction.
